# Directional antigenic drift forecasts influenza vaccine effectiveness and guides strain selection

**DOI:** 10.64898/2026.08.27.747648

**Authors:** Omid Arhami, Pejman Rohani

## Abstract

Anticipating antigenic evolution is essential for selecting effective seasonal influenza A/H3N2 vaccine strains. To this end, we integrated data from multiple immunological assays spanning two decades into a unified Bayesian antigenic map. The map resolves twelve antigenic clusters advancing in discrete steps, with several clusters co-circulating in most seasons. In 15 of 21 seasons, the vaccine composition belonged to an earlier cluster than the dominant circulating cluster. We identified geometric features of antigenic space that accounted for three-quarters of the variation in vaccine effectiveness. The direction of each vaccine update relative to recent viral drift predicted effectiveness one season ahead out of sample. In every season, our analyses identified a virus that would have raised predicted effectiveness by 10 percentage points.

---

Influenza A(H3N2) causes more severe disease and greater mortality than other seasonal influenza subtypes (*1*), yet vaccine effectiveness (VE) against outpatient H3N2 illness is among the lowest and most variable: over the past two decades, point estimates in Northern Hemisphere (NH) seasons have ranged from near zero to ∼50% (*2, 3*). This variability can arise from egg adaptation during vaccine development (*3*) but is primarily attributed to antigenic evolution, leading to a mismatch between the vaccine strain and circulating viruses (*4, 5*). Reducing this variability requires rapid pipelines to assess candidate viruses for inclusion in seasonal vaccines. Such pipelines depend on updated antigenic cartography of the circulating viruses, which informs the World Health Organization (WHO) strain-selection process (*6*). However, selecting a strain ahead of next season’s circulation is inherently difficult, as it requires predicting viral evolution (*7–10*).

Using immunological assay data, work over the past three decades has transformed our understanding of how H3N2 evolves antigenically (*4, 11*). Rather than isotropic diffusion through antigenic space, H3N2 follows a predominantly one-dimensional evolutionary trajectory (*8, 12*), punctuated by discrete cluster transitions (*11, 13*). Prior theoretical work on the relation between antigenic distance and VE led to the antigenic distance hypothesis, which holds that VE depends not only on vaccine–virus match but on whether the vaccine update is large enough, and suitably directed, to bypass prior immune memory (*14, 15*). These findings suggest that the *geometry* of antigenic evolution, its directionality and structure, may matter as much as the magnitude of vaccine–virus distance. Here, we link this directional geometry of antigenic escape to a unified, testable model of vaccine effectiveness.

A challenge to the construction of informative antigenic maps for contemporary viruses has been the progressive loss of H3N2 hemagglutination capacity, driven by receptor-binding-site substitutions that reduce binding avidity (*13, 16*). Neuraminidase–mediated agglutination further confounds hemagglutination inhibition (HI) (*16*). This has led to increasing reliance on other immunological assays, especially plaque reduction neutralization (PRNT). Thus, over the past 20 years H3N2 has been characterized antigenically by a mixture of HI and PRNT. To understand H3N2 antigenic evolution and inform vaccine strain selection, we mapped 21,459 HI and 6,607 PRNT titers spanning 2,053 H3N2 viruses (2002–2025) with a Bayesian latent-variable model that treats both assays as noisy observations of a shared underlying antigenic distance, each with its own scale, bias, and noise, ultimately yielding unified antigenic map coordinates (Methods, *Unified HI–PRNT latent antigenic model*; Supplementary Text, *Biological basis for integrating HI and neutralization assays*).

## Twelve antigenic clusters from latent antigenic maps

We constructed an antigenic map from the posterior-mean latent distances obtained from the Bayesian model, with 2,053 unique virus positions in a five-dimensional antigenic space. We then used *k*-means clustering and identified *k* = 12 antigenic clusters over 2002–2025 (Fig. 1; see Supplementary Text, *Choosing the number of antigenic clusters*).

**Figure 1:**
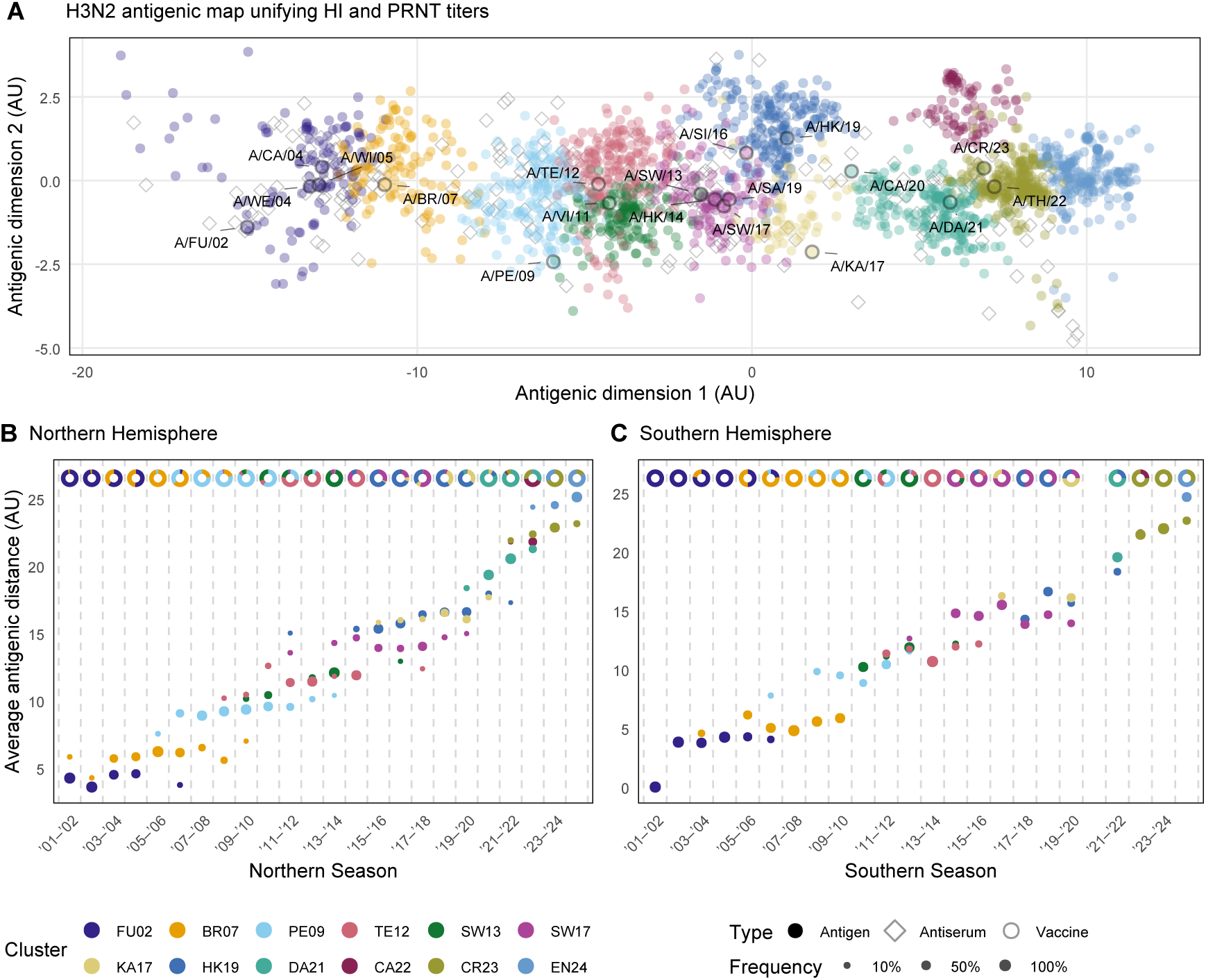
Twelve antigenic clusters of H3N2, punctuated antigenic advance, and cluster cocirculation. (**A**) The 2,053 latent virus positions in five-dimensional antigenic space (projected onto 2-D), colored by the twelve antigenic clusters and named for their representative strains. Clusters were identified by *k*-means clustering (Supplementary Text, *Evidence accumulation clustering of antigenic positions*). Reference antisera (grey diamonds) are shown as map reference points but are not assigned to an antigenic cluster. Labeled points are the recommended vaccine strains. The twelve clusters span 2002–2025, each named for its representative strain, either the vaccine strain whose year is closest to the cluster’s median member year, or the cluster medoid where no vaccine strain is present: FU02 (A/Fujian/411/2002), BR07 (A/Brisbane/10/2007), PE09 (A/Perth/16/2009), TE12 (A/Texas/50/2012), SW13 (A/Switzerland/9715293/2013), SW17 (A/Switzerland/8060/2017), HK19 (A/Hong Kong/2671/2019), KA17 (A/Kansas/14/2017), DA21 (A/Darwin/9/2021), CA22 (A/Castilla-La Mancha/4547/2022, cluster medoid), CR23 (A/Croatia/10136RV/2023), and EN24 (A/England/185/2024, cluster medoid). (**B**) NH series: each bubble is one antigenic cluster in one season plotted at the mean antigenic distance of its strains from the reference strain A/Fujian/411/2002, sized by the fraction of that season’s characterized strains in the cluster (SEE legend), and colored by antigenic cluster. Each ring is divided into the fractions of that season’s characterized strains belonging to each cluster. Antigenic distance accumulates roughly as a staircase, and two or more clusters co-circulate within most seasons. (**C**) Southern Hemisphere (SH) series: same display as (B) for SH surveillance strains. All panels derive from the latent map integrating HI and PRNT data.

## Punctuated antigenic advance and pervasive cluster co-circulation

Consistent with the findings of Smith *et al*. for 1968–2003, we found antigenic advance to be punctuated rather than gradual (*4,11,12*). Measured against a reference strain (A/Fujian/411/2002), the mean antigenic distance of circulating Northern and Southern Hemisphere (SH) viruses rose stepwise: each of the twelve clusters occupied a narrow band of antigenic distance, and most successive clusters were separated by discrete jumps (Fig. 1). Adjacent jumps averaged 1.9 antigenic units (AU) in both hemispheres (1.88 AU Northern, 1.91 AU Southern). One AU corresponds to a two-fold change in HI titer.

The clusters were not temporally exclusive: counting a cluster as present in a season when it exceeded 10% of characterized strains, each cluster remained present for an average of 4.0 seasons in the NH (3.8 in the SH; excluding the three clusters censored at the limits of the record, FU02, CR23, and EN24). In 18 of 24 NH seasons (17 of 22 in the SH), two or more clusters coexisted (Fig. 1; Fig. S2). This pattern of cluster persistence and coexistence contrasts with the greater sequential turnover Smith *et al*. (*4*) reported for 1968–2003, when each antigenic cluster persisted for an average of 3.3 years before being displaced. The contemporary H3N2 population is therefore antigenically fragmented: in most seasons the vaccine confronts not one antigenic target but a mixture of co-circulating clusters present at appreciable frequencies.

Fragmentation degrades vaccine effectiveness directly. Population-level VE is a frequency-weighted average of cluster-specific effectiveness, so a vaccine well-matched to one cluster can still perform poorly when an antigenically distinct cluster co-circulates alongside it (*17*). Summarizing that mixture by a single vaccine-to-mean-virus distance discards its structure: two seasons with the same mean vaccine–virus distance can differ in how widely the circulating viruses are dispersed and whether they form one cluster or several. This motivates a geometric characterization of the antigenic space when several clusters co-circulate.

## Antigenic evolution across the COVID-19 pandemic gap

Annual antigenic change averaged 1.5 AU globally before the pandemic (Fig. S3A). Global influenza circulation collapsed over the two pandemic seasons, 2020–2021 and 2021–2022 (*18–21*). Across those two seasons antigenic advance was 2.8 AU and then 2.5 AU (5.3 AU combined), a per-year rate about 1.75× the pre-pandemic mean. Because our map measures antigenic phenotype directly, this pattern complements reports of accelerated H3N2 genetic and antigenic evolution during the COVID-19 pandemic (*22, 23*). The mean post-pandemic rate rose modestly to 1.8 AU per year, about 1.2× the pre-pandemic mean. Antigenic evolution, therefore, briefly accelerated through the so-called “pandemic gap” (*20, 24*).

Antigenic diversity, by contrast, collapsed under deep transmission suppression: viral population dispersion was at a typical level (2.3 AU) just before the pandemic onset, contracted in 2021–2022 to 1.9 AU, before rebounding (Fig. S3B). The coincidence of lost standing diversity with a large antigenic advance is the signature of a population bottleneck, in which a few antigenically displaced surviving lineages come to define the population, consistent with the documented pandemic contraction of influenza lineage diversity (*18*).

## A recurrent lag between circulating viruses and vaccine composition

Comparison of each season’s circulating strains to that season’s recommended vaccine revealed a persistent lag. While the standing vaccine belonged to the dominant viral cluster in 6 of 21 NH seasons, in every other season it belonged to an earlier cluster than the dominant one (Fig. 2). The same pattern held in the SH, where the vaccine belonged to an earlier cluster in every non-matching season (Fig. 2B; Fig. S4), although sparser SH antigenic characterization precludes a quantitative hemisphere comparison.

**Figure 2:**
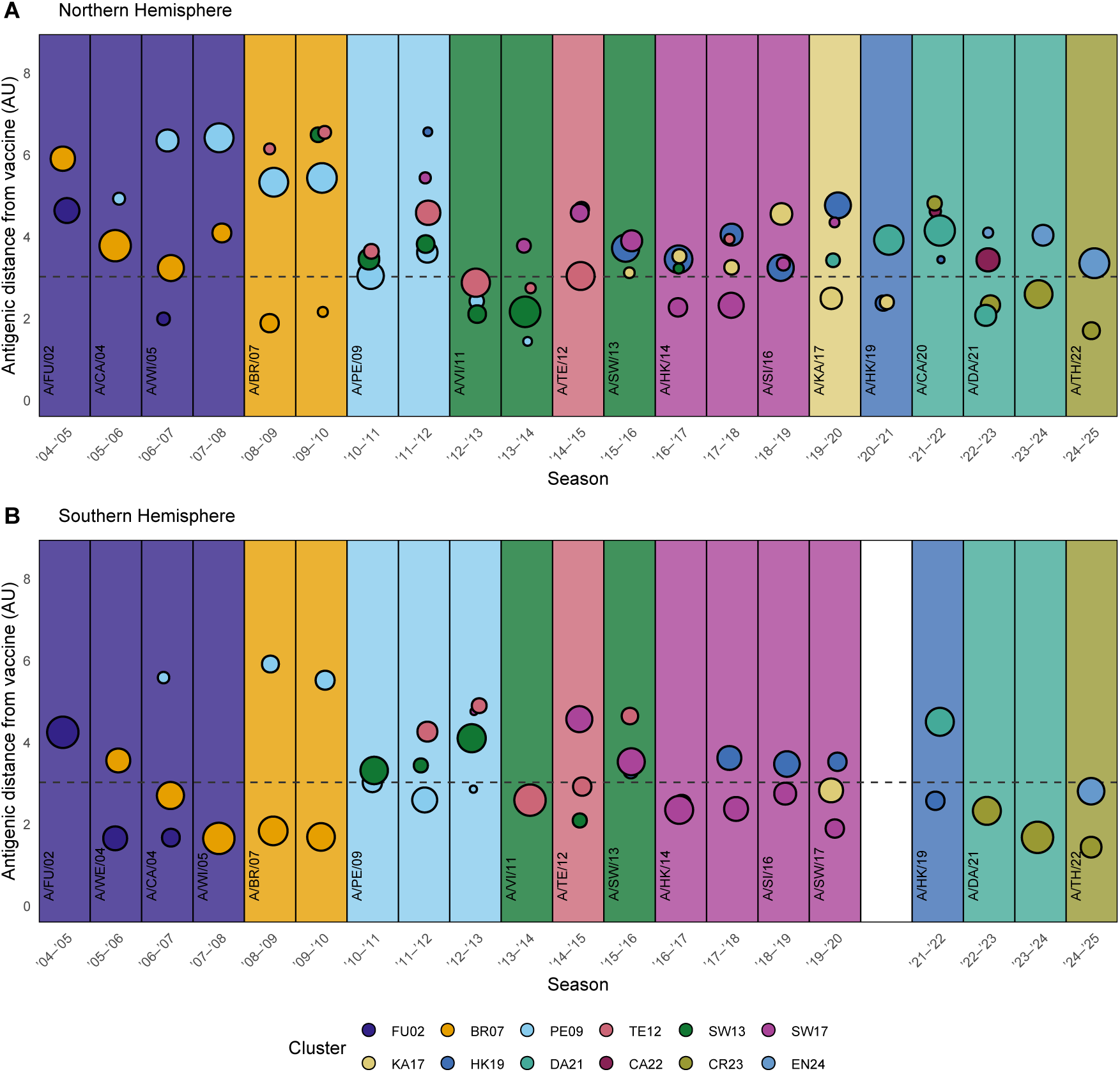
H3N2 vaccine composition lags the circulating population by one or more seasons. (**A**) NH seasons. Bubbles show the mean antigenic distance of each circulating viral cluster from that season’s recommended vaccine strain (vertical axis, AU), sized by the cluster’s frequency and colored by antigenic cluster. Shaded background bands mark vaccine clusters, colored by the cluster of the vaccine strain and labeled with its name; the dashed line marks 3 AU, the conventional antigenic distinction threshold (*25, 26*). In most seasons the dominant circulating clusters (largest bubbles) belong to a more advanced antigenic cluster than the standing vaccine. The vaccine updated into each new cluster only one or more seasons after that cluster rose to dominance. (**B**) SH seasons and WHO SH vaccine recommendations, displayed as in (**A**); 2020–2021 is blank because of no SH strains in our data. The key below the panels gives the cluster colors; the cluster-to-color mapping is identical to Fig. 1. Cluster-aligned trajectories are shown in Fig. S5 (NH), Fig. S4 (SH).

Before updating the vaccine to a new cluster, that cluster’s strains typically circulated 6–8 AU from the prior vaccine for one to three seasons, and the distance from the available vaccine dropped below 3 AU with the update, the conventional antigenic-distinction threshold of an eightfold reduction in HI titer (*25, 26*) (Fig. S5). The lag was sometimes prolonged. The PE09 cluster dominated NH circulation for four consecutive seasons (2007–2008 to 2010–2011) while the vaccine contained viruses from the earlier FU02 and BR07 clusters; the vaccine updated to PE09 in 2010–2011, by which point the TE12 had become the dominant cluster (Fig. 2). The HK19 cluster followed a similar pattern: dominant across most NH seasons from 2015–2016 to 2019–2020, it was first represented in the vaccine for 2020–2021, by which point DA21 had become the dominant cluster (Fig. 1).

Two features of H3N2 dynamics make such a lag difficult to avoid. First, to accommodate vaccine manufacturing constraints, the NH vaccine composition is decided in February, roughly six months before seasonal circulation begins (*6*). A cluster that emerges after that February decision cannot be matched until the following year’s meeting. Second, because cluster transitions are punctuated (Fig. 1), the frontier may advance by a whole antigenic cluster in a single step rather than incrementally, so one selection meeting delay could translate into a large antigenic mismatch. Matching the dominant viral cluster did not, however, guarantee protection. For example, in 2014–2015 (NH), the vaccine belonged to the dominant TE12 cluster, yet the reported VE was ≈9%. Ranking vaccine candidates therefore may be improved by accounting for the geometry of antigenic evolution.

## Directional vaccine updates and strain-cloud geometry as correlates of VE variation

To identify which features of antigenic space are associated with observed VE, we obtained population-weighted VE estimates across 16 NH seasons (2006–2007 through 2024–2025). These were modeled with five theory-derived mechanistic categories (illustrated in Fig. 3), each capturing a distinct route by which antigenic dynamics may shape VE: (i) vaccine–virus match (*4, 5, 27*) (Fig. 3B), which quantifies the antigenic distance between a season’s virus population centroid and the chosen vaccine; (ii) drift geometry (*8, 28, 29*) (Fig. 3C), which measures the aspect ratio of how strongly the circulating strain cloud is elongated along the axis joining the population centroid to the vaccine, relative to isotropic dispersion; (iii) vaccine-update orientation (*14, 15*) (Fig. 3D), which captures both the magnitude of the season-to-season change in the vaccine strain and how closely that change is aligned with the direction of recent viral drift; (iv) viral population structure (*17, 30*) (Fig. 3E), which describes the community composition and diversity of circulating viruses; and (v) vaccination coverage (*31, 32*) (Fig. 3F), which is the population-weighted fraction of individuals vaccinated.

**Figure 3:**
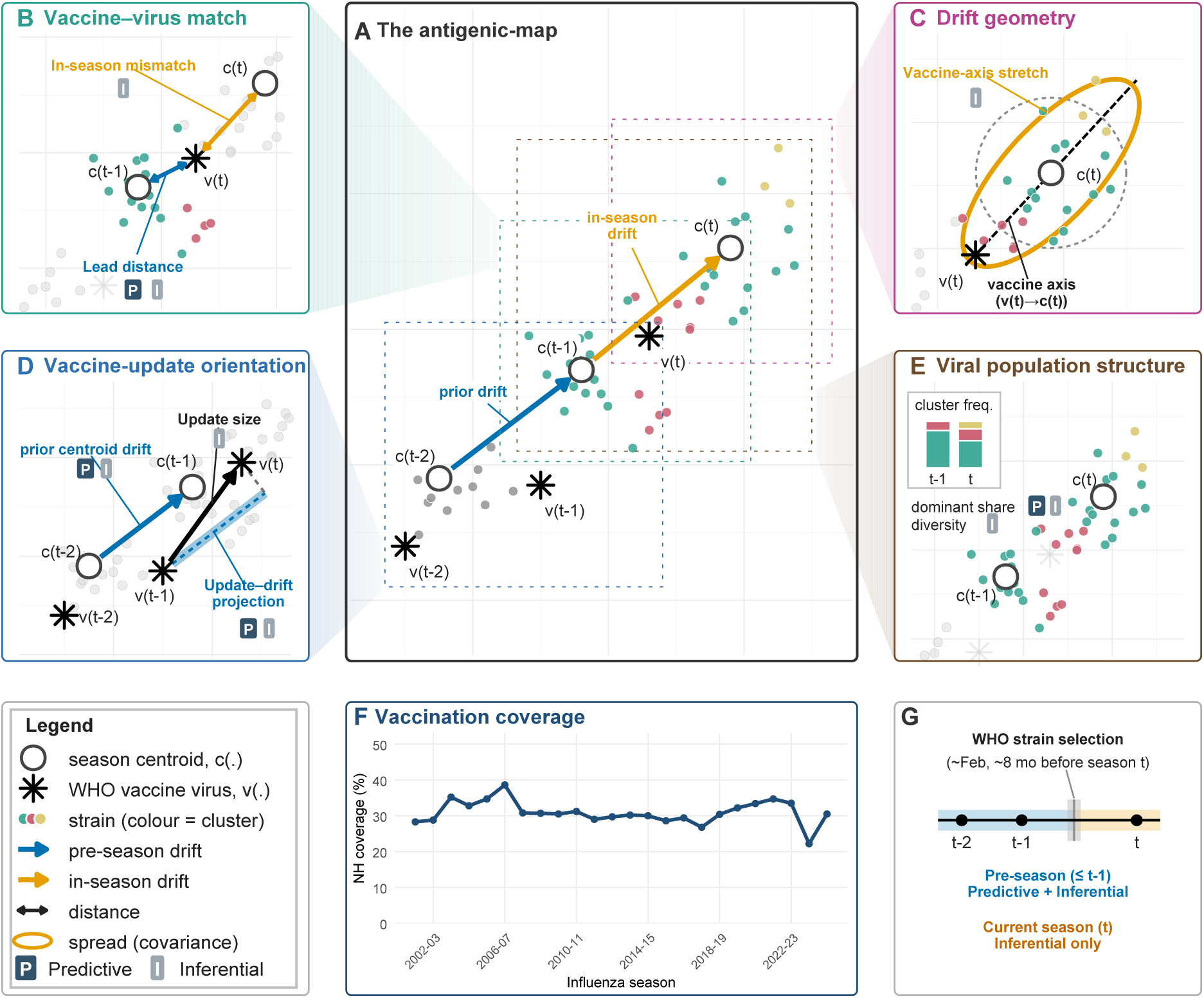
Schematic of the geometric measurements of the vaccine–virus relationship on the antigenic map. Panels **A**–**E** are a schematic: the points, arrows, and ellipses are drawn to define each measurement, not plotted from data. Only **F** and **G** show real values. The five mechanistic categories linking H3N2 antigenic evolution to VE are shown: four are geometric measurements on a common antigenic map (**A**) and the fifth (**F**) is a population covariate. (**A**) Season virus-population centroids (open circles), WHO vaccine picks (stars), and individual strains colored by antigenic cluster, for three consecutive seasons (*t*−2, *t*−1, *t*), with the pre-season drift (blue) and in-season drift (orange). Each faint dashed square on **A** marks the region magnified in one detail panel, linked by a tinted beam. (**B**) Vaccine–virus match: in-season mismatch = |*v(t)* − *c(t)*|; lead distance = |*v(t) − *c (t*−* 1)|. (**C**) Drift geometry: vaccine-axis stretch = variance along the vaccine axis, *v(t)* → *c(t)*, relative to the isotropic reference (ellipse vs. dashed circle). (**D**) Vaccine-update orientation: update size = *v(t) − v(t* − 1)); update–drift projection = projection of the vaccine update (*v*(*t*−1 *v(t)*) onto the prior centroid drift (*c(t)*−2 →*c(t)* − 1)). (**E**) Viral population structure: dominant-cluster share and antigenic diversity (in-season Shannon entropy), read from the cluster-frequency inset. (**F**) Vaccination coverage: NH population-weighted vaccination coverage (%) by season. (**G**) The timeline marks the strain selection time (8 months pre-season): only pre-season information (blue, *t* 1) enters the predictive model; in-season variables (orange, *t*) enter the inferential model only. Pills: P, predictive; I, inferential.

We operationalized these categories with 13 candidate variables: 12 antigenic variables (one to four per category, each a geometric measurement on the antigenic map; Fig. 3; formulae in Methods, *Mechanistic category definitions*) and a pandemic-season indicator as a control. Each variable, with its mechanistic category, temporal window, and hypothesized direction, is listed in Table 1. Published VE estimates derive from the test-negative design (TND), in which the odds ratio (OR) of infection is compared between vaccinated and unvaccinated individuals, and VE = 1 − OR (*33*). We used log odds-ratio (log(OR)) for modeling, which is unbounded, and a more negative value denotes higher protection (Supplementary Text, *Collection and processing of influenza A(H3N2) vaccine effectiveness estimates*).

**Table 1:**
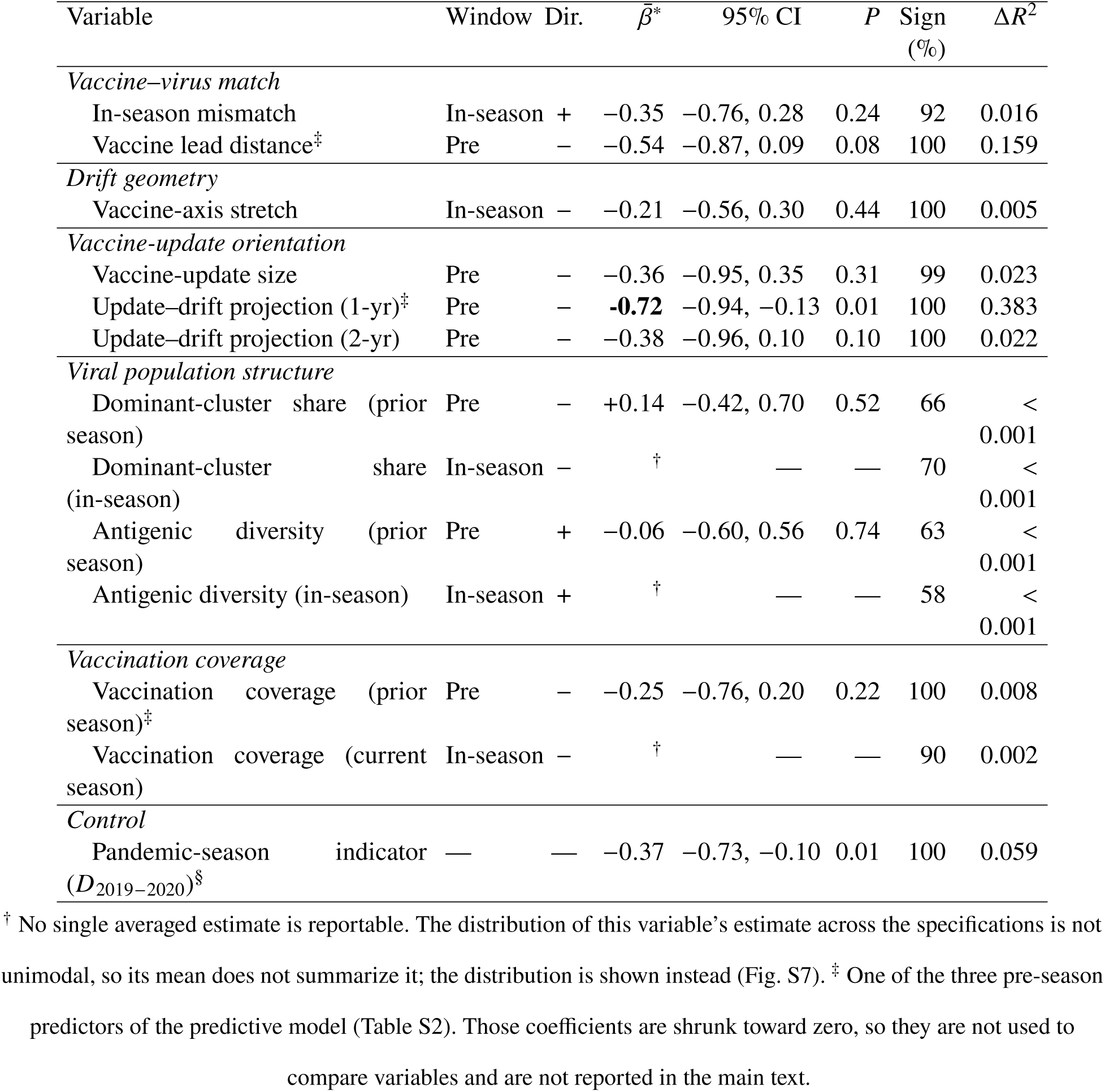
The five mechanistic categories, the 12 antigenic variables that operationalize them, and their estimated associations with. log OR . Each category is a mechanism by which antigenic dynamics may shape VE; its variables measure that mechanism geometrically on the antigenic map (constructions in Fig. 3 and Methods, *Mechanistic category definitions*). “Window” gives when a variable becomes available: pre-season (known before season *t* circulates, hence usable for forecasting) or in-season (concurrent with circulation, usable only in the inferential analysis). “Dir.” is the hypothesized sign of the association with log OR, set before fitting (positive = higher log OR = lower VE). *β̄*^∗^ is the model-averaged partial association with its 95% bootstrap interval. Every specification with at most three predictors drawn from these 13 candidates is fitted, and the estimate is averaged over those containing the variable, so that no estimate is conditional on a selected model (Methods, *Inferential model: multimodel estimation*). *P* is the two-sided bootstrap *P* value, read off the same resamples (Methods, Eq. S12). “Sign” is the percentage of tested specifications in which the estimate keeps the sign shown. Δ*R*^2^ is the variable’s incremental *R*^2^, the part of the fit apportioned to it across the specification space, computed within the three-predictor lattice (Methods, *Variance decomposition by category*). *β̄*^*^ gives the direction and strength of a partial association, Δ*R*^2^ the amount of variance attributable to the variable. At *N* = 16 an interval covering zero marks an association these data cannot resolve, not one shown to be absent (Supplementary Text, *Inferential model: detailed results*).

| Variable | Window | Dir. | $\bar{\beta}^*$ | 95% CI | $P$ | Sign (%) | $\Delta R^2$ |
| --- | --- | --- | --- | --- | --- | --- | --- |
| *Vaccine–virus match* |  |  |  |  |  |  |  |
| In-season mismatch | In-season | + | −0.35 | −0.76, 0.28 | 0.24 | 92 | 0.016 |
| Vaccine lead distance<sup>‡</sup> | Pre | − | −0.54 | −0.87, 0.09 | 0.08 | 100 | 0.159 |
| *Drift geometry* |  |  |  |  |  |  |  |
| Vaccine-axis stretch | In-season | − | −0.21 | −0.56, 0.30 | 0.44 | 100 | 0.005 |
| *Vaccine-update orientation* |  |  |  |  |  |  |  |
| Vaccine-update size | Pre | − | −0.36 | −0.95, 0.35 | 0.31 | 99 | 0.023 |
| Update–drift projection (1-yr)<sup>‡</sup> | Pre | − | **−0.72** | −0.94, −0.13 | 0.01 | 100 | 0.383 |
| Update–drift projection (2-yr) | Pre | − | −0.38 | −0.96, 0.10 | 0.10 | 100 | 0.022 |
| *Viral population structure* |  |  |  |  |  |  |  |
| Dominant-cluster share (prior season) | Pre | − | +0.14 | −0.42, 0.70 | 0.52 | 66 | < 0.001 |
| Dominant-cluster share (in-season) | In-season | − | † | — | — | 70 | < 0.001 |
| Antigenic diversity (prior season) | Pre | + | −0.06 | −0.60, 0.56 | 0.74 | 63 | < 0.001 |
| Antigenic diversity (in-season) | In-season | + | † | — | — | 58 | < 0.001 |
| *Vaccination coverage* |  |  |  |  |  |  |  |
| Vaccination coverage (prior season)<sup>‡</sup> | Pre | − | −0.25 | −0.76, 0.20 | 0.22 | 100 | 0.008 |
| Vaccination coverage (current season) | In-season | − | † | — | — | 90 | 0.002 |
| *Control* |  |  |  |  |  |  |  |
| Pandemic-season indicator ($D_{2019-2020}$)<sup>§</sup> | — | — | −0.37 | −0.73, −0.10 | 0.01 | 100 | 0.059 |

† No single averaged estimate is reportable. The distribution of this variable’s estimate across the specifications is not unimodal, so its mean does not summarize it; the distribution is shown instead (Fig. S7). ‡ One of the three pre-season
predictors of the predictive model (Table S2). Those coefficients are shrunk toward zero, so they are not used to

Because the data include only 16 seasons, selecting a single model among 13 candidates would make each estimated association conditional on an unstable model choice. We therefore estimated each variable’s association with log(OR) by model averaging: we fitted every specification with at most three predictors and averaged each variable’s coefficient over the specifications containing it, weighting by AICc (*34,35*). The reported *β̄*^*^ is the mean partial association with log(OR), standardized by the partial standard deviation (*36*) (Methods, *Inferential model: multimodel estimation*). Uncertainty was estimated by Monte Carlo resampling of the whole procedure (*37*).

Across the 16 seasons, the model-averaged prediction reached *R*^2^ = 0.75 (approximate adj. *R*^2^ = 0.70), *P* = 0.03 against a permutation null (Fig. S9A; Methods, *Permutation calibration*).

Vaccine-update orientation (variable Update–drift projection 1-yr) was the strongest predictor of VE. Averaged over the specification space, it had *β̄*^*^ = −0.72, with a bootstrap *P* = 0.01. It was the only antigenic variable whose interval excluded zero. The same projection at the twoseason horizon carried *β̄*^∗^ = −0.38 (bootstrap *P* = 0.10), and the size of the update, independent of its direction, had a smaller and non-significant association *β̄*^∗^ = −0.36 (bootstrap *P* = 0.31).

Apportioning the explained variance among the mechanistic categories, vaccine-update orientation carried an incremental *R*^2^ of 0.43, against 0.18 for vaccine–virus match, 0.06 for the pandemicseason control, and 0.01 or less for each remaining category (Methods, *Variance decomposition by category*; computed within the three-predictor lattice; table S11). This pattern is consistent with the canalized, largely one-dimensional trajectory of H3N2 antigenic evolution (*8, 28*): a vaccine update aligned with the dominant drift axis better anticipates the direction of subsequent escape (Supplementary Text, *Inferential model: detailed results*).

Scalar antigenic distance carried little signal on its own. A benchmark model containing only the two vaccine–virus distances explained little of the variation (*R*^2^ = 0.20, adj. *R*^2^ = 0.08), indistinguishable from no association at *N* = 16 (*F*_2,13_ = 1.66, *P* = 0.23), against 0.75 for the model-averaged prediction (Fig. S8). The partial associations of both scalar distances with log(OR) increase in magnitude once vaccine-update orientation is controlled for (Table 1; table S12).

Two further associations were consistently negative but small: prior-season vaccination coverage (*β̄*^∗^ = −0.25), which is consistent with a herd-protection mechanism (*31*), and vaccine-axis stretch (*β̄*^∗^ = −0.21; Table 1). Finally, viral population structure category had no variable with a significant association neither at the prior-season nor in-season horizon; Dominant-cluster share and antigenic

diversity were both small and inconsistently signed.

## Forecasting vaccine effectiveness from pre-season antigenic geometry

To evaluate whether antigenic geometry quantified before the start of a season can forecast VE, we fitted a Bayesian ridge regression (*38, 39*) to three pre-season predictors: the update–drift projection, vaccine lead distance, and prior-season vaccination coverage (Table 1). We selected these three before fitting, for the reasons given in Methods (*Predictive model: Bayesian ridge regression*).

The Bayesian formulation propagates three sources of uncertainty into every prediction and into every comparison between candidate strains: the published sampling error of each season’s VE estimate, the noise inherent in antigenic mapping, and the uncertainty in the degree of shrinkage (Methods, *Predictive model: Bayesian ridge regression*; Supplementary Text, *Rationale for a Bayesian predictive model*).

Antigenic geometry fixed at the approximate date of the annual strain-selection meeting (about six months before circulation begins (*6*)) forecasts a substantial fraction of the season-to-season variation in VE. H3N2 escape is canalized along a largely one-dimensional drift axis (*8, 28*), so the pre-season features jointly capture the direction in which the viral population is heading, and effectiveness is partly predictable at selection time. In a leave-one-season-out procedure, the model predicted the observed log(OR) with *R*^2^ = 0.47 (permutation *P* = 0.003; Fig. 4A; Fig. S9B). The 95% predictive intervals contained the observed value in all 16 seasons. Measurement error in the published VE estimates accounts for a large share of the season-to-season scatter the model is scored against (Supplementary Text, *Forecast calibration and skill*).

**Figure 4:**
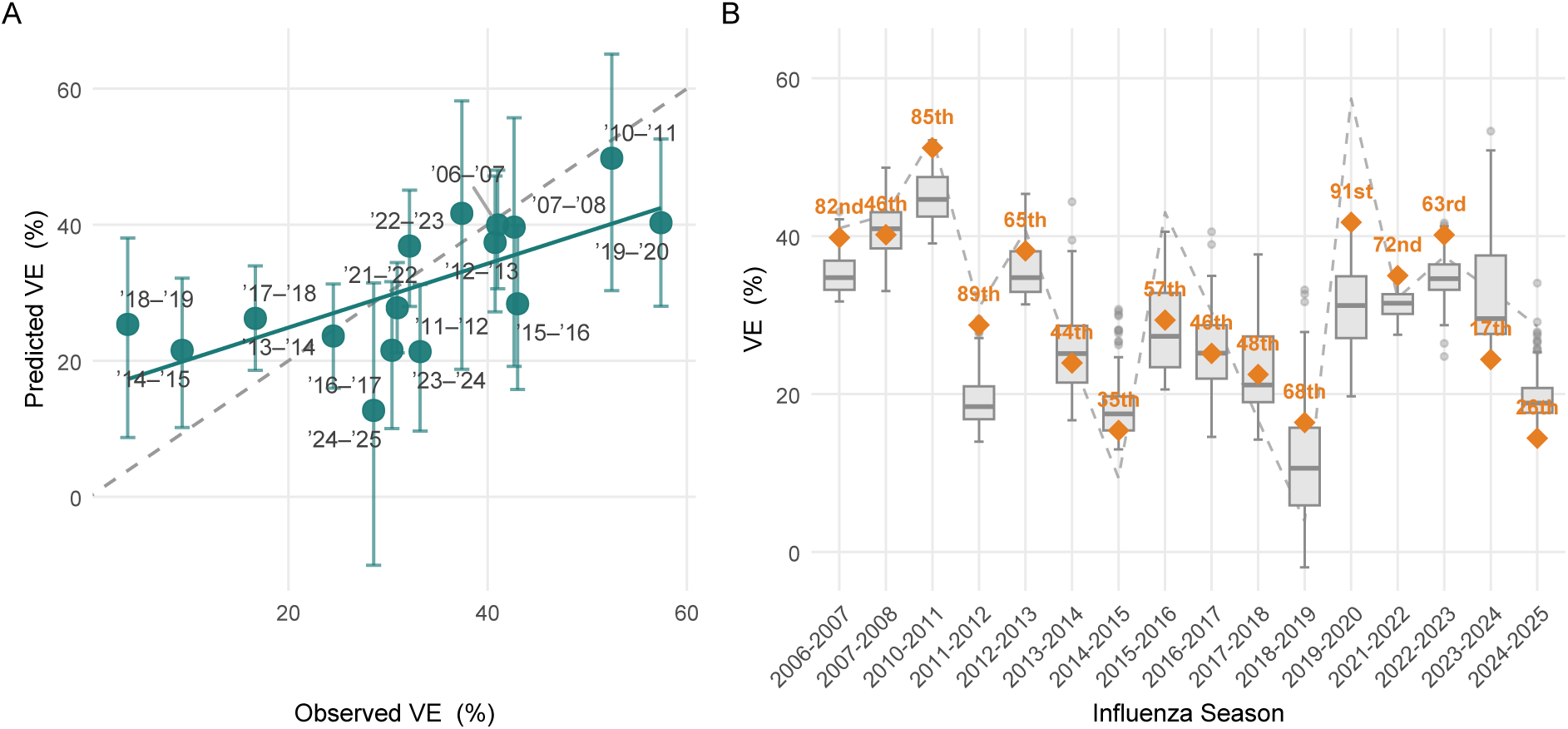
Pre-season antigenic geometry predicts vaccine effectiveness and ranks candidates relative to the WHO selection. (A) Predicted versus observed VE, with each season held out in turn (Pearson *r* = 0.71; *R*^2^ = 0.47; all metrics on the log OR scale); error bars show 95% credible intervals; dashed line indicates perfect prediction. (**B**) Posterior mean predicted VE for each admissible candidate strain, by season (box plots). Candidates assayed by 15 January of the meeting year, two-year look-back. Amber diamonds mark the WHO-selected vaccine’s predicted VE and its posterior mean rank percentile. The dashed grey line shows the observed VE per season; the vertical spread indicates the model’s discriminatory range among candidates.

## Forecast performance under strictly prospective validation

In the results presented so far, we estimated the antigenic map once from the full titer panel. Because no autocorrelation was detected in the outcome series (Fig. S20), seasons can be held out singly. We therefore evaluated the predictive model by holding out one season at a time. Given only 16 VE estimates, this uses the data most efficiently. Because the map was obtained from the complete panel, features at season *t* could in principle depend on titers from later seasons. This dependence only concerns the covariates’ construction, not the outcome; *VE* never enters the antigenic map, so the held-out season’s outcome stays out of model training. Nevertheless, to assess the consequences of this dependence, we refitted the antigenic map (*40*) adding one season at a time. Each added season moved the strains already in the map by a median of 0.5–0.8 antigenic units, less than the impact of antigenic measurement uncertainty alone (0.9–1.0; Fig. S18; table S15), and the predictor values used for the prospective forecasts agreed with their full-data values (*r* = 0.87–0.96; Fig. S19). The same model, restricted to data available before each season, matched the leave-one-season-out performance (expanding-window out-of-sample *R*^2^ = 0.45, permutation *P* = 0.008, over the nine prospectively scored seasons, against LOOCV *R*^2^ = 0.47 over all 16; Supplementary Text, *Antigenic-map stability and leakage audit*).

## Next-season vaccine candidates with higher predicted effectiveness

The WHO Collaborating Centers select strains representative of circulating viruses by integrating antigenic characterization, antigenic cartography, and genetic surveillance (*6*). Our scenario analysis points to one additional source of information this process could draw on: a model that maps antigenic-geometry features to VE and ranks candidate strains by predicted effectiveness, a criterion distinct from antigenic representativeness.

The pre-season predictive model enables a scenario analysis of substitute vaccine candidates. For each of the 16 seasons, we compared the WHO-selected vaccine against every strain serologically characterized before the nomination deadline, that is, every strain whose assay date fell in the two years ending 15 January before that season’s vaccine composition meeting (Methods, *Admissibility of candidates*). For each season *t* we set aside that season’s VE and antigenic features, refitted the model on the other 15 seasons, and used it to predict the effectiveness of every candidate and of the deployed vaccine with uncertainty (Methods, *Scenario analysis of substitute candidates*).

The analysis yields a consistent result: in every one of the 16 seasons, at least one candidate already antigenically characterized by the decision date had higher predicted effectiveness than the WHO-selected vaccine. The mean gain was 10.4 pp (95% credible interval 6.6–14.3 pp). The deployed vaccine nonetheless ranked at or above the median admissible candidate in 9 of 16 seasons (Fig. 4B; Table S5). The gain was not uniform across seasons: it was positive with probability at least 0.95 in 6 seasons, at least 0.90 in 8, and at least 0.50 in all 16.

The per-season gain ranged from 0.16 pp in 2021–2022 to 28.9 pp in 2023–2024, where the selected A/Darwin/9/2021 (predicted VE 24.4%) trailed the model candidate A/Togo/771/2020 (predicted VE 53.3%; gain 95% credible interval 4.1–52.2 pp, *P* = 0.99). In 2024–2025 the selected A/Thailand/8/2022 (predicted VE 14.4%) trailed A/Brandenburg/3/2023 (predicted VE 34.0%) by 19.6 pp (95% credible interval 6.4–34.1 pp, *P* = 0.996). In 2014–2015 the model’s top-ranked candidate was A/Glasgow/407585/2012. It belonged to the HK19 cluster, which rose to dominate the following season, whereas the recommended vaccine’s cluster (TE12) did not persist beyond 2014–2015.

A more conservative rule, which selects the tenth-percentile candidate, still yielded 10.0 pp (95% credible interval 6.8–13.5 pp). It coincided with the top candidate in 11 of 16 seasons and, in 2021–2022, identified the deployed strain itself. Averaged over seasons, the probability of the model’s candidate falling short of the deployed vaccine was 0.19, and by 3.8 pp when it did.

We measured how early each antigenic cluster was nominated for NH vaccine inclusion relative to the season it first dominated NH circulation. The model identified the dominant clusters from four seasons later to two seasons earlier, against five seasons later to one season earlier for the WHO recommendation. On the seven clusters both systems chose, the model’s median lag was one season against three for the WHO. The earlier response did not sacrifice match or specificity. The model’s candidate matched the season’s dominant cluster in 5 of 16 seasons, the same count as the WHO-selected vaccine, and all eight clusters the model ever nominated went on to dominate circulation or enter the vaccine, so it produced no false positives (Table S6).

## Antigenic geometry as a framework for vaccine strain selection

The geometry of H3N2 antigenic evolution, specifically the orientation of each vaccine update relative to recent drift, is associated with seasonal VE and predicts it from pre-season variables, outperforming scalar antigenic distance alone on both counts. The geometric features yield a model-averaged in-sample fit of *R*^2^ = 0.75 across the 16 seasons (permutation *P* = 0.03). The predictive model, using only pre-season variables, recovered 47% of the out-of-sample variance in effectiveness (leave-one-season-out *R*^2^ over all 16 seasons, *P* = 0.003; 45% under a strictly prospective expanding window over nine). In the scenario analysis, a higher-ranked vaccine candidate existed in every one of the 16 seasons, with the gain positive at probability ≥ 0.95 in 6 seasons and ≥ 0.90 in 8.

H3N2 evolves along a dominant antigenic axis, with immune escape concentrated in that direction (*12, 13, 28*). Therefore, choosing the vaccine along the recent drift direction anticipates where the virus will next escape immune pressure. Our results support that logic: the update– drift projection has the strongest partial association with effectiveness among the candidates we examined.

Scalar antigenic distance (vaccine–virus match) carried little signal on its own, despite its established role as a correlate of protection (*5, 41*). Vaccine-axis stretch kept the hypothesized negative sign in every specification we fitted, but the association was small and imprecise (*β̄*^∗^ = −0.21, 95% bootstrap interval −0.56 to 0.30, *P* = 0.44). The variable is also measured in season, so it is subject to reverse causation: a more protective vaccine could exert stronger immune selection, driving escape away from the vaccine and elongating the strain cloud along the vaccine axis (*28,32*). Prior-season vaccination coverage had a positive association with VE (*β̄*^∗^ = −0.25, *P* = 0.22), consistent with the herd-protection mechanism rather than with immune-selection pressure (*31,32*); this result may have been compounded by features of the TND design (collider stratification and coverage-dependent mixing between vaccinated and unvaccinated individuals (*33, 42*)) through which direct VE can understate individual-level protection, more so at lower coverage.

The antigenic features and VE enter the model at the population level, but individual-level VE is also shaped by host factors the model does not include, e.g., immunosenescence (*43*), vaccination and prior-infection history (*44–46*). These host-level sources of variation likely contribute to the residual variation not captured by the inferential model.

The scenario analysis of substitute candidates is the most policy-relevant finding. The WHO-selected vaccine ranked above the median candidate on average (mean 58.5th percentile), yet every season held a candidate with higher predicted VE, by a mean of 10.4 pp (95% credible interval 6.6–14.3 pp). The model’s gain is due to selecting candidates that are farther along the recent drift axis than the WHO pick in 13 of 16 seasons (Fig. 5B). The same preference brought the response to new antigenic clusters forward; the model’s median lag behind first dominance of various clusters was one season, against three for the WHO.

**Figure 5:**
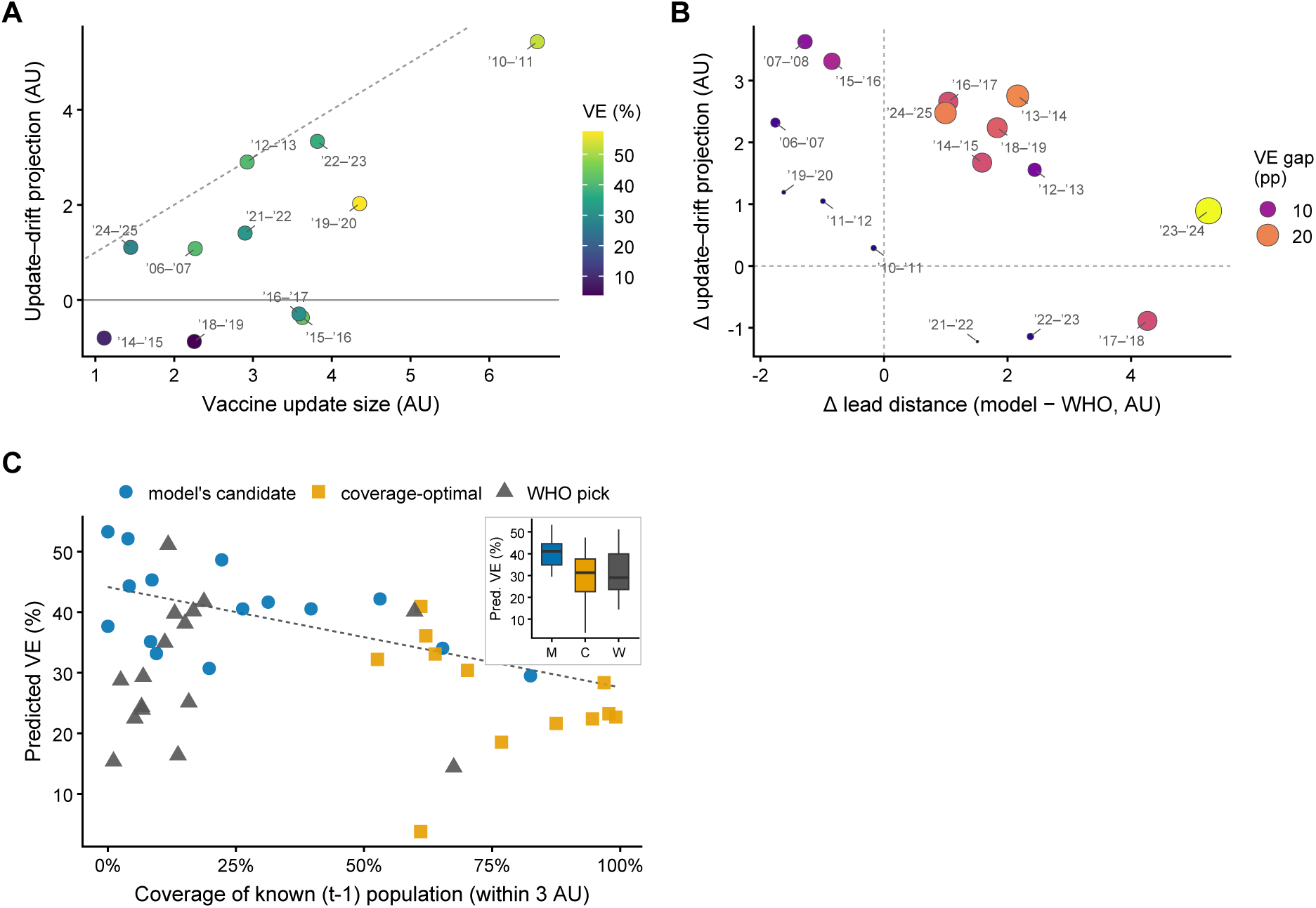
Vaccine strain selection trades population coverage for drift anticipation. (**A**) For each season, the size of the vaccine update (|*v(t)* − *v(t)* − 1)|, AU) versus its signed projection onto the recent drift direction (AU), colored by observed VE (seasons without a VE estimate are omitted); the diagonal marks an update fully aligned with drift and the horizontal line an update orthogonal to it. (**B**) Per season, the model’s candidate minus the WHO pick in lead distance (horizontal) and update–drift projection (vertical; both AU), with the posterior mean predicted-VE gap encoded by point size and color; nearly all seasons lie in the upper region, where the model’s candidate projects farther along recent drift than the WHO pick. (**C**) The coverage–VE trade-off: for the model’s candidate, coverage-optimal, and WHO-selected strains, coverage of the most recently characterized (season *t* 1) viruses (fraction within the 3 AU match radius) versus posterior mean predicted VE; the dashed line is the linear trend for the model’s-candidate and coverage-optimal points; inset, boxplots of the three predicted-VE distributions across seasons. All correlations are computed on the log(OR) scale; *N* = 16 seasons.

A fragmented, fast-moving antigenic landscape constrains any single-strain vaccine. The punctuated advance of H3N2 and the co-circulation of antigenic clusters (Fig. 1) mean that in most seasons no single strain matches the entire circulating population, and that antigenic shifts occur by a whole cluster at once. The recurrent lag between circulation and vaccine composition (Fig. 2) then follows from a roughly six-month selection-to-circulation interval (*6*) colliding with punctuated jumps, and a vaccine chosen to match the current viruses is often overtaken before it is deployed. This reframes the scenario-analysis result: the VE gap between the WHO-selected vaccine and the model’s candidate reflects a structural timing problem rather than misjudgment.

Flexible vaccine platforms that shorten the selection-to-deployment interval (*47*) would narrow the lag directly. Pairing them with a geometry-based ranking that anticipates the next antigenic step, rather than matching the last observed position, could convert part of the effectiveness gap identified here into protection. That conversion is bounded by an intrinsic trade-off in the selection decision: a candidate advanced along the drift to anticipate the coming viruses covers fewer of the viruses already in circulation, so such forward-looking selections carry some risk (Fig. 5C).

H3N2’s declining hemagglutination capacity has left every model built on these assays with a heterogeneous-assay data, mixing HI and PRNT titers (*16, 48*). Our Bayesian latent-variable model addresses this directly: it integrates receptor-binding (HI) and functional-neutralization (PRNT) titers into a single antigenic map, supplying positions that neither single-assay map could provide alone. The shared latent variable hypothesis (*49*) is supported by the model comparison: the fourobservation-process latent model was preferred by ELPD over an independence model and over a correlation model that posits no common latent distance(table S8; table S7). The PRNT observation process is rescaled (*a*_2_ = 0.675, 90% credible interval 0.654–0.697), which the observation model absorbs into its scale and offset. Because the model is robust to missing data, it remains applicable to ongoing surveillance, where assay coverage is incomplete by design.

We note several limitations. First, the analysis spans only 16 seasons with valid VE estimates, a small sample for regression modeling. We mitigated this by averaging over all candidate models rather than selecting a specific model, by regularization, and by held-out validation. In an expanding-window check we refit the antigenic map from prior seasons only and scored ten target seasons (2015–2016 to 2024–2025, nine with VE estimates); errors fell within the held-out distribution and the prospective *R*^2^ was 0.45 (Supplementary Text, *Antigenic-map stability and leakage audit*). Permutation tests that re-ran each procedure on randomly reassigned VE placed both the in-sample fit (*P* = 0.03) and the held-out forecast skill (*P* = 0.003) beyond what these procedures attain when geometry and VE are unassociated. Statistical power nonetheless remains limited: in any individual three-predictor specification, 80% power at *α* = 0.05 requires *f* ^2^ = 0.67 (partial *R*^2^ = 0.40), and power against a moderate effect (*f* ^2^ = 0.15) is very low (0.27) (Supplementary Text, *Inferential model: detailed results*).

Second, the VE outcome is a population-weighted composite of TND estimates (*50*) from three surveillance networks (CDC, SPSN, and I-MOVE). This aggregation smooths over structural heterogeneity between networks while remaining subject to the foundational TND biases arising from healthcare-seeking behavior (*33, 42*) and prior vaccination (*51*).

Third, the strain-substitution analysis does not model feedback of vaccine choice on population immunity or the circulating viral population in later seasons. Ranking candidates within a single season is therefore valid, but the predictions should not be sequentially applied to project multiseason outcomes.

Fourth, the analysis treats admissibility as a question of timing alone. Candidates are limited to isolates characterized early enough to have been weighed at the vaccine composition meeting, but we did not verify that a candidate vaccine virus (CVV) had been derived from each one, nor that it would have satisfied the egg growth-yield, biosafety, and supply constraints that govern manufacture (*6*). Because CVVs are developed 9–12 months ahead of possible inclusion (*52*), the set that was deployable in any season is a subset of the set we ranked. The effectiveness gap we reported is therefore an upper bound on what could have been realized on the egg-based platform. The constraint is weaker for platforms that begin from sequence rather than from a CVV (recombinant HA and mRNA vaccines) (*47*), but it is not eliminated by them: a candidate must still be identified before the composition meeting.

Fifth, our antigenic coordinates derive from a single mapping method (*40*); replication with alternative cartography approaches would strengthen the conclusions.

Sixth, egg-adapted vaccine strains acquire HA mutations that alter antigenicity relative to wildtype viruses (*53, 54*). Our design partially mitigates this: the HI data underlying the map include both egg- and cell-passaged isolates, so the in-season vaccine mismatch measures antigenic distance from the vaccine as characterized in the assay, not from its genomic sequence. In addition, because the map integrates PRNT alongside HI, our results are less dominated by the receptor-binding-site epitopes where egg-adaptation substitutions concentrate (*55*) and to which HI is most sensitive (*13*). However, the mitigation is partial in a specific way: a strain whose egg-propagated manufacturing seed drifted further than the isolate we place on the map would deliver a different VE than its mapped position implies. Our features cannot see that difference, so it enters the unexplained season-to-season variation.

Finally, the VE analysis is restricted to NH seasons. SH antigen sampling is far sparser (median ∼5 versus ∼55 characterized strains per season), no continuous multi-network SH VE series comparable to our composite was available for out-of-sample testing, and no large, consistent SH vaccination-coverage analog exists (Supplementary Text, *Collection and processing of influenza A(H3N2) vaccine effectiveness estimates*). Where SH antigenic dynamics can be characterized, they track the NH series (Fig. 1C).

## Conclusions

Flexible vaccine platforms, including messenger RNA (mRNA) and recombinant technologies, are compressing the timeline between strain selection and manufacturing (*47*), creating an opportunity to base selection decisions on more current antigenic surveillance data. Quantitative antigenic geometry could support automated, real-time strain ranking within the WHO Collaborating Center pipeline, complementing expert virological judgment with a reproducible, data-driven metric. A natural extension is to combine the antigen-side geometric model presented here with host-side immunological correlates of protection, such as pre-vaccination antibody landscapes (*29*) and systems-level immune signatures (*56*), to explain the residual VE variance attributable to host heterogeneity. Our results demonstrate that the directional structure of H3N2 antigenic evolution, not just the magnitude of vaccine–virus distance, is associated with seasonal VE and predicts it, and that exploiting this structure could substantially improve influenza vaccine strain selection.

## Data and statistical framework

Antigenic characterization data were drawn from WHO Collaborating Centre reports for 2002– 2025: 104,183 HI titers on 12,374 viruses and 12,022 PRNT titers on 2,304 viruses. Map fitting is superlinear in the number of strains, so we selected a panel rather than mapping the full record. Viruses were prioritized by the number of distinct antisera against which they returned a titer within the assay’s detection range. A titer at the detection limit bounds an antigenic distance without locating a strain, and a strain measured against fewer than *d* + 1 antisera is not uniquely placed in a *d*-dimensional map. Vaccine strains and viruses characterized by both assays were retained irrespective of rank; the dual-assay viruses are the only ones that carry information linking the two assays. A per-year floor preserved every year of the record. The retained panel comprises 21,459 HI and 6,607 PRNT titers on 2,053 viruses and 269 antisera, of which 1,107 viruses were measured by both assays, contributing 4,851 of the 5,308 available dual-assay pairs. Selection used titer data only and no vaccine-effectiveness outcome. Full rules and before-and-after statistics are given in Supplementary Text, *Serological assay data preparation*.

The latent antigenic distances were estimated via a Bayesian latent variable model (the four-observation-process latent-distance model) implemented in Stan (*57*). For each virus–serum pair, four data sources (HI titer, PRNT titer, and Euclidean distances from independent HI and PRNT antigenic maps) constrain a shared latent distance *θ_i_* ∼ Uniform(0, *L*). Observation equations link *θ_i_* to each source through assay-specific scaling (*a _j_*), intercept (*b _j_*), and noise (*σ_j_*), with the HI titer anchoring the latent scale (*a*_1_ = 1, *b*_1_ = 0). Censored titers enter via the normal cumulative distribution function (CDF) or survival function. We placed a uniform prior on *θ_i_* to avoid imposing a population distribution or shrinkage on the latent positions, which are themselves the inferential targets; the global calibration parameters instead received weakly informative priors that regularize estimation and aid identifiability (*57, 58*): *a _j_* ∼ Normal(1, 1.5) truncated to *a _j_* > 0 (centered on the equal-scale hypothesis), *b _j_* ∼ Normal(3, 5), and a half-Cauchy scale prior *σ_j_* ∼ Cauchy(0, 2.5) with *σ_j_* > 0. Because *N* = 23,215, the posterior is likelihood-dominated. Model validation relied on posterior predictive checks as the primary criterion (table S7). We compared five candidate models for the latent structure. The four-observation-process latent-distance model was preferred by expected log pointwise predictive density (ELPD) over an independence model, whereas the three remaining alternatives (titers only, free four-observation-process correlation without a latent distance, and within-assay correlated errors added to the latent model) failed to converge (table S8). Parameter estimates were stable across four prior specifications (weakly informative, informative, vague, and uniform) and under random missingness and new virus introduction (Methods, *Unified HI–PRNT latent antigenic model*). The Topolow algorithm (*40*) then mapped these latent distances into a five-dimensional antigenic space. *k*-means clustering of the extracted virus positions identified twelve antigenic clusters (Supplementary Text, *Evidence accumulation clustering of antigenic positions*).

We treat the antigenic map as a measurement instrument estimated once from the full panel, not a feature recomputed per season. The estimation runs two independent single-assay maps, unifies them through the Bayesian latent model, and maps the latent distances into five-dimensional coordinates. We verify empirically (Supplementary Text, *Antigenic-map stability and leakage audit*) that adding each successive season of titers moves the strains already in the map less than antigenic measurement uncertainty does, and that the predictor values used for prospective forecasts closely track their full-data values.

Antigenic dynamics were summarized descriptively from the same coordinates. For each season we computed the mean antigenic distance of circulating NH viruses from a fixed reference strain (A/Fujian/411/2002) and from that season’s WHO-recommended vaccine, together with the frequency of each antigenic cluster; co-circulation was defined as two or more clusters each exceeding 10% of characterized strains. Full procedures, cluster-aligned distance trajectories, and the concordant SH replication are given in Methods, *Antigenic dynamics and vaccine–distance analyses*, and Supplementary Text, *Antigenic dynamics and the vaccine–circulation lag*.

We organized 12 antigenic variables into five mechanistic categories, each operationalized by one to four variables: vaccine–virus match, vaccine-update orientation, viral population structure, drift geometry, and vaccination coverage (Table 1), and included a pandemic-season indicator as a control, for 13 variables in total. The vaccination-coverage covariate is a NH population-weighted composite of United States (CDC FluVaxView), European (Eurostat/ECDC), and Canadian (Statistics Canada) coverage; its sources and cross-region harmonization are detailed in Supplementary Text, *Collection and processing of vaccination coverage data*. Variables span two temporal windows: pre-season and in-season. Pre-season variables include both measures derived from *t*−1 circulating strains and vaccine-update metrics; because the WHO fixes the season-*t* composition at the February meeting, vaccine (*t*) is an observable well before season *t* begins.

Two complementary VE models were fitted. The inferential model selects no specification: it fits all 377 models of at most three predictors drawn from the 13 pre-specified candidates and averages each coefficient over the 79 models containing its predictor, weighted by AICc (*34, 35*), with intervals and *P* values from bootstrapping the entire procedure (*37, 59*). The predictive model used Bayesian ridge regression (*38, 39*) with measurement-error terms for both the published VE estimates and the antigenic predictors, fitted by Hamiltonian Monte Carlo (*60*), on three preseason predictors. Scenario analysis of substitute candidates used per-season refitted models to draw predicted VE for all candidate strains per season (Methods, *Scenario analysis of substitute candidates*). The geometry underlying the strain-selection trade-off (per-candidate coverage of the recently characterized population and drift-relative placement) is described in Methods, *Strainselection trade-off and candidate coverage*.

## Acknowledgments

We gratefully acknowledge John Drake, Derek J. Smith, Christian Gunning, Toby Brett, and Maria A. Gutierrez for valuable discussions and insights.

## Funding

This project has been funded with Federal funds from the National Institute of Allergy and Infectious Diseases, National Institutes of Health, Department of Health and Human Services, under Contract No. 75N93021C00018 (NIAID Centers of Excellence for Influenza Research and Response, CEIRR).

## Author contributions

O.A.: Conceptualization, Methodology, Software, Formal Analysis, Data Curation, Visualization, Writing Original Draft. P.R.: Conceptualization, Supervision, Investigation, Funding Acquisition, Writing Original Draft & Editing.

## Competing interests

The authors declare no competing interests.

## Data, code and materials availability

The HI and PRNT titers underlying the antigenic maps were compiled from vaccine-composition meeting reports published by the Worldwide Influenza Centre at the Francis Crick Institute. The raw titers may be requested directly from The Francis Crick Institute. All derived antigenic-map coordinates, model code, and analysis code are available at (*61*). Data S1 holds the source data for every figure and table in this paper. VE estimates were compiled from the published reports of the CDC US Flu VE Network, I-MOVE/VEBIS, and the Canadian SPSN, cited individually by network and season in Supplementary Text, *Collection and processing of influenza A(H3N2) vaccine effectiveness estimates*. No physical materials were generated in this work.

## Materials and Methods

All analyses were implemented in R and Stan. Code development was assisted by Claude Code (Anthropic; various model versions, 2025–2026); methodology was designed by the authors, and all AI-assisted code was reviewed by the authors before use.

### Vaccine effectiveness data

Seasonal H3N2-specific vaccine effectiveness (VE) estimates were compiled from three test-negative design (TND) surveillance networks (*50, 62*): CDC Flu VE Network (United States, population ∼330 million), I-MOVE/VEBIS (European Union member states, ∼450 million), and the Sentinel Physician Surveillance Network (SPSN; Canada, ∼38 million). For each network– season combination, a single adjusted, all-ages, H3N2-specific, end-of-season outpatient TND VE point estimate and its 95% confidence interval (CI) were extracted from the published literature. For each season, we computed a population-weighted average VE across available networks and converted to log(OR) for linear modeling (*33*). The three networks’ season-level estimates were concordant, with between-season variation dominating between-network variation (Fig. S11). Pooling subtype-specific VE across surveillance locations follows established meta-analytic practice (*2*); the rationale, concordance statistics, and limitations are detailed in Supplementary Text, *Basis for pooling across networks*. Seasons with no H3N2-specific VE from any network (2005–2006, 2008– 2009, 2009–2010, and 2020–2021) were excluded, yielding *N* = 18 seasons with valid VE estimates (2004–2005 through 2025–2026; table S9). After listwise deletion of seasons missing antigenic predictor data, the effective analysis sample was *N* = 16. Full details of source networks, selection criteria, aggregation procedure, and limitations are provided in Supplementary Text, *Collection and processing of influenza A(H3N2) vaccine effectiveness estimates*.

### Antigenic cartography

We obtained HI and PRNT titers from the WHO Collaborating Centre for Reference and Research on Influenza (Crick Worldwide Influenza Centre, London), with a subset of HI titers accessed through their published digitization (*63*). Contemporary H3N2 antigenic characterization is a mixture of the two assays because the progressive loss of hemagglutination has forced growing reliance on neutralization (Supplementary Text, *Biological basis for integrating HI and neutralization assays*). Of the *N* = 23,215 virus–serum pairs in the retained panel (2,053 viruses, 269 reference sera; 2002– 2025), most carry a measurement from only one assay: HI titer for 92.4% of pairs, PRNT titer for 28.5%; only 20.9% carry both. Rather than mapping one pooled titer table, we produced antigenic coordinates in three passes of the Topolow algorithm (*40*), which jointly estimates strain and serum positions in a low-dimensional antigenic space via topological optimization and accommodates the sparse, incompletely observed distance matrices characteristic of serological panels.

#### Pass 1: independent single-assay maps

We mapped the HI titer panel and the PRNT titer panel separately, tuning each map’s algorithm hyperparameters and dimensionality independently on its full (not common-subset) panel. This yielded two self-consistent antigenic maps, one per assay.

#### Pass 2: four-observation-process measurement set

For every virus–serum pair we recorded its Euclidean distance in each of the two single-assay maps and paired these with the two raw titers, giving four measurements of the same antigenic separation—the HI titer, the PRNT titer, and the HI- and PRNT-map distances. The Bayesian latent-variable model below reconciled these four observation processes, each with its own scale, offset, and noise, into one shared latent antigenic distance *θ_i_* per pair (*Unified HI–PRNT latent antigenic model*).

#### Pass 3: final latent map

We ran Topolow a final time on the matrix of posterior-mean latent distances to map all strains into a single five-dimensional antigenic space. This final latent map—not either single-assay map—supplied the coordinates for every downstream antigenic variable. One antigenic unit (AU) corresponds to a two-fold change in titer (*4*).

Using the two single-assay maps as inputs, rather than mapping the pooled titers once, places the HI and PRNT evidence on a common scale before the final mapping: each map contributes a network-consensus distance constrained by every titration involving the two strains, and the latent model resolves their differing scale and noise rather than averaging incompatible readouts. The identifiability role and surplus information of the two map distances are established in the Supplementary Text (*Map distances as independently informative indicators of the latent distance*).

### Unified HI–PRNT latent antigenic model

Each virus–serum pair *i* (*i* = 1, …, *N*; *N* = 23,215) carried up to four measurements of the same antigenic separation, assembled in the cartography step (Pass 2): the HI titer, the PRNT titer, and the Euclidean distances from the independent HI and PRNT Topolow maps, all expressed on the base2 logarithmic antigenic-distance scale (AU) defined above. We modeled the four measurements *y*_1_*_i_*, …, *y*_4_*_i_* as noisy readings of a single latent antigenic distance *θ_i_* shared by the pair,

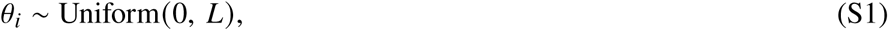

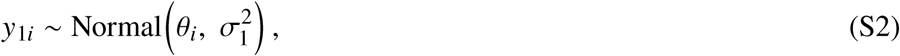

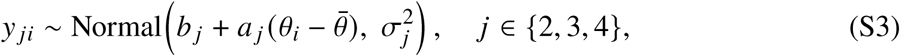

where *a _j_* > 0 is an assay-specific scaling, *b _j_* an offset, *σ_j_* the measurement noise, and *θ̄* the mean HI titer distance (centering decorrelates each slope from its offset). The HI titer observation process anchors the latent scale by construction (*a*_1_ = 1, *b*_1_ = 0), so *θ_i_* is expressed in HI antigenic units and the other three observation processes are calibrated to it; each offset *b _j_* is then the expected reading of assay *j* for a pair at the mean antigenic distance, and each slope *a _j_* describes how that assay rescales true antigenic distance.

We placed a uniform prior on every *θ_i_* (upper bound *L* = 16 AU, well above the largest observed distances) so that the latent positions—themselves the inferential targets—are shaped by the data rather than by an imposed population distribution or shrinkage. The global calibration parameters received weakly informative priors that regularize estimation and aid identifiability (*57, 58*),

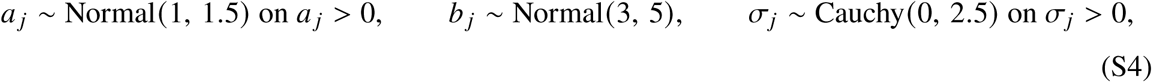

centering the scaling priors on the equal-scale hypothesis (*a _j_* = 1) and giving the noise parameters a heavy-tailed half-Cauchy scale. Because *N* = 23,215, the posterior is dominated by the likelihood. HI and PRNT titrations are reported to the nearest two-fold dilution and are bounded by each assay’s detection limits, so the two titer observation processes (*j* ∈ {1, 2}) enter the likelihood as censored observations, whereas the two map distances (*j* ∈ {3, 4}) are continuous networkconsensus estimates and enter as exact. Writing *φ* and Φ for the standard normal density and cumulative distribution function and *μ _ji_* for the observation-process mean in Eqs. (S2)–(S3), a titer observation process contributes

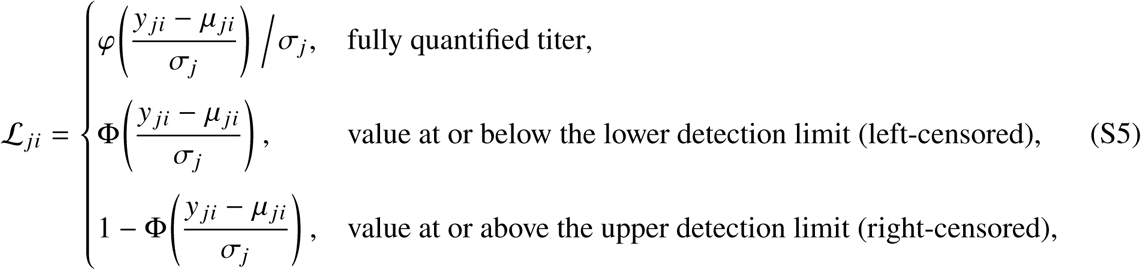

so a censored titration contributes the probability that the latent distance is consistent with the reported detection limit rather than a point density. The posterior-mean latent distances *θ̂_i_* for all pairs were then assembled into a pairwise distance matrix and mapped by Topolow (Pass 3 above) into the five-dimensional antigenic space from which all downstream variables were computed.

To confirm that a shared latent distance with conditionally independent observation-process errors best describes the four observation processes, we compared five models by their expected log pointwise predictive density (ELPD), estimated by Pareto-smoothed importance-sampling leaveone-out cross-validation on the pair-level log-likelihood (*64*). Besides the four-observation-process latent model above, we fitted: (i) a no-latent independence model in which each observation process has its own mean and variance with no shared structure, *y _ji_* ∼ Normal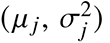; (ii) a twoobservation-process latent model using only the HI and PRNT titers, which is under-identified—the two titers supply three second moments (two variances and one covariance) for four unknowns—and failed to converge (*R̂* = 2.25) (*65*); (iii) a no-latent correlation model placing a saturated four-variate normal on the observation processes, (*y*_1_*_i_*, …, *y*_4_*_i_*)^⊤^ ∼ Normal_4_(***μ***, **Σ**) with **Σ** = diag(***σ***) **R** diag(***σ***) and **R** ∼ LKJ(2), which captures correlation across observation processes without positing a common antigenic distance, and failed to converge (*R̂* = 4.47); and (iv) a correlated-error extension of the four-observation-process latent model that adds within-assay residual correlations—between the HI titer and HI-map observation processes (*ρ*_HI_) and between the PRNT titer and PRNT-map observation processes (*ρ*_PRNT_)—conditional on the shared latent distance, so that it nests the four-observation-process model at *ρ*_HI_ = *ρ*_PRNT_ = 0 and departs from it only if the paired observation processes share error beyond that distance; this model failed to converge (*R̂* = 1.074).

The four-observation-process latent model was preferred by ELPD (table S8), converged well (*R̂* ≤ 1.003), and recovered all four data sources in posterior predictive checks (table S7). The two map distances were necessary for identifiability and additionally carry cross-strain information absent from the single pairwise titers (*Map distances as independently informative indicators of the latent distance*).

We confirmed the model’s robustness in three additional analyses. A sensitivity analysis across four prior specifications (informative, weakly informative, vague, and uniform) yielded convergent models with stable parameter estimates in all cases. A random-missingness test (M MISS: 40% of observations removed at random) recovered the scaling and offset parameters within 2.3% relative change and the noise parameters within 8% (largest for the low-noise HI observation processes, *σ*_3_ = 7.4%), confirming robustness to incomplete data. A virus-level missingness simulation (M NEW: 50 viruses with HI removed, 50 with PRNT removed) showed negligible bias for HI-only recovery (mean absolute error 0.016 log_2_ units) but substantial uncertainty for PRNT-only recovery (mean absolute error 1.51 log_2_ units), reflecting the intrinsically noisier PRNT observation process.

### Antigenic dynamics and vaccine–distance analyses

To characterize the temporal structure of the antigenic map we used the posterior-mean latent positions of the 2,053 virus (antigen) strains and their twelve cluster assignments. For the cumulative-distance analysis (Fig. 1) we fixed A/Fujian/411/2002 as a common reference and, for each season and cluster, computed the mean Euclidean distance of the cluster’s strains from the reference in the five-dimensional space and the cluster’s share of that season’s strains. For the vaccine-distance analysis (Fig. 2) each strain’s Euclidean distance was measured to the coordinates of its season’s WHO-recommended H3N2 vaccine strain, and seasons were grouped into vaccine clusters (maximal runs of consecutive seasons sharing one vaccine). The cluster-aligned view (Fig. S5) re-centered every vaccine cluster at its first season so that distances before, during, and after each cluster could be compared on a common relative-time axis. Co-circulation was quantified as the number of antigenic clusters each accounting for ≥10% of a season’s characterized strains. Analyses were run separately for the NH and SH, with vaccine recommendations matched to hemisphere; the main text reports the NH series (Fig. 1B) and the SH series is concordant (Fig. 1C and Fig. 2B). These analyses are descriptive: distances and cluster shares depend on the estimated map and cluster assignments, and the vaccine-distance comparison uses vaccine strains as characterized in the assay rather than as specified by their genomic sequence (egg- versus cell-passage caveats as in the main-text Limitations). Because the cluster assignment of an individual vaccine strain can shift between map re-estimations, we report the lag as a population-level regularity in the distance distributions rather than as an exact per-season cluster-match count.

### Mechanistic category definitions

Guided by the literature, we specified five mechanistic categories, each acting through a distinct mechanism by which antigenic dynamics may shape VE, and operationalized them with 12 antigenic variables, whose geometric construction is shown schematically in the main-text Fig. 3 and whose temporal group, hypothesized direction, and modeling status are listed in Table 1; a pandemic-season indicator (*D*_2019−2020_) was included as a 13th, control variable. Throughout, *v_t_* denotes the vaccine-strain coordinates for season *t* and *c_t_* the centroid (mean coordinates) of the *n_t_* strains circulating in season *t*, both points in the *D* = 5-dimensional antigenic space; *x_i_*_,*t*_ denotes the coordinates of strain *i* and ‖ · ‖ the Euclidean (ℓ_2_) norm.

*Vaccine–virus match* (Fig. 3B). Cross-reactivity decays with antigenic distance (*4, 5*) (see (*66*) for the theoretical framework). We therefore hypothesized that a greater vaccine-to-population antigenic distance is associated with lower VE (larger log(OR)). We measured the Euclidean distance from the vaccine strain to the season centroid in 5D antigenic space (mismatch inseason = ‖*v_t_* − *c_t_* ‖). We additionally measured the distance between vaccine(*t*) and the prior-season strain centroid (vaccine lead distance = ‖*v_t_* − *c_t_*_−1_ ‖). Because vaccine(*t*) is selected from *t* − 1 surveillance, a larger value indicates a deliberate forward pick that leads the previous season’s population along the drift axis rather than tracking it. Given the directional, canalized trajectory of H3N2 evolution (*8, 28*), we hypothesized that a greater lead distance is associated with higher VE (more negative log(OR)), opposite in sign to the same-season mismatch. These per-season vaccine–circulation distances are shown in the main-text Fig. 2 and, re-centered on each vaccine cluster, in Fig. S5.

*Vaccine-update orientation* (Fig. 3D). We hypothesized that larger vaccine updates and updates better aligned with recent viral drift are associated with higher VE (more negative log(OR)). We measured vaccine-update size (update size = ‖*v_t_* − *v_t_*_−1_ ‖) and the projection of the vaccine-update vector onto the prior viral-drift direction. With the unit drift vector *d̂ _t_*_−1_ = (*c_t_*_−1_ − *c_t_*_−2_)/‖ *c_t_*_−1_ −*c_t_*_−2_ ‖, the two temporal windows are the one- and two-season update vectors projected onto it, update drift projection = (*v_t_* − *v_t_*_−1_) · *d̂ _t_*_−1_ and update drift projection 2y = (*v_t_* − *v_t_*_−2_) · *d̂ _t_*_−1_; the signed projection is positive when the update tracks recent drift and negative when it opposes it. The consecutive vaccine-strain distances underlying update size are shown in Fig. S12.

*Viral population structure* (Fig. 3E). Population-level VE is a frequency-weighted average of clade-specific effectivenesses, so a vaccine matching one co-circulating cluster can still perform poorly when an antigenically distinct cluster circulates alongside it (*17, 30*). We therefore hypothesized that a higher dominant-cluster share is associated with higher VE (more negative log(OR)), whereas greater antigenic diversity is associated with lower VE (more positive log(OR)). Writing *p_k_*_,*t*_ for the frequency of antigenic cluster *k* in season *t*, we measured dominant-cluster share (dominant share t = max*_k_ p_k_*_,*t*_) and antigenic diversity (antigenic diversity t = Shannon entropy Σ *_k_ p_k_*_,*t*_ ln *p_k_*_,*t*_). The antigenic-cluster frequency series *p_k_*_,*t*_ from which these two measures are computed are shown in Fig. S2, and their cumulative-distance summary in the main-text Fig. 1.

*Drift geometry* (Fig. 3C). H3N2 evolution is largely confined to a dominant antigenic dimension (*8*), so the strain cloud in a given season is typically elongated rather than isotropic. Because cross-reactive protection extends only over a finite antigenic radius around the vaccine (*29*), the orientation of this elongation relative to the vaccine—not just the mean vaccine–virus distance— governs how many circulating viruses fall within range of that cross-reactive protection. We hypothesized that greater concentration of strain variance along the vaccine axis is associated with higher VE (more negative log(OR)), by keeping a larger fraction of circulating viruses within that range. We measured the vaccine-axis stretch, the concentration of strain variance along the vaccine axis *c_t_* → *v_t_* relative to an isotropic reference. With unit vaccine axis *û _t_* = (*v_t_* − *c_t_*)/‖ *v_t_* − *c_t_* ‖ and centered-strain projections *a_i_*_,*t*_ = (*x_i_*_,*t*_ − *c_t_*) · *û_t_*, the along-axis variance is 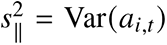 and the per-axis variance expected under isotropic dispersion is 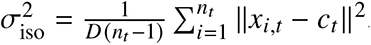 giving 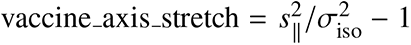 when strain variance concentrates along the vaccine axis beyond the isotropic expectation, = 0 under isotropy). The dominant-axis elongation motivating this mechanistic category is visible in the seasonal antigenic map (Fig. S13).

*Vaccination coverage* (Fig. 3F). Prior-season vaccination coverage modulates VE through competing mechanisms: herd protection (*31*), which predicts higher VE, versus immune-selection pressure (*32*), which predicts lower VE. Because the two mechanisms imply opposite signs, we treated the direction of this mechanistic category as tested rather than assumed a priori. We included both current-season (vaccination coverage t) and prior-season (vaccination coverage tm1) coverage rates; both rates, with their hypothesized signs, are listed in Table 1. The construction of these coverage rates—source systems, cross-region age harmonization, and population weighting—is detailed under *Collection and processing of vaccination coverage data*.

Variables span two temporal windows: pre-season (data available before season *t* circulation begins) and in-season (concurrent with circulation). All variables involving vaccine (*t*)—including vaccine-update distance and drift-projection metrics—are classified as pre-season because the annual NH vaccine composition meeting occurs in February, roughly six months before the autumn onset; vaccine (*t*) is therefore selected and publicly known before season *t* begins.

### Inferential model: multimodel estimation

#### Estimand

The inferential analysis estimates, for each candidate variable *j*, its partial association with log(OR) accounting for other variables and not conditional on which model was fitted. A selection procedure reports a different quantity: the coefficient of *j* in the single specification that survived elimination. At *N* = 16 with 13 candidates, the elimination step itself carries large sampling variance, and the coefficients it yields are conditional on that selection, so their nominal *p*-values do not support confirmatory inference (*67*). Averaging over a fixed specification space defines the estimand independently of the selection outcome. This approach extends the extreme-bounds method of characterizing how an estimate varies across a specification space (*68, 69*). Leamer and Leonard summarized that variation by the interval between the smallest and largest coefficient in the space; we follow Sala-i-Martin, who weighted each specification by its fit rather than retaining only the extremes, and report a weighted mean and its spread (*70*).

#### Specification space

The candidate pool is the 12 antigenic variables of the five mechanistic categories plus the pandemic-season control, fixed by theory before any model was fitted. For a specification *m* we fit by ordinary least squares

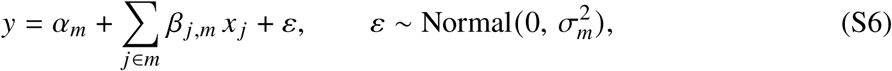

with *k_m_* ∈ {1, 2, 3} predictors and df*_m_* = *n* − *k_m_* − 1 ≥ 12 residual degrees of freedom; the inversevariance weighted sensitivity arm below uses the same equation with weights 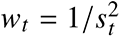. Applying the pre-specified complexity cap of three predictors gives

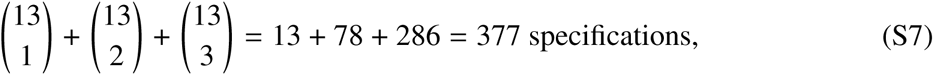

and each variable appears in exactly 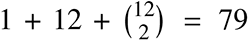 of them. Both counts are fixed by combinatorics, so neither the size of the space nor a variable’s representation in it is a modelling choice.

#### Weights and averaging

Each specification is scored by the small-sample corrected Akaike criterion. With *K* the number of estimated parameters—the *k_m_* slopes, the intercept and *σ*—and RSS*_m_* its residual sum of squares,

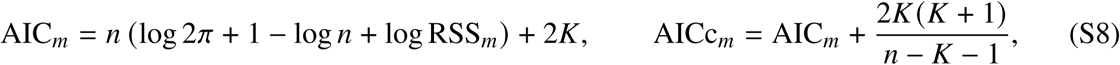

counting *σ* as a parameter in both the penalty and the correction; this convention is applied identically wherever AICc is reported in this work. Specification *m* then receives the Akaike weight *w_m_* = exp(−ΔAICc*_m_*/2)/*_m_*′ exp(−ΔAICc*Σ_m_* /2) (*34*). The reported coefficient for variable *j* is the average over the specifications that contain it, with weights renormalized within that set,

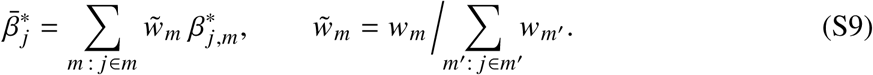

Averaging over containing specifications, rather than setting *β_j_* to zero where *j* is absent, keeps the estimand a partial association. The zero-filled alternative averages over the whole space with the coefficient set to zero wherever *j* is absent,

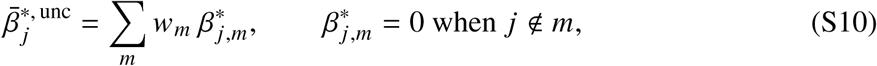

with the weights normalized over the full space; it confounds effect size with how often a variable appears (*34*) and is reported alongside for completeness. Estimates under uniform weights are also reported, because estimated weights carry their own uncertainty and equal weighting is often competitive (*71*).

#### Standardization

Under collinearity the scale in the denominator of a partial regression coefficient changes from specification to specification, so the conventional standardization *β̂ s_x_*/*s_y_* is not commensurate across the models being averaged and averaging it is not meaningful (*36*). We therefore standardize by the partial standard deviation,

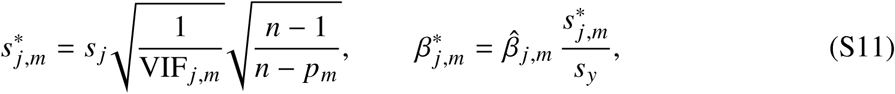

where VIF*_j_*_,*m*_ is the variance inflation factor of *j* in specification *m* and *p_m_* its number of parameters. The partial standard deviation is that of Cade (*36*); we divide by *s_y_* in addition, so that *β*^∗^ is expressed in standard-deviation units of the outcome and its magnitude is directly readable against the conventionally standardized coefficients of the predictive model. Because *s_y_* is the same in every specification, this rescaling leaves unaffected the commensurability across specifications that motivates the partial standard deviation. The averaged quantity *β̄*^*^*_j_* of Eq. S9 is what the inferential results report, and is written *β̄*^∗^ throughout. The predictive model, being a single fixed specification, uses the conventional standardization instead, written *β*_std_ (Eq. S25).

Commensurate scaling is necessary but not sufficient for averaging to be sensible: Cade names a unimodal distribution of the estimates and a valid interpretation of the individual parameter as further requisite conditions (*36*). The interpretability condition is not trivial, because a coefficient need not carry the same interpretation in every model in which it appears (*72*). We treat unimodality as a condition to be checked rather than assumed, and report no averaged estimate for a variable whose weighted mean and median disagree in sign or whose sign consistency falls below 60%; for those variables the distribution itself is reported (Fig. S7). Interpretability is addressed by construction: the pool is a pre-specified set of geometric measurements on one map, so a coefficient carries the same meaning in every specification containing it, and the sensitivity analysis restricting each mechanistic category to a single indicator tests whether that assumption is load-bearing.

#### Uncertainty

Two sources are separated. Specification uncertainty is the spread of 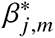 across the 79 containing specifications, summarized by its median, its 5th and 95th percentiles, and the fraction sharing the reported sign, and displayed as a specification curve (*73*) (Fig. S7). Sampling uncertainty is obtained by bootstrapping the entire procedure: for *b* = 1, …, *B* with *B* = 1,000, we resample the 16 seasons with replacement, re-enumerate all 377 specifications on the resample, refit each, recompute the AICc weights and re-average, giving 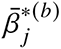. The variance of model weighting therefore enters the interval, which an analytic Buckland-type standard error conditions away (*35, 37*). The reported interval is the 2.5th to 97.5th percentile of 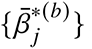, with the bias-corrected and accelerated interval (*59*) reported alongside. A variable can be signed consistently across specifications and still have an interval covering zero; the two statements are compatible, and the interval is the wider one.

#### Significance

We report one *P* value per variable, read off the same resamples as the interval.

Let 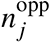 be the number of resamples whose averaged estimate opposes the sign of 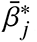, and *B_j_* the number in which *j* is estimable, so that

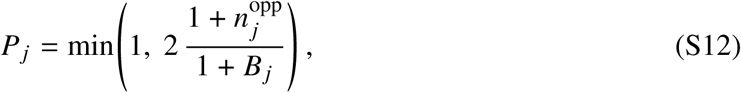

where adding one to numerator and denominator keeps any *P* value from being reported as exactly zero. This is the achieved significance level of the percentile interval: *P_j_* < *α* exactly when the 1 − *α* interval excludes zero, so the interval and the *P* value are two readings of one quantity and cannot disagree. It is therefore a *P* value obtained by inverting an interval, not an exact test against a null-sampling distribution, and we read it as a measure of how stable the estimate is under resampling of the seasons. Bootstrapping an estimator that follows model selection is inconsistent, because selection makes the estimate discontinuous in the data (*67*). Akaike weights are continuous in the residual sum of squares, so the averaged estimate is a continuous function of the data and that objection does not transfer. Sign consistency across specifications is reported alongside as a robustness diagnostic, not as evidence of an association; it is high for most candidates and therefore cannot rank them.

#### Permutation calibration

Two quantities lack a null reference: the per-variable association, whose bootstrap *P* value measures stability, and the in-sample *R*^2^ of the averaged prediction, which has no degrees of freedom from which to derive one. We calibrated both against a permutation null (*73*). For *r* = 1, …, *R* with *R* = 2,000, the 16 values of log(OR) were randomly reassigned among seasons with the predictor matrix held fixed, and the entire procedure—all 377 fits, the AICc weights and the averaging—was repeated. Under the null hypothesis that log(OR) is unassociated with every candidate, the seasons are exchangeable, and the replicates sample the null distribution of any statistic the procedure produces. For a statistic *T* with observed value *t*,

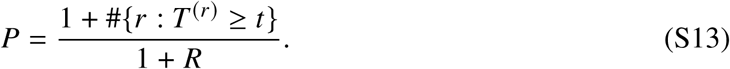

We applied Eq. S13 to the *R*^2^ of the averaged prediction, to the largest *R*^2^ in the space, and to | *β̄*^*^_*j*_ | for each variable. To account for all 13 candidates, we also set 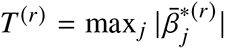, the largest averaged association in replicate *r*, and compared it with each variable’s observed | *β̄*^*^*_j_* |. Taking the maximum within each replicate retains the correlation among candidates, which a Bonferroni correction ignores. Exchangeability requires the absence of serial dependence in log(OR), which the tests reported in Supplementary Text (*Antigenic-map stability and leakage audit*) did not detect. Because a reassignment removes every association at once, the per-variable *P* values are exact only under the null that no candidate is associated; the maximum-based value addresses the claim that the largest of the 13 associations exceeds chance.

#### Variance decomposition by category

Within specification *m*, variable *j*’s incremental *R*^2^ is its contribution to 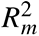 averaged over all *k_m_*! orders in which the predictors could enter (*74, 75*); these contributions sum to 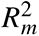 exactly. These shares were then averaged across specifications with the model weights and summed within mechanistic category. The full averaging-over-orderings decomposition requires the saturated 13-predictor model, which has 2 residual degrees of freedom at *N* = 16; we therefore compute it within the three-predictor lattice. We describe the result as an order-averaged incremental *R*^2^ restricted to that lattice rather than as the full decomposition.

#### Conditional contrasts

To ask what controlling for one mechanistic category does to another, we split the specifications containing variable *j* by whether they also contain an indicator of category *c* and averaged within each half. The difference between the two averages is the part of *j*’s estimate attributable to confounding with *c*. This quantity has no counterpart in a single fitted model, which can only report the estimate conditional on the covariates it happens to include.

#### Diagnostics and sensitivity

Classical assumption diagnostics are reported as a distribution over the specification space rather than for one favored fit, and the averaging is repeated on the subset satisfying them. Further sensitivity analyses re-run the averaging under uniform weights, under inverse-variance weighted least squares using the published VE confidence intervals, restricted to specifications carrying at most one indicator per mechanistic category, and omitting each season in turn. The last of these is the influence audit appropriate to this estimand: leaving out one season and re-running the whole procedure asks which observations move the reported quantity, whereas Cook’s distance asks only which observations move one fit.

#### Predictive model: Bayesian ridge regression

We fitted the predictive model on three pre-season predictors drawn from the 13 theory-defined candidates. In the inferential model, two mechanistic categories carry the great majority of the incremental *R*^2^: vaccine-update orientation at 0.428 and vaccine–virus match at 0.176, against 0.059 or less for each remaining category (Table S11). We took the leading variable of each category. The update–drift projection (incremental *R*^2^ = 0.383) is the component of the vaccine update that lies along the recent drift direction, and it indexes how well the update tracks the axis along which the virus population escapes. Vaccine lead distance (incremental *R*^2^ = 0.159) is the antigenic distance from the new vaccine to the strain population circulating when the strain is chosen, and it measures how far ahead of that population the vaccine is placed. Both are fixed at the February composition meeting and read from the antigenic map before the season begins, so the model can be scored prospectively. We added prior-season vaccination coverage as the only non-antigenic predictor: its own incremental *R*^2^ is small (0.008), but prior-season population immunity shifts realized effectiveness in a way antigenic geometry cannot express (*31*). These three predictors leave four estimated parameters including the intercept, and 12 residual degrees of freedom at *N* = 16. A pool this small is also what prospective evaluation demands: the earliest fold of the expanding window trains on seven seasons (Supplementary Text, *Antigenic-map stability and leakage audit*), so a larger pool would approach one parameter per observation in the early folds.

Let *y_t_* denote the published composite log(OR) for season *t* and *s_t_* its standard error, obtained by mapping the published 95% confidence interval through log(OR) = log(1 − VE/100), which is monotone, and dividing the resulting half-width by 1.96. Let *z_t_* collect the three predictors, standardized to zero mean and unit variance on the seasons used for fitting, so that each coefficient is expressed in log(OR) per predictor standard deviation and does not depend on the units of the antigenic map. Both inputs to the regression are estimated rather than observed. The outcome is a published VE estimate that carries a reported confidence interval, and each predictor is a function of the latent antigenic distances *θ* (Eq. S1) rather than a directly measured quantity. The model carries a known-variance measurement layer for each input. The outcome layer separates what is measured from what is predicted:

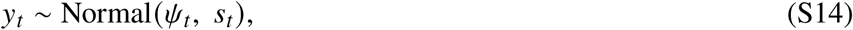

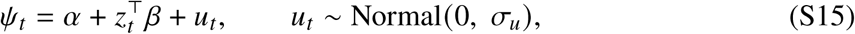

where *ψ_t_* is the true but unobserved effectiveness of season *t*, *α* the expected log(OR) of a season with average predictor values, *β* the three coefficients, and *u_t_* the part of true effectiveness that antigenic geometry does not explain. The season-level latent quantity is written *ψ* throughout to keep it distinct from the latent antigenic distance *θ* of Eq. S1, which belongs to the measurement model upstream of the map. Equation S14 is a measurement model with a *known* variance, the standard device for combining estimates that carry their own reported uncertainty (*76–78*). Integrating *u_t_* out gives the outcome-side marginal likelihood,

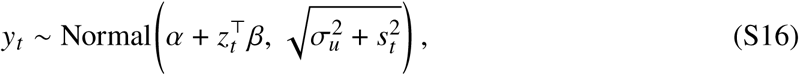

which makes *σ_u_* the season-to-season variability *in excess of* the uncertainty already present in the published VE estimates rather than a mixture of the two.

We placed a normal prior on the coefficients, *β_k_* ∼ Normal(0, *τ*). This is the Bayesian counterpart of ridge regression: the posterior mode under this prior is the ridge estimate, with the ridge penalty *λ* and the prior scale *τ* related by *λ* = *σ*^2^/(*nτ*^2^) (*38, 39, 79*). Rather than fix *τ*, we gave it a half-normal hyperprior, *τ* ∼ Normal^+^ (0, 0.10), a scale of one tenth of a log(OR) unit per predictor standard deviation, so the degree of shrinkage is learned from the data around a defensible centre rather than asserted. We used *σ_u_* ∼ Normal^+^ (0, 0.10) and *α* ∼ Normal(*ȳ*, 0.5), weakly informative choices of the kind recommended for variance parameters in small hierarchical models (*80, 81*). Sensitivity to all three prior scales is reported in Table S4.

The predictor layer applies the same known-variance device to each *z_t_*. This error has no closed-form expression, because a predictor is derived from the titers through a non-linear chain: the latent-distance posterior, a stochastic map, a cluster assignment, and a geometric construction. We computed it by Monte Carlo. We drew *D* = 400 configurations *θ*^(^*^d^*^)^ from the antigenic posterior *p*(*θ* | titers), passed each through the full chain using the same analysis code, and took the standard deviation across draws of every predictor as its measurement error *S_ik_* . For candidate strains we used the standard deviation relative to the season centroid: candidates within a season share the prior-season centroid, the vaccine anchor, and the drift axis, so the shared component of their positional error cancels in the candidate contrast (*z _j_* − *z_w_*).

Writing 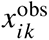 for the measured value of predictor *k* in season *i* and *x_ik_* for its unobserved true value, the model is augmented with a measurement layer and a structural layer,

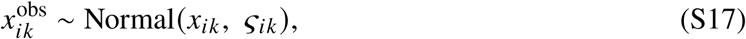

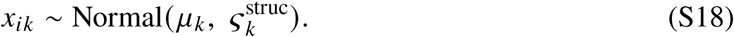

The measurement standard deviation *S_ik_* is the standard deviation of predictor *k* for season *i* across the *D* = 400 antigenic configurations of the previous paragraph, and it enters the model as data rather than as a parameter. Equation S17 has the same form as Equation S14, applied to the predictors instead of the outcome; Equation S18 is the structural equation that completes the classical measurement-error model (*78*). Because the predictors are standardized on the training fold, *μ_k_* is near zero and 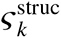 near one, but both are estimated rather than fixed: the observed spread that the standardization uses is itself inflated by the measurement error, so 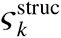 cannot be set to one. Of the two standard deviations, only *S_ik_* is known (near 0.25 on the standardized scale; per-predictor values in Table S3); 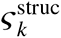 is the between-season spread of the true predictor values, estimated near 1.0. Treating *S_ik_* as known data also makes the effectiveness layer consistent with the antigenic layer upstream, where the mapping error enters as a known quantity.

A predictor enters through its latent value *x_ik_* only when *S_ik_* > 0; when *S_ik_* = 0 the observed value is used directly,

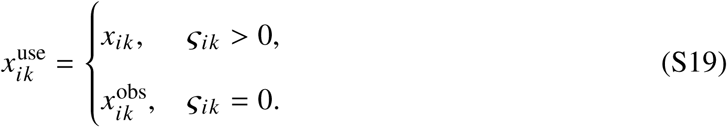

A measurement error of zero is a substantive statement, not a missing value. Vaccination coverage is not derived from the antigenic map and is treated as measured without error, so all its values are used as observed. The update-drift projection is zero by construction in the five seasons with no vaccine update, independent of the map configuration, so those five entries are used as observed while the eleven seasons with a vaccine update inform its structural parameters.

Replacing *z_i_* in the outcome equation by 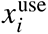 and integrating out the season residual *u_i_* gives the likelihood evaluated by the complete model,

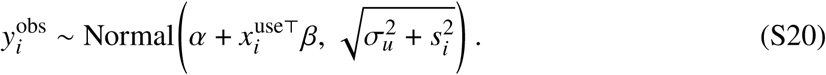

Marginalizing *u_i_* analytically eliminates per-season latent funneling, prevents the pointwise leaveone-out density from including the season’s own outcome, and identifies *σ_u_* as the variability beyond the measurement noise. The latent predictors *x* are retained, so the posterior is over (*α*, *β*, *τ*, *σ_u_*, *μ*, *S*^struc^, *x*). The latent season effect *ψ_i_* is recovered in closed form as the conjugate posterior of Equation S14 and anchors the counterfactual scoring (Eq. S24).

Fitting used the non-centred parameterization 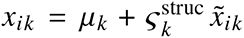 ∼ Normal(0,1); the centred form induces funneling in the latent cells because *S*^struc^ is unconstrained and weakly identified at *N* = 16. The structural layer extends the prior specification from the ridge subsection with

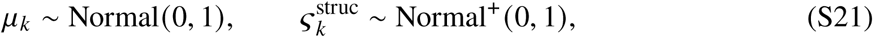

both weakly informative around the training-fold standardization.

Two properties of this specification matter for interpretation. It corrects rather than merely acknowledges the bias that measurement error induces: regressing on a mismeasured predictor attenuates its coefficient toward zero, and conditioning on the latent value removes that attenuation, which a procedure that only widened intervals would not do. The correction is multivariate, so the change in any one coefficient is not determined by that predictor’s own reliability but by the covariance of the whole predictor set.

We fitted the complete model, Equations S14–S21, by Hamiltonian Monte Carlo with the noU-turn sampler (*82, 83*) in Stan (*60*), four chains of 2,000 warmup and 2,000 sampling iterations; across all 17 fits the sampler produced no divergent transitions, a maximum *R̂* of 1.0026, and bulk effective sample size above 2,256 at adapt delta = 0.95. The fitted coefficients are reported in Table S2; because the prior shrinks them toward zero, they reflect predictive contribution rather than inferential effect sizes and are not interpreted as hypothesis tests.

### Propagating antigenic-map uncertainty into the inferential model

For the inferential model, which averages over specifications rather than fitting one, we propagated the same *D* configurations through the model-averaging procedure: we re-ran the entire averaging in each configuration *d*, obtaining the model-averaged coefficient *β̂*(*^d^*) and its bootstrap variance *V̂*(*^d^*^)^, and combined them as

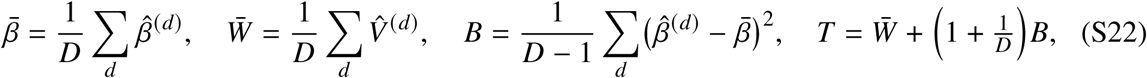

so that *W̄* is the within-configuration estimation error, *B* the between-configuration antigenic uncertainty, and the reported quantity is the between-configuration share 100 (*T* − *W̄*)/*T*. The arithmetic is Rubin’s combination rule, but the procedure is not multiple imputation: a valid imputation of a mismeasured covariate conditions on the outcome, and ours does not. We therefore report it as Monte Carlo error propagation and attach no degrees-of-freedom or fraction-of-missing-information statement.

### Reported measures of predictive accuracy

We report two out-of-sample coefficients of determination for the predictive model, each alongside the ceiling imposed by the VE measurement error, both on the log(OR) scale.

The primary measure is the leave-one-season-out *R*^2^: we refitted the complete model sixteen times, each time holding out one season’s VE and scoring the prediction against the held-out observation, the exact-refit form of the leave-one-out *R*^2^ (*64, 84*). Its total sum of squares is centred on the full-sample mean of *y*, the conservative convention, rather than on each fold’s training mean. The secondary measure is the prospective *R*^2^ of the expanding window, which holds out the antigenic map as well as the outcome and is scored over ten target seasons, 2015–2016 to 2024– 2025, nine of them with VE estimates (Supplementary Text, *Antigenic-map stability and leakage audit*). This is the stronger design; it is secondary only because it rests on fewer seasons, and we report it alongside its fold count.

Each observed-scale *R*^2^ carries the ceiling the VE measurement error imposes: where 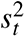 is the known sampling variance of season *t*’s published VE (Eq. S14), a model predicting true effectiveness exactly would leave, in expectation, 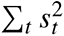 of squared error, against the same fixed SS_tot_ (the full-sample-mean convention above) that the corresponding *R*^2^ itself uses, so

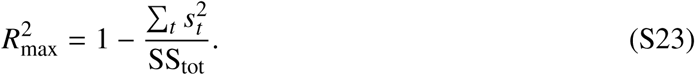

We report this measurement-noise ceiling beside each *R*^2^ (*78*).

Each *R*^2^ was also compared with its distribution under a permutation null (Eq. S13). For the leave-one-season-out *R*^2^, each of *R* = 500 replicates reassigned the 16 published VE estimates among seasons together with their sampling variances 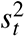, so that each estimate kept its own measurement error, and repeated all 16 refits and the scoring. Each refit used 1,000 warmup and 1,000 sampling iterations per chain, sufficient for the posterior mean on which each held-out prediction is based. Refitting the expanding window’s per-season antigenic maps within every replicate is not computationally feasible, so for the prospective *R*^2^ the forecasts were held fixed and the nine observed values were reassigned among them in all 9! = 362,880 ways. This exact test asks whether the forecasts are matched to their own seasons; unlike the leave-one-season-out test, it does not repeat the fitting.

### Scenario analysis of substitute candidates

For each eligible season *t* (*t* = 1, …, 16) we refitted the model on the *N* − 1 remaining seasons so that season *t*’s observed VE never informs the coefficients used to score it, then computed the linear predictor 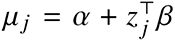 for every admissible candidate strain *j*, recomputing vaccine lead distance and the update-drift projection for each.

### Admissibility of candidates

A strain is admissible for season *t* = *Y*–(*Y*+1) only if its assay date—the date its HI or PRNT titer batch was run, recorded in the serological processing—falls in the two-year window ending on 15 January of year *Y*, that is, after 15 January of year *Y*−2 and on or before 15 January of year *Y*. Basing admissibility on the assay date captures the quantity the scenario analysis requires: whether a strain had been antigenically characterized early enough to be nominated. The NH vaccine composition meeting (VCM) is held in late February of year *Y*, and a strain must reach a WHO Collaborating Centre and be assayed before it can be evaluated there (*52*), so the 15 January cutoff falls just before the meeting and excludes any strain characterized after the decision point. A strain lacking an assay date was never characterized in a WHO panel and cannot be a candidate vaccine virus, so we excluded it; this requirement replaces a filter on the presence of a published sequence, which does not by itself establish antigenic characterization. The window length is fixed at two years so that pool size remains comparable across seasons (57 to 376 strains per season). Selecting candidates by season membership would admit strains characterized after the meeting, because under our season convention (collection month ≥ 10 → season *Y*–(*Y*+1)) season *t*−1 extends to 30 September of year *Y*, approximately seven months past the VCM. Writing *w* for the deployed vaccine, we formed two estimands from the posterior draws.

The *anchored* estimand conditions the season’s overall level on what that season actually delivered. Combining the observation *y_t_* with the model’s prediction for the deployed strain gives the conjugate posterior

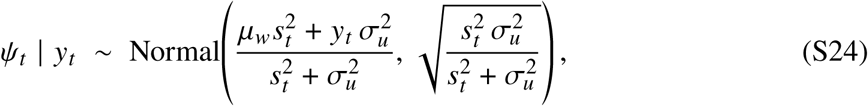

after which each candidate is scored as *η _j_* = *ψ_t_* + (*z _j_* − *z_w_*)^⊤^ *β*. The deployed strain and every substitute then share the same season level and differ only through the geometric contrast, which avoids comparing a fitted value for the deployed strain, shrunk toward its own observation, against an unshrunk extrapolation for a candidate. This is a retrospective counterfactual: it asks what a substitute would have delivered given what the deployed strain delivered, not what could have been forecast beforehand. The *prospective* estimand instead draws *u_t_* ∼ Normal(0, *σ_u_*) once per season and shares it across candidates, and is the quantity available before the season begins; the two agree to within 0.1 pp in the pooled gain. Predicted effectiveness follows as VE*_j_* = 100 (1 − *e^ηj^*), evaluated draw by draw so that the nonlinearity of that transformation is propagated rather than approximated.

Within a season the two estimands differ only by a level shift common to every candidate, and VE is monotone in *η*, so the within-season ranking is identical under either and only the magnitude of the gain depends on the choice. The one fixed predictor, coverage, likewise shifts every candidate by the same amount; only the two per-candidate predictors—vaccine lead distance and the update–drift projection—discriminate among strains within a season.

We report two candidates, differing in how each accounts for the uncertainty of the prediction. The model’s candidate is the strain whose posterior mean of predicted VE is highest, the Bayes-optimal choice when the utility is the realized effectiveness of a strain deployed once (*85, 86*). The tenth-percentile candidate instead maximizes the tenth percentile of that posterior, a risk-averse criterion that discounts a candidate whose apparent advantage rests on a wide posterior. We did not use expected improvement, E[max(VE*_j_* −VE*_w_*, 0)]: its truncation at zero removes the downside, so posterior variance can only raise a candidate’s score and strains far outside the range of the training data score highest. Expected improvement is an acquisition function for sequential experimental design, where the outcome will be observed and the search will continue (*87*); that is not the decision faced here. Because ranking many uncertain predictions and reporting the maximum is upward biased (*88, 89*), we also report the posterior of max*_j_* Δ*_j_*, the gain available if the best strain were known in advance, explicitly as an upper bound, together with the posterior probability that each candidate is the season’s optimum and the smallest set of candidates whose probabilities sum to 0.95.

This remains a model-based substitution rather than a causal counterfactual: the coefficients were estimated only on deployed strains, so ranking never-deployed candidates assumes the fitted distance- and drift-projection relationships extrapolate to those strains, including candidates beyond the observed predictor range. It further assumes that the unexplained component of log(OR) is invariant to the strain choice.

### Strain-selection trade-off and candidate coverage

For the strain-selection analyses (Fig. 5) we characterized three strains per season, drawn from the same candidate pool: the model’s candidate (highest leave-one-out-refit predicted VE, as above), the WHO-selected vaccine, and the coverage-optimal candidate, defined as the strain maximizing coverage of the most recently characterized (season *t*−1) viral population. Coverage was the fraction of the season-*t*−1 NH antigen strains lying within a match radius of *R* = 3 AU of a candidate— that is, strains not antigenically distinct from it, under the surveillance convention by which a reduction in HI titer of eight-fold or more, equivalently 3 AU on the base-2 antigenic-distance scale used here, marks a circulating virus as antigenically distinct from the vaccine reference strain (*25, 26*). To locate each strain relative to the population’s motion we used a drift-relative frame with origin at the prior-season centroid *c*(*t*−1) and axis along the recent drift direction *d̂* = (*c*(*t*−1)−*c*(*t*−2))/‖ *c*(*t*−1)−*c*(*t*−2) ‖; in this frame each strain’s offset from *c*(*t*−1) decomposes into an along-drift projection and an orthogonal (lateral) magnitude. Because coverage and predicted VE are two distinct selection objectives, plotting one against the other (Fig. 5C) exposes the tradeoff without circularity; predicted VE is used only to define the model’s candidate, never regressed on its own predictors.

### Standardized coefficients

Because predictors are measured on different scales (percentages, AU, fractions, binary indicators), raw regression coefficients are not directly comparable across variables. We report standardized coefficients computed as:

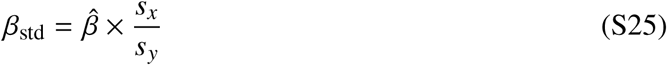

where *β̂*is the unstandardized regression coefficient, *s_x_* is the standard deviation of the predictor, and *s_y_* is the standard deviation of log(OR) in the analysis sample. Standardized coefficients represent the expected change in log(OR) (in standard deviation units) per one-standard-deviation increase in the predictor. This form applies to the predictive model, which is a single fixed specification. The inferential model instead uses the partial standard deviation (Eq. S11), because *s_x_* does not reflect the scale of a partial coefficient, which changes across the specifications being averaged (*36*); the two are reported in separate columns and are not interchangeable. Standardized coefficients for both models are compared in Table S10.

## Supplementary Text

### Serological assay data preparation

The WHO Collaborating Centre record for 2002–2025, after cleaning and replicate averaging to one value per virus–serum pair, contains 104,183 HI titers on 12,374 viruses against 241 antisera, and 12,022 PRNT titers on 2,304 viruses against 98 antisera. Fitting an antigenic map to the full record is not feasible—optimization cost grows superlinearly in the number of strains—so we selected a panel. Because the map is the study’s measurement instrument, we selected on measurement depth rather than by a fixed quota per year.

#### Basis for the ordering

An antigenic map places strains so that inter-strain distances reproduce the measured titers, so a strain’s coordinates are determined by the measurements connecting it to others. In *d* dimensions, *d* distances confine a strain to a mirror-image pair of locations and a (*d*+1)th resolves the ambiguity; with fewer, a continuum of coordinates reproduces the measurements equally well. Cartography algorithms still return an answer in that situation, supplied by initialization and regularization rather than by the data, and that answer can shift when the input set changes. We therefore counted, for each virus, the number of distinct antisera against which it returned a titer within range, hereafter its quantitative partner count. However, these counts range from 0 to 57 in HI (median 8), so a fixed per-year quota admits very different quality in different years. Hence, the selection rules below were applied.

#### Selection rules

Applied in order, to each assay. (i) Vaccine strains and viruses characterized by both assays with a quantitative partner count of at least eight across the two assays combined were admitted irrespective of rank and irrespective of their year’s allocation. Vaccine strains are required because every vaccine-update variable is a difference involving a vaccine position. Dual-assay viruses are required because the latent model relates HI to PRNT through antigen–antiserum pairs measured in both assays, and only such pairs inform that relationship; the partner-count condition excludes dual-assay viruses too sparsely measured to be placed. (ii) Remaining viruses were ordered within each year by quantitative partner count, ties broken by total titer count and then by the number of distinct years the partnering antisera span. (iii) A budget of 2,000 viruses per assay was distributed across years: each year first received a floor of 25 viruses, or all it had if fewer, and the remainder was allocated in proportion to the square root of the viruses each year still held. The square root makes the allocation concave, matching the 1/^√^*n* decline in the standard error of season-level quantities, so capacity is directed to sparse seasons rather than to seasons already precisely characterized. (iv) Antisera with fewer than three retained virus partners were dropped, as they anchor little and are themselves poorly placed. (v) Each panel was reduced to the largest connected component of its virus–antiserum measurement graph, since a distance spanning two components has no measurement path supporting it. (vi) Once both panels existed, viruses characterized by a single assay were required to have at least eight quantitative partners, with vaccine and dual-assay strains exempt; steps (iv) and (v) were then repeated, as removing viruses can disconnect a panel. All quantities used are functions of titer data alone; no vaccine-effectiveness outcome entered selection at any stage.

#### Retained panels

HI retained 21,459 titers on 2,000 viruses and 241 antisera; PRNT retained 6,607 titers on 1,160 viruses and 98 antisera. Median partner count rose from 9 to 10 in HI while 16% of viruses were retained, reflecting a record holding more well-measured viruses than the budget admits. In PRNT the median rose from 5 to 6, because the budget barely binds there; PRNT positional precision is set by the assay record rather than by selection. The union of the two panels, which the latent model consumes, comprises 2,053 viruses and 269 antisera, 23,215 antigen–antiserum pairs, and 2,322 map points. Of the retained viruses, 1,107 were characterized by both assays, 893 by HI alone, and 53 by PRNT alone. The dual-assay pairs that identify the HI–PRNT relationship number 4,851 of the 5,308 available, retained in greater proportion than the 83% reduction in map points. Across both assays, the share of viruses with fewer than eight partners fell from 30.7% to 4.1%. All 24 years are retained with at least 28 viruses each, and all 19 recommended H3N2 vaccine components falling within the study window are present.

#### Limitations

Partner count measures how many antisera constrain a virus, not how those antisera are arranged: a virus measured against eight antisera drawn from one antigenic cluster is well connected by count and poorly conditioned in fact. The number of distinct antiserum years is recorded and used to break ties, but is not enforced, since enforcing it requires the positions being estimated. The *d*+1 criterion is necessary rather than sufficient. The 3.4% of retained viruses below it are chiefly protected vaccine and dual-assay strains, carrying real positional uncertainty that is propagated rather than assumed away. Seasons 2002–2007 sit near the per-year floor at 28–33 viruses against 207–295 for 2022–2024, so early season-level quantities rest on fewer strains. The PRNT-only stratum is reduced to 53 viruses, limiting power to detect antigenic behaviour specific to neutralization.

### Biological basis for integrating HI and neutralization assays

The four-observation-process latent-distance model assumes that HI and PRNT titers are noisy observations of a single shared antigenic distance. This section summarizes the mechanistic and empirical evidence supporting that assumption.

#### Mechanistic complementarity

HI measures antibody-mediated blockade of viral attachment to *α*2,6-linked sialic acid receptors—exclusively detecting antibodies targeting the HA head receptor-binding site (*90*). PRNT measures functional prevention of productive infection through multiple effector pathways, including receptor-binding blockade, membrane fusion interference, and contributions from broadly neutralizing antibodies targeting the conserved HA stalk domain (*91*). Because PRNT captures a broader spectrum of antibody effector functions, it is more biologically comprehensive than HI (*48*) but also intrinsically noisier (*92*): the assay requires live virus, cell culture, and plaque counting over 2–4 days (*93*), introducing variability that HI’s standardized red blood cell agglutination readout avoids. Inter-laboratory geometric coefficients of variation range from 83–192% for neutralization assays versus 83–125% for HI (*92*), a reproducibility gap repeatedly documented in collaborative serology studies (*94, 95*). This difference in measurement precision is directly reflected in our model’s estimated noise parameters (*σ*_2_ = 1.55 for PRNT versus *σ*_1_ = 0.35 for HI). Despite these mechanistic differences, the two assays share a common dependence on the same underlying HA antigenic structure, which predicts strong—but not perfect—empirical correlation.

#### Empirical convergence across influenza studie

. Published cross-assay correlations for influenza viruses consistently fall in the range *r* = 0.73–0.96 (Fig. S14). For seasonal influenza, Truelove et al. (*96*) reported Spearman *ρ* = 0.86 between neutralization and HI titers; for H1N1pdm09, Veguilla et al. (*97*) found *ρ* = 0.84. The FLUCOP consortium harmonization study (*92*) measured Pearson correlations between ELISA-based microneutralization and HI across influenza subtypes: *r* = 0.81 (H1N1), 0.90 (H3N2, egg-grown), 0.92 (H3N2, cell-grown), and 0.95 (influenza B). For H5N1, pseudoparticle neutralization (PPN) correlated with HI at *r* = 0.96 (*98*) and *r* = 0.73 (*99*), and with conventional microneutralization at *r* = 0.88 (*98*) and *r* = 0.78 (*99*). Threshold equivalences are also well established: an HI titer of 1:40 (=2 fold distance)—the classical seroprotection threshold (*100*)—corresponds to a neutralization titer of approximately 1:20 (=1 fold distance), reflecting the systematic offset between the two assay types (*96*) (Fig. S15). The same coupling extends beyond influenza: Figure S14 compiles published cross-assay correlation coefficients across influenza viruses, flaviviruses, and coronaviruses, with strong HI–neutralization agreement also reported for dengue (*101*), Japanese encephalitis virus (*102, 103*), tick-borne encephalitis virus (*104*), and SARS-CoV-2 (*105*). This convergence—across diverse influenza subtypes, assay variants, laboratory protocols, and unrelated viral families—indicates that tight HI–neutralization coupling is a general property of antibody responses to enveloped viruses rather than an influenza-specific artifact, and supports the shared latent variable assumption.

#### The H3N2 hemagglutination crisis

The practical necessity of integrating HI and PRNT data arose from the progressive failure of HI to characterize post-2005 H3N2 viruses. Receptorbinding-site substitutions progressively reduced binding avidity to human-like *α*2,6-sialic acid receptors by approximately 200-fold between 2001 and 2004 (*16*), and the later K160T substitution added a glycosylation site in antigenic site B that altered antibody recognition (*53*). Concurrently, neuraminidase mutations (D151G, H150R) conferred receptor-binding functionality to NA, causing false NA-mediated agglutination that confounded HI measurements of HA-directed antibodies (*106*). Countermeasures—including oseltamivir addition to block NA-mediated agglutination (*107*) and glycan-remodeled erythrocytes (*108*)—partially mitigated these issues but did not fully restore HI reliability for contemporary H3N2 strains. This forced the WHO Collaborating centers to adopt neutralization assays for antigenic characterization of H3N2 (*6*), producing the mixed-assay datasets (HI-only, PRNT-only, or both) that necessitate formal statistical integration.

### Map distances as independently informative indicators of the latent distance

Combining HI and PRNT *titers* needs little justification—they are different assays of the same hemagglutinin (HA). The substantive choice is to treat the two Topolow *map* distances as additional observations, since each map is built from the titer data and might merely re-encode it. However, the two-titer model is under-identified (three second moments—Var(*y*_1_), Var(*y*_2_), Cov(*y*_1_, *y*_2_)—for four unknowns), and M LAT 2OBS indeed failed to converge (*R̂* = 2.25) (*65*). Adding the map distances is warranted only because they are valid measurements of the same *θ_i_* that also carry information the titers lack.

That extra information is structural. Topolow places each strain using the entire titer table, so a pair’s map distance is a network-consensus estimate constrained by every titration involving the two strains, whereas its titer is a single, discrete (two-fold) local readout (*40*). The consequence is visible in Fig. S16: although titer and map distance are strongly associated (Pearson *r* = 0.98 for HI, *r* = 0.98 for PRNT), the discrete titer levels fan out into a continuous range of map distances, so the map resolves within-level differences the titer cannot represent. A mutual-information estimate (*k*-nearest-neighbour, *k* = 5, normalized by the Kozachenko–Leonenko entropy of *y*_3_; preferred over a correlation because it captures non-linear dependence and the relationship is compressed and censored at the extremes) quantifies this surplus: *H*(*y*_3_ | *y*_1_)/*H*(*y*_3_) = 0.20, so about a fifth of the HI map distance’s information remains after its paired HI titer is known (an approximate magnitude, given the sensitivity of *k*-NN estimators to ties on the discrete titer grid). The observation model keeps each map an unbiased indicator by absorbing its scale, offset, and noise into *a _j_*, *b _j_*, *σ_j_* (HI map *a*_3_ = 0.869 [90% credible interval (CrI) 0.866, 0.871], *σ*_3_ = 0.226; PRNT map *a*_4_ = 0.630 [0.616, 0.644]); the HI map’s low noise relative to the raw HI titers (*σ*_3_ = 0.226 vs. *σ*_1_ = 0.348) indicates that cartographic mapping denoises the measurements.

### Choosing the number of antigenic clusters

Antigenic space here does not read as a set of distinct, well-separated clusters. That has a direct consequence for how *k* should be chosen. The common internal criteria we tracked (average silhouette width, the Calinski–Harabasz index, and the Davies–Bouldin index) all decline monotonically toward the coarse partitions, so each reaches its optimum at the coarsest end of the range and, within the range where structure is actually resolved, none separates one candidate count from another. This is the expected behaviour of these criteria rather than an artefact of our data. Silhouette width, Calinski–Harabasz, and Davies–Bouldin all rest on a within-cluster versus between-cluster contrast of compactness and separation, and that contrast presupposes clusters that are convex, of comparable size, and well separated from one another; on a continuum without such groups the indices are known to degrade, collapsing toward the coarsest partition or otherwise failing to discriminate (*109,110*). We therefore bounded *k* from above with two criteria that make no assumption about cluster shape, and then required the surviving partition to be interpretable (Fig. S1).

The first criterion is cluster occupancy. Rebuilding the evidence accumulation consensus at every *k* from 5 to 16 and recording the size of its smallest cluster identifies *k* = 12 as the largest count at which no cluster falls below 50 strains: the smallest holds 64 strains there, whereas every larger count drops at least one cluster below 50, to as few as 2 strains at *k* = 13 and 1 at *k* = 14 (Fig. S1A). Past *k* = 12 the consensus no longer subdivides antigenic groups but sheds residual strains into vestigial ones. Occupancy is not monotone below that ceiling either: *k* = 7 and *k* = 9 each isolate a cluster of fewer than 30 strains (18 and 25).

The second criterion asks whether a cluster count is a genuine level of the hierarchy. Cutting the Ward.D2 dendrogram at rising heights, each attainable count persists over an interval of cut heights whose width measures how robustly that count is defined. Fig. S1B shows a dendrogram cut height and Fig. S1C shows the frequency of the wide cut areas over 200 map re-estimations. Twelve clusters frequently persist over 1.43 antigenic units (AU) of cut height, a relatively wide interval. This criterion propagates the uncertainty of the cartographic mapping into the choice of *k*. For each of the 200 independent re-mappings that form the consensus, we located the cluster counts at which the dendrogram height difference carries a prominent local maximum—a merge whose cost stands well above that run’s typical merge height, following Mojena (*111*)—and counted the re-mappings voting for each count (Fig. S1C). The votes are broadly distributed across the mid-range and peak at *k* = 12, then fall away, dropping by roughly two-thirds at *k* = 13 and into single figures by *k* = 16.

Twelve clusters therefore attract the most support across re-estimations of the map. The two criteria agree on where the evidence stops: *k* = 12 is the last count that both keeps every cluster populated and is most often recovered across re-estimations of the map, and each argues against every larger count.

Cluster composition then decides that the surviving partition is the interpretable one. At *k* = 12, ten clusters contain a World Health Organization vaccine strain and are named for it, and the remaining two are named for their medoid. Because the vaccine is updated once the antigenic difference to the strains expected to circulate reaches about two antigenic units, each antigenic cluster is expected to contain at least one vaccine strain (*4*). The recovered clusters correspond to recognised antigenic groups where the record allows the comparison—the FU02, BR07, and PE09 clusters reproduce the previously reported Fujian/2002 cluster (*4,12*) and the Brisbane/2007 and Perth/2009 clusters (*12*)—and the remainder track successive vaccine strains through to A/England/185/2024.

The criteria bound *k* from above; they do not single out 12 against 11 or 10, which neither panel separates. A conventional silhouette analysis does not resolve the choice either: on an antigenic continuum the silhouette stays at the level Kaufman and Rousseeuw associate with the absence of substantial cluster structure (*109, 110, 112*), the expected reading here and the reason we did not use it to select *k*.

### Evidence accumulation clustering of antigenic positions

To assign the 2,053 antigen positions (*θ_i_*) in five-dimensional antigenic space to the adopted *k* = 12 clusters, we applied evidence accumulation clustering (*113*), an ensemble approach that aggregates many base partitions into a single consensus partition; the reference antisera were not assigned to antigenic clusters, serving only as reference points in the map. We generated 200 base partitions, each from an independently re-estimated antigenic map: for every run we re-mapped the posterior-mean latent distance matrix into five-dimensional space with Topolow (*40*) from a fresh random initialization, standardized the resulting coordinates, and applied *k*-means at *k* = 12 with *n*_start_ = 1,000 random initializations. Re-mapping on every run—rather than repeatedly clustering a single fixed map—propagates the stochastic uncertainty of the cartographic mapping into the ensemble alongside *k*-means initialization variability. From the 200 partitions we formed a coassociation matrix recording, for each virus pair, the fraction of runs in which the two co-clustered, and obtained the consensus partition—used throughout—by average-linkage hierarchical clustering of the associated dissimilarity (1 minus the normalized co-association), cut at *k* = 12; average linkage is appropriate here because co-association entries are similarity counts rather than squared-Euclidean distances. The 200 base partitions were concordant: the mean pairwise adjusted Rand index over the 19,900 run pairs was 0.73 (range 0.61–0.83), confirming that cluster assignments are stable to both mapping and *k*-means initialization stochasticity.

### Collection-date assignment and season labeling of antigenic strains

Each strain required a collection date to assign it to an influenza season. Collection dates were taken from the same WHO Collaborating Centre reports that supplied the titer tables: the per-virus collection dates printed in those reports were extracted and standardized to YYYY-MM-DD, or to YYYY-MM where the source withheld the day, in which case the day was set to 15 rather than fabricated (the day never affects a season label, only the month does). We matched the titer tables’ strain names to these dates through a three-stage cascade—an exact match on a normalized key (uppercase, delimiters removed, slashes retained), then a fallback on a fully punctuation-stripped key, then a constrained fuzzy stage that accepts a candidate only when it shares the same isolate number and year, its location token is within two edits of the query’s, and the recovered date is unambiguous—and the matched collection month set the season label (Northern Hemisphere: month ≥ 10 → season *Y*–(*Y*+1), else (*Y*−1)–*Y*; Southern Hemisphere: month ≥ 3 → *Y*–(*Y*+1), else (*Y*−1)–*Y*). Two properties of the dated strains justified how we treated the remainder. First, among strains carrying a collection date printed in the reports, the collection year equaled the year in the strain name for ∼98% of strains (98.0% for HI, 98.7% for neutralization), so the strainname year is a reliable temporal anchor. Second, the season a dated strain fell in was predictable from its hemisphere: most Northern Hemisphere strains named year *Y* belonged to the (*Y*−1)–*Y* season (62.7% for HI, 55.4% for neutralization; modal collection month January for HI, March for neutralization), whereas most Southern Hemisphere strains belonged to the *Y*–(*Y*+1) season (87.4% for HI, 93.5% for neutralization; modal collection month June for both).

For the strains whose names matched no dated record (26.8% of HI and 5.0% of neutralization titer measurements), we therefore imputed a collection date equal to the empirical majority season for the strain’s hemisphere, placed at that season’s modal collection month (mid-January for HI and mid-March for neutralization in the Northern Hemisphere, mid-June for the Southern in both assays) in the strain-name year. We learned these placements from the dated strains of each assay separately rather than fixing them a priori. Because only the strain-name year and hemisphere are available for an unmatched strain, this deterministic rule cannot exceed the majority-season share for any individual strain: an estimated 37–45% of imputed Northern strains and 7–13% of imputed Southern strains may belong to the adjacent season. The rule fixes season direction, not the individual-strain label, and the season labels of dated strains—the large majority of measurements—were unaffected.

### Antigenic dynamics and the vaccine–circulation lag

The descriptive analyses underlying Figs. 1 and 2 place the VE models in their evolutionary context; the main text reports their three recurring features—punctuated antigenic advance, pervasive cluster co-circulation, and the recurrent vaccine–circulation lag. Two caveats bear on any per-season reading of the lag. First, the cluster assignment of an individual vaccine strain can shift between map re-estimations, so the 6-of-21 match count reflects the consensus partition and may shift by a season under re-estimation, whereas the directional lag and the mean vaccine–circulation distance (3.8 AU; median 3.7) are robust. Second, cluster match alone is only weakly related to VE, because scalar antigenic distance predicts effectiveness only once the orientation of the vaccine update is held fixed, so the lag is best read as a structural feature of strain selection rather than a direct determinant of effectiveness.

### Inferential model: detailed results

#### The specification space

All 377 specifications were estimable: none was rank deficient, and the fitting path was verified against lm() and the study’s AICc convention to a tolerance of 10^−8^ before any weight was used. Fit varied widely across the space (*R*^2^ from 0.000 to 0.757, median 0.368), with a weighted mean of 0.680. No single specification dominated: the best-supported one—prior-season lead distance and the one-season update–drift projection, a two-predictor model—carried 22.6% of the AICc weight, leaving 77.4% distributed over the other 376. That distribution is the reason a single fitted specification is not reported as the result.

#### Calibration against permuted VE

Across 2,000 random reassignments of VE among seasons, each repeating the whole procedure (Methods, *Permutation calibration*), the averaged prediction reached a mean *R*^2^ of 0.43 and a 95th percentile of 0.72; 61 reassignments reached the observed 0.75 (*P* = 0.031; Fig. S9A). The largest *R*^2^ of any single specification averaged 0.52 under the null, and 67 reassignments reached the observed 0.757 (*P* = 0.034). Both null centres sit well above zero because a search over 377 specifications fits noise, so the observed fits are judged against those centres rather than against zero.

#### Model-averaged estimates

Table S1 gives, for each candidate, the averaged *β̄*^*^, its bootstrap interval and *P* value, the fraction of containing specifications sharing its sign, and the fraction in which it is individually significant. The two update–drift projections were negative in all 79 of their containing specifications, as was prior-season lead distance. Individual significance follows the three vaccine-update-orientation terms rather than the two projections alone: the one-season projection was individually significant in 75% of its containing specifications, update size in 68% and the two-season projection in 58%, against 51% for the pandemic-season control and 27% or less for every other candidate. Three variables—in-season dominant-cluster share, in-season antigenic diversity and in-season coverage—failed the unimodality condition and are reported as distributions rather than as averaged effects (Fig. S7).

#### Specification versus sampling uncertainty

These are distinct and behave differently here. Sign consistency is computed across specifications with the 16 seasons held fixed; the bootstrap interval is computed across resamples of those seasons. The one-season update–drift projection is negative in 100% of specifications, and 99.4% of bootstrap resamples also keep it negative, giving a 95% interval of [−0.94, −0.13] that excludes zero (*P* = 0.014). The two agree here, but they need not: the two-season projection is likewise negative in 100% of specifications while 5.0% of resamples cross zero, so its interval is [−0.96, 0.10] and *P* = 0.10. Vaccine-update size separates them further, at 99% sign consistency and *P* = 0.31. Sign consistency measures robustness to what is controlled for; the interval and the *P* value measure robustness to which 16 seasons were observed. The latter, not the sign count, is the summary statement.

#### Both quantities condition on one antigenic map

Varying the specification changes which covariates are controlled. Resampling changes which seasons are drawn. Neither changes the predictor values: every fit behind Fig. S7—the 377 specifications and the 1,000 bootstrap resamples—reads one predictor matrix, computed from the posterior-mean latent distances and a single map of them. Those predictors are estimates rather than measurements, and their uncertainty enters none of the intervals reported here. The corresponding check on the outcome side is the inverse-variance weighting reported under *Sensitivity analyses*, which changes no reportable sign. The check on the covariate side is reported in Methods, *Propagating antigenic-map uncertainty into the inferential model*: refitting the entire model-averaging procedure on 400 independent draws from the antigenic posterior decomposes the variance of each model-averaged coefficient into a within-world estimation component and a between-world antigenic component. Antigenic measurement accounted for between 3.5% and 25.8% of the total variance across the 13 candidates, so the intervals reported here understate the full uncertainty modestly, by the amount just quantified.

#### Sign consistency against the bootstrap P

Six candidates are negative in all 79 of their containing specifications. Their *P* values range from 0.010 to 0.44, so sign consistency cannot rank them, which is why the *P* value carries the inferential statement and the sign count is reported as a robustness diagnostic. Two candidates reach *P* < 0.05: the one-season update–drift projection at 0.014 and the pandemic-season control at 0.010. The next smallest are prior-season lead distance at 0.084 and the two-season projection at 0.102. Vaccine-update size reaches only 0.31, despite being individually significant in 68% of its containing specifications, because the specifications in which it is large are not the ones that carry the AICc weight. These are per-variable values. Against permuted VE, the one-season projection had *P* = 0.007 on its own and *P* = 0.089 once all 13 candidates are accounted for (Methods, *Permutation calibration*). We therefore read them as descriptive of how stable each estimate is under resampling rather than as confirmatory tests, consistent with an estimand that is a partial association and not a selected model. The control is a special case: an indicator for one season is estimable only in the 612 resamples that contain 2019–2020, so its interval and *P* value are conditional on that season being drawn and are not comparable with the other rows.

#### Conditional contrasts

Restricting to specifications that also control for vaccine-update orientation moves concurrent mismatch from *β̄*^∗^ = −0.21 to −0.35 and prior-season lead distance from −0.38 to −0.54 (table S12). The contrast runs in both directions: controlling for vaccine–virus match moves the one-season projection from −0.60 to −0.74, while leaving the two-season projection at −0.37. The largest single contrast is vaccine-update size, which strengthens from −0.31 to −0.64 once drift geometry is controlled and weakens from −0.50 to −0.25 once vaccine–virus match is. Both update–drift projections strengthen when viral population structure is controlled (−0.30 to −0.65 at the two-season horizon), and the pandemic control weakens from −0.55 to −0.37 once vaccine-update orientation is.

#### Assumption satisfaction across the space

Reported as a distribution rather than for one fit: 8 of 377 specifications had Shapiro–Wilk *p* < 0.05, none had Breusch–Pagan *p* < 0.05, 8 had a Durbin–Watson statistic outside [1.2, 2.8], 24 had a maximum VIF above 5, none had a condition number above 30, and 8 had a maximum Cook’s distance above 1. The 79 specifications containing the pandemic-season indicator reproduce 2019–2020 exactly, because an indicator for a single observation gives that observation a leverage of 1; Cook’s distance and the leave-one-out residual are undefined there by construction, and that season is excluded from those two summaries rather than counted as influential. The best-supported specification satisfies the standard assumptions (Shapiro–Wilk *W* = 0.967, *p* = 0.79; Breusch–Pagan *χ*^2^ = 0.64, *p* = 0.73; Durbin–Watson = 2.14; two seasons above Cook’s *D* > 4/*n*), and its residual diagnostics are shown in Fig. S6.

#### Sensitivity analyses

Re-averaging over the 345 specifications free of multicollinearity and high-influence flags (maximum VIF ≤ 5, condition number ≤ 30, maximum Cook’s distance ≤ 1; adding the normality, homoscedasticity and autocorrelation criteria would leave 329) changes no reportable estimate by more than 0.07; the largest movement anywhere is in-season coverage, which shifts by 0.09 and changes sign, and whose averaged estimate is not reported. Restricting to the 255 specifications carrying at most one indicator per mechanistic category—so that two measurements of one construct never compete—strengthens all three vaccine-update-orientation terms and changes no sign, moving the one-season projection to −0.74, update size to −0.62 and the two-season projection to −0.62. Inverse-variance weighting by the published VE confidence intervals changes the sign of no reportable variable; it leaves the one-season projection at −0.69 but strengthens the two-season projection to −0.99, a shift of 0.62 and the largest any sensitivity produces. Uniform weights in place of AICc weights shift estimates by up to 0.24, and the two signs that change belong to variables whose averaged estimate is not reported. Omitting each season in turn and re-running the whole procedure moves the two-season projection most, by 0.35 after 2011–2012, and moves the one-season projection least among the update terms, by 0.06; the only sign change among the reportable variables is prior-season antigenic diversity, whose averaged estimate is −0.06 and therefore sits at the sign boundary already; in-season coverage and in-season dominant-cluster share also change sign, and neither has a reported averaged estimate.

#### Power

In a three-predictor specification at *N* = 16, an effect must reach *f* ^2^ = 0.67, equivalently a partial *R*^2^ of 0.40, for 80% power at *α* = 0.05. Power against a conventionally moderate effect (*f* ^2^ = 0.15) is 0.27. Only very large effects are detectable in any individual specification, which is why individual significance counts are reported as one column among several rather than as the criterion. These power figures describe single specifications; the averaged quantities are calibrated directly by permutation (*Calibration against permuted VE*, above).

### Baseline scalar distance model comparison

To isolate what antigenic geometry adds over standard scalar distance metrics, we fit a baseline ordinary least squares model using only the two scalar distances in the candidate pool: concurrent vaccine-to-centroid distance (mismatch inseason) and prior-season vaccine-to-centroid distance (vaccine lead distance). Because the inferential model targets explanation rather than out-of-sample forecasting (*114*), both sides were evaluated by in-sample explanatory power.

The comparator on the geometry side is the AICc-weighted model-averaged prediction, *ŷ* = Σ*_m_ w_m_ ŷ _m_*, which exists without any selection step and is the quantity the inferential analysis reports. The baseline explained little of the observed variation (*R*^2^ = 0.20, adj. *R*^2^ = 0.08, RMSE = 0.186), while the averaged prediction tracked observed VE across its dynamic range (*R*^2^ = 0.75, RMSE = 0.103; Fig. S8A).

The *R*^2^ of the averaged prediction (0.75) exceeds the weighted mean of the individual specifications’ *R*^2^ (0.68) because an average of several fits is smoother than any one of them; the two are different quantities and neither substitutes for the other. Taken with the conditional contrasts above, the comparison shows that scalar distance carries almost no signal on its own, while its partial association is larger and negative in every specification containing it (*β̄*^∗^ = −0.54 for prior-season lead distance, 95% CI −0.87 to 0.09). Whether the weak marginal signal reflects masking by update orientation or a genuinely small effect is not resolvable at *N* = 16.

### Why scalar antigenic distance appears weak

The positive theoretical basis for the geometric features is given under *Mechanistic category definitions*; here we explain why scalar antigenic distance behaves as it does. Taken alone it orders seasons poorly. The concurrent vaccine-to-centroid distance ranges from 1.5 to 5.4 AU, yet seasons with nearly identical mismatches span nearly the full range of observed effectiveness: 3.05 AU with VE ≈ 4% in 2018–2019 against 3.34 AU with VE ≈ 57% in 2019–2020, the larger mismatch carrying the higher effectiveness. Its marginal rank correlation with log(OR) is correspondingly small and indistinguishable from zero (*ρ* = −0.34, *P* = 0.20; Fig. S10), and a model restricted to the two scalar distances explains *R*^2^ = 0.20.

The mechanism is straightforward once stated in terms of what each variable measures. Distance records how far the vaccine sits from the circulating population; orientation records whether the update moved toward where that population was heading. Two seasons at the same distance differ in effectiveness according to whether the vaccine was placed ahead of the drift or behind it, so distance predicts VE only among seasons whose updates were comparably oriented. Holding orientation fixed supplies that comparison, which is why the partial association is roughly two-thirds larger than the marginal one. The practical consequence is that antigenic distance should not be dismissed as uninformative for strain selection, but neither should it be read without reference to the direction of the update that produced it. Consistent with a marginal reading, the ferret-serology antigenic distance of Bonomo and Deem (*41*) explained only *r*^2^ = 0.23 of H3N2 VE over the most recent decade.

### Cross-reactive immunity and the protection floor

The geometric model captures HA-head-directed antigenic variation, but clinical protection against influenza also depends on broadly cross-reactive immune responses that are largely independent of HA head antigenic distance. HA stalk-directed antibodies, which target the conserved membrane-proximal domain of HA, provide heterosubtypic protection that decays slowly with antigenic drift (*91*). Cross-reactive CD8^+^ T cells targeting conserved internal proteins (NP, M1, PB1) maintain cytolytic function across antigenically divergent strains, a component of protection independent of HA/NA antigenic distance (*115*). Together, these responses establish a protection floor that explains why VE rarely reaches zero even in severely mismatched seasons. This dual-decay architecture—fast HA-head-specific antibody decline overlaid on slow stalk- and T-cell-mediated decay—means that the geometric model captures the component of protection most sensitive to antigenic evolution (HA head), while the residual variance partly reflects host-side immune mechanisms operating below this floor. Incorporating stalk antibody titers or T-cell response signatures as covariates could further partition this residual variance, although such data are not routinely available at the population level.

### Rationale for a Bayesian predictive model

The predictive analysis asks three things of a 16-season series: how well pre-season geometry forecasts effectiveness, how candidate strains rank within a season, and how much effectiveness a substitute strain would have supplied. Four features of this problem shaped the choice of model.

#### The outcome is measured with known, substantial error

Each season’s effectiveness is not observed but estimated, and published with a confidence interval. The mean variance implied by those intervals is 0.032 on the log(OR) scale, comparable to the total variance a regression on these predictors leaves unexplained. Equation S14 uses the reported intervals as data, which is the standard treatment for combining estimates that carry their own uncertainty (*76–78*) and which yields the separation between explained and unresolvable variance reported above.

#### Sixteen seasons cannot separately identify three correlated coeflcients

Regularization is therefore not optional, and the amount of it is a consequential choice. Estimating the shrinkage scale jointly with the coefficients propagates uncertainty about that choice into every forecast (*79*).

#### Ranking requires a joint distribution over candidates

Selecting the best of 100 to 500 candidates from point predictions returns a single strain and no statement of whether it is distinguishable from the next several. The joint posterior returns the *probability* that each candidate is the season’s optimum and hence the set of strains genuinely in contention.

A reported advantage over the deployed vaccine is a difference between two uncertain quantities.

Both the size of that difference and the probability that it is positive require a distribution over both terms, and the maximum over many uncertain predictions is upward biased (*88,89*). Posterior draws supply the interval, the probability, and the separation between the gain from an implementable rule and the larger gain available under perfect foresight.

### Predictive model: detailed results

#### Sampling and convergence

All four chains converged for every fit, with R̂ < 1.01 under rank normalization (*116*), bulk effective sample size above 2,256, and no divergent transitions.

#### The held-out quantity is genuinely held out

Because the data contain exactly one observation per season, leaving out an observation and leaving out a season are the same operation. Integrating *u_t_* out, as in Equation S16, is what makes the result a held-out quantity: a model retaining a free season-level effect would evaluate each season conditionally on a parameter fitted to that same season. The reported out-of-sample statistics come from 16 exact refits, each holding out one season and scoring it from a model estimated on the other fifteen.

#### Prior sensitivity

The coefficient signs were stable across twelve combinations of the *τ* and *σ_u_* prior scales, and the model’s candidate picks and pooled gain were stable under a tight and a vague prior (Table S4).

#### Predictive performance (out-of-sample)

Holding out each season in turn gave *R*^2^ = 0.47 and RMSE = 0.152 on the log(OR) scale (table S13), with Pearson *r* = 0.711 between predicted and observed log(OR). Across 500 shuffled reassignments of the published VE estimates and their sampling variances among seasons, each refitted through all 16 folds (Methods, *Reported measures of predictive accuracy*), the held-out *R*^2^ had mean −0.25 (standard deviation 0.18) and maximum 0.31. None reached 0.47, hence *P* = 0.003 (Fig. S9B).

#### Forecast calibration and skill

A model that reports a distribution rather than a single number must be judged on the whole distribution, so we scored the forecasts with the continuous ranked probability score (CRPS), a strictly proper scoring rule: it is minimized only by reporting one’s true predictive distribution, and so cannot be improved by overstating confidence (*117*). CRPS is measured in the units of the quantity forecast—here log(OR)—and is lower for better forecasts, reducing to the absolute error when a forecast is a single number. Its value is interpretable only against a reference, so we compared each forecast to a *historical-mean* forecast for the same season: the distribution obtained from the mean and spread of VE in the training seasons alone, which is what one would predict knowing the historical record but nothing about antigenic geometry. Averaged over the 16 held-out seasons the model scored 0.088 against 0.128 for the historical-mean forecast, a paired difference of 0.040. Antigenic geometry therefore adds substantial information beyond the historical distribution of effectiveness. The corresponding prospective score under the expanding window, against its own fold-training-mean forecast, is reported in the main text (*Forecast performance under strictly prospective validation*).

Following the principle that a forecast should be as sharp as possible subject to being calibrated (*118*), we report interval width alongside coverage. The 95% predictive intervals contained the observed value in all 16 held-out seasons and in 9 of 9 expanding-window seasons, with a mean width of 0.739 on the log(OR) scale. The probability integral transform—the quantile of the predictive distribution at which each observation fell, which is uniformly distributed if the forecasts are calibrated (*119*)—had a standard deviation of 0.285 against 0.289 for a perfectly calibrated forecast, indicating intervals close to nominal width rather than overconfident. At 16 seasons these calibration statistics are themselves imprecise and we read them descriptively.

#### Residual variation is no larger than the VE measurement noise

The variation the model could not explain was not distinguishable from the noise in the VE estimates themselves. Because Equation S14 treats each published VE as a measurement of an underlying true effectiveness, the model separates its own residual error from that noise: the season-to-season variation left unexplained by geometry was small and indistinguishable from zero (*σ_u_* posterior mean 0.065, 95% CrI [0.004, 0.155]), beside a mean measurement variance of 0.022 in the published estimates. The posterior median for *σ_u_* (0.061) sat close to its prior median (0.065, posterior-to-prior ratio 0.94), so this small residual variation is consistent with the prior. Note that *σ_u_* and *σ*_meas,*t*_ are not independent: both are estimated from the same residual scatter, so a larger assumed measurement error leaves less for *σ_u_* to absorb. The posterior for *σ_u_* therefore bounds rather than measures the unexplained season-to-season variation.

Every observed-scale *R*^2^ reported here is bounded by the VE series’ own measurement noise (Methods, *Reported measures of predictive accuracy*). The measurement-noise ceiling (Eq. S23) is 0.483 under leave-one-season-out, and the realized leave-one-season-out *R*^2^ (0.470) sits just below it: the residual sum of squares (0.368) is close to, and marginally above, the summed measurement variances of the published estimates (0.358). Thus, 0.47 is not to be read as “47% of variance explained”. This is the position a correctly specified model is expected to occupy – *R*^2^ at or below its ceiling – rather than a finding in itself. The evidence that pre-season geometry predicts effectiveness at all is the permutation test above (*P* = 0.003); the ceiling is reported to give *R*^2^ its scale. The practical implication is that observed-scale *R*^2^ in this project should always be read alongside its measurement-noise ceiling, since it establishes how much of the reported number even the true effectiveness signal could explain.

Two features of the VE series temper this reading. The mean measurement variance is inflated by five wide-CI seasons (2015–2016, 2019–2020, 2006–2007, 2013–2014, and 2024–2025; each 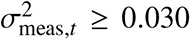 against a median of 0.013), two of which rest on a single network. And *σ*_meas,*t*_ inherits the precision cost of population weighting described below (*Collection and processing of influenza A(H3N2) vaccine effectiveness estimates*, *Population weighting versus meta-analytic pooling*): because population weights are fixed a priori rather than chosen to minimize variance, the composite standard error exceeds the inverse-variance-optimal pooled standard error, for the 14 of 16 seasons combining two or three networks (six with two, eight with three; 2006–2007 and 2015–2016 rest on a single network), by a median factor of 1.15 (range 1.00–1.75). Both features inflate the estimated measurement noise, which raises *σ*_meas,*t*_ and lowers the measurement-noise ceiling.

### Antigenic-map stability and leakage audit

#### Two leakage categories: response versus covariate construction

We distinguish two forms of information leakage (audited per variable in table S14). Response leakage (L1) occurs when a predictor uses *VE* (*t*) or a deterministic function of it; this is absent throughout, because every feature is a function of titers only. Covariate-construction leakage (L2) occurs when predictors at season *t* depend on the joint antigenic map, which was fit using titer data from seasons > *t*. L2 is real but unsupervised—*VE* never enters the map step—so its defense is empirical rather than theoretical: we test how much the map and the predictors built on it change when later titers are withheld.

#### Cross-season pre-season covariates do not violate leave-one-out

Pre-season covariates such as update drift projection 2y reference seasons *t*−2 and *t*−1 when predicting season *t*. Using the past to predict the future is the forecasting design, not a violation of it. In LOOCV, the held-out season’s *outcome* never enters the training loss—that is the only independence the procedure requires.

#### Expanding-window map-refitting protocol

For each cutoff season *t* in the expanding-window sequence, we (i) restricted the HI and PRNT panels to titers from seasons ≤ *t*, (ii) re-tuned the Topolow hyperparameters and refit the HI and PRNT maps, (iii) re-ran the joint Bayesian HI+PRNT unification (the four-observation-process latent-distance model) and refit the latent map at its per-fold robust optimum, (iv) carried the cluster labels over from the full-data map, (v) Procrustes-aligned the fold’s latent map to the consecutive fold’s map on the shared training strains, and (vi) recorded the residual sum of squares (the Procrustes *M*^2^). Step (iv) does not re-estimate the clusters: their construction rests on visual and voting checks that cannot be automated, and the labels are stable to the position shifts a fold introduces. Step (v) uses symmetric Procrustes analysis. To measure strain shifts in the frame in which the maps are displayed, every fold map was additionally aligned to the full-data map (rotation, reflection, translation and one scale factor) and projected onto the first two principal components of the full-data map, which carry 94.7% of its variance. We then recomputed every feature on each fold map and predicted season *t* + 1 with the three-predictor Bayesian model used throughout (vaccination coverage tm1, vaccine lead distance, and the oneseason update–drift projection), under the same priors and likelihood as the full-data fit. The fold feature tables differ in one respect: they carry no per-season antigenic standard deviation, so the fold fits set *S_ik_* = 0 and drop the errors-in-variables layer. The predictor comparison below covers the three map-derived predictors: the two in the model and the two-season update–drift projection.

#### Map positions are stable as seasons are added

Each additional season of titers left the positions of the strains already in the map largely unchanged (table S15; Fig. S18). The reference is the measurement-uncertainty ensemble: one latent map per posterior draw of the latent antigenic distances, so that two of its maps differ only through antigenic measurement uncertainty. Over the ten consecutive map pairs, the Procrustes residual ranged from *M*^2^ = 0.079 to 0.175 (Fig. S17). It fell below the median *M*^2^ between 300 pairs of reference maps on the same strains in every pair (ratio 0.55 to 0.86). In the expanding-window comparison of Fig. S18, the median shift of the prior strains was 0.53 to 0.78 antigenic units per added season, against 0.87 to 1.01 between pairs of reference maps on the same strains; the 95th percentile of the shift was 1.64 to 2.47 units. Adding a season of titers therefore moves the existing map less than antigenic measurement uncertainty does.

#### Predictor values at the prospective forecasts track their full-data values

For each of the nine prospectively scored target seasons (2015–2016 to 2024–2025, excluding 2020–2021, which has no VE estimate), we compared the value of each map-derived predictor used for that forecast, computed on the expanding-window map built from titers up to the preceding season, with its value for the same season on the full-data map (Fig. S19). The two were closely correlated: Pearson *r* = 0.96 for vaccine lead distance, 0.87 for update drift projection, and 0.91 for update drift projection 2y.

The median absolute difference was 0.17, 0.14, and 0.17 standard deviations of the respective predictor across seasons. vaccination coverage tm1 is not map-derived and is identical in both.

#### Static OLS is supported by the absence of serial correlation

At *N* = 16, autocorrelation tests are underpowered; we report them as a falsifiability check on the static specification, not as part of the leakage argument. No test detected serial dependence in log(OR): bootstrap Durbin–Watson = 2.13 (*p* = 0.79), Ljung–Box *Q*(1)-*Q*(3) (*p* = 0.59, 0.74, 0.76), Spearman *ρ*(1) = −0.12 (*p* = 0.66), and Lo–MacKinlay variance ratios VR(2)–VR(3) (*p* = 0.49, 0.42); the autocorrelation function is shown in Fig. S20. The defensible reading is “consistent with no serial dependence,” not “independence demonstrated.”

#### Prospective errors are consistent with held-out errors

Refitting the map and predicting the ten target seasons of the expanding window (2015–2016 through 2024–2025, nine with VE estimates) gave errors consistent with the held-out residual distribution that anchors the main predictive claim (table S16; Fig. S21). The prospective *R*^2^ across the nine predictions was 0.455, against 0.47 when only the outcome is held out; RMSE and MAE on the log(OR) scale were within 5% and 7% of their leave-one-season-out values (0.159 vs. 0.152; 0.130 vs. 0.122), and predicted and observed VE were rank-concordant (Spearman *ρ* = 0.767). Reassigning the nine observed values among the fixed forecasts in all 362,880 ways matched the forecasts at least as closely in 2,785 cases (exact *P* = 0.008). All nine prospective residuals fell within the range of the held-out residuals (Fig. S22). The gap in *R*^2^ is expected and does not indicate leakage in the main analysis: the prospective folds train on 7 to 15 seasons rather than 15, and *R*^2^ is computed over nine seasons whose own variance sets its denominator, so it is unstable at this size. The probabilistic comparison is more informative because it accumulates evidence from every forecast distribution rather than from nine point errors: scored against its own fold-training-mean forecast, the prospective CRPS was 0.084 against 0.130, a paired difference of 0.046, close to the paired difference of 0.040 between the leave-one-season-out CRPS (0.088) and its own historical-mean forecast (0.128); the two reference forecasts are not interchangeable, so each is reported as its own difference rather than a combined ratio or skill score (a ratio of two means has a small, noisy denominator at this number of seasons). The 95% predictive intervals covered all nine prospective seasons. The expanding-window results therefore reproduce the forecast quality that anchors the predictive claim, with the modest gap in RMSE and MAE reflecting the smaller training sets available in the earliest prospective folds rather than information leaking from the full-data map.

#### Quantifying predictive uncertainty: confidence intervals, prediction intervals, and the measurementnoise ceiling

We distinguish two uncertainty bands that answer different questions. A CI quantifies uncertainty in the *expected* VE for a given antigenic configuration—where the regression surface lies—and reflects estimation (coefficient) uncertainty only. A prediction interval (PI) quantifies uncertainty in a *single future observed*VE and is wider, because it adds the irreducible scatter of individual seasons about that surface. The two combine in variance, not in width: writing *σ*^2^ for the irreducible conditional variance of log(OR) and Var(*ŷ*_0_) for the estimation variance of the fitted value, Var(*y*_0_ − *ŷ*_0_) = Var(*ŷ*_0_) + σ_2_, so the half-widths satisfy 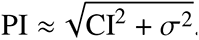. A prediction interval that approaches the irreducible scale therefore implies the estimation term is small *relative to σ* (not that it is zero).

#### Intervals for the held-out predictions (*Fig. 4A*)

For each held-out season the model returns a full predictive distribution, from which we report two intervals with distinct meanings. The credible interval on the expected value (table S13) reflects uncertainty in *α* and *β* alone and answers how precisely the season’s mean effectiveness is estimated. The wider predictive interval additionally carries the season-level residual *σ_u_* and that season’s measurement error *s_t_*, and is the interval a forecast of an observable VE estimate should be judged against; it is the one used for the coverage and calibration statistics above. Because VE = (1 − *e*^log(OR)^) × 100 is monotone decreasing in log(OR), interval bounds invert on back-transformation, VE_lo_ = (1 − *e*^log(OR)hi^) × 100.

#### Prediction intervals for the expanding-window forecasts (Fig. S21)

Each fold’s interval is the posterior predictive interval returned by the model fitted to that fold, so its width varies by season with the training size and that season’s measurement error, and it is not constructed from the residuals it is then assessed against. This is a material improvement over a flat band estimated from the nine prospective residuals, which would both assume constant predictive uncertainty and be evaluated on the same errors that set its width. The 95% intervals covered all nine target seasons. At *n* = 9 each season contributes about 11 percentage points of coverage, so we read this as a calibration check rather than a precise coverage estimate.

### Strain-substitution discrimination analysis

The candidate rankings (Fig. 4B) are meaningful only if the model separates candidates within a season by more than its own uncertainty; if it scored every candidate near-identically, the reported percentiles and gains would reflect numerical noise rather than antigenic separation. Of the model’s predictors, only two vary across candidates within a season—vaccine lead distance and update drift projection—and therefore do all of the within-season ranking; the remaining one is constant across candidates and shifts every candidate’s predicted VE by the same amount, leaving their order unchanged.

The posterior answers the separation question directly, by assigning each candidate the probability that it is the season’s most effective strain. That probability was distributed across many candidates: the smallest set carrying 95% of it held a median of 24.5 strains (range 12 to 51) out of pools of 57 to 376, and the leading candidate held a median probability of 0.25 (range 0.10 to 0.56) of being the optimum. The deployed vaccine entered that set in 5 of 16 seasons, and a median of 1 candidate per season (range 0 to 69) exceeded it with probability at least 0.95. The model thus produced a real spread of predicted effectiveness across the field (Fig. 4B) without sharply resolving which single strain was most effective. Across the 16 seasons the WHO-selected vaccine sat at a mean posterior rank percentile of 58.5.

### Support of the candidate rankings

The candidate that maximizes a fitted linear surface over a finite pool tends to fall near the edge of the predictor range, where the fit is least constrained by data. This is a property of the selection rule, not of the estimation method, and it is the principal limitation of the substitution analysis. We therefore report where each selected strain sits relative to the seasons the model was fitted on, measured by Mahalanobis distance from the centroid of the 16 training seasons in the three-dimensional predictor space.

In 6 of 16 seasons the model’s candidate lay outside the range of predictor values spanned by the training seasons; across the 16 selected strains the mean Mahalanobis distance from the training centroid was 2.49, below the maximum of 2.85 reached by the training seasons themselves. Across all candidates, 17.6% lay beyond that maximum. Predictions at such points are extrapolations, and the posterior widens there because the predictive variance grows with distance from the center of the design. That widening is reflected in the credible intervals reported throughout, although it does not solve the extrapolation issues completely.

Two features limit how far this matters. First, the tenth-percentile candidate, which maximizes the tenth percentile of the posterior rather than its mean and so penalizes uncertain predictions, coincided with the model’s candidate in 11 of 16 seasons and reduced the pooled gain from 10.4 pp to 10.0 pp; averaged over all 16 seasons it sat closer to the training region (mean Mahalanobis distance 1.98, against 2.49 for the model’s candidate). Second, the gain is a difference between two candidates in the same season, so the components of the extrapolation shared by both—including the season’s overall level—cancel. What does not cancel is the assumption that the fitted relationship between geometry and effectiveness continues to hold beyond the observed range, which these data cannot test. A mean function that reverts toward the prior away from the data, such as a Gaussian process, would encode this caution structurally, but its length-scales are not identifiable from 16 observations and it would express a prior assumption as though it were an inference.

### Collection and processing of influenza A(H3N2) vaccine effectiveness estimates

To quantify the degree to which seasonal influenza vaccines protect against circulating A(H3N2) viruses, we compiled published, adjusted, all-ages influenza A(H3N2)-specific VE estimates from three major NH sentinel surveillance networks spanning the 2004–05 through 2025–26 influenza seasons (table S9). VE served as a season-level outcome variable in our regression models linking antigenic evolution of H3N2 viruses to observed vaccine performance. This section describes the source networks, the criteria for selecting and excluding individual estimates, the procedure for aggregating network-specific estimates into a single NH composite, the transformation applied before regression modeling, and the limitations of this compilation.

#### Source surveillance networks

Estimates were drawn from three ongoing, population-based sentinel surveillance networks, each employing the TND (*50, 62*):

1. **US Flu VE Network / CDC** (United States). A multi-site outpatient surveillance network coordinated by the Centers for Disease Control and Prevention (CDC), enrolling patients with acute respiratory illness (ARI) at ambulatory care facilities across 4–5 US sites. VE is estimated by comparing the odds of influenza vaccination among RT-PCR-confirmed influenza cases versus test-negative controls, adjusting for age, calendar time, site, highrisk medical conditions, and specimen collection interval (*120–122*). For seasons 2004–05 through 2023–24, estimates are drawn from end-of-season publications by the US Flu VE Network. For 2024–25 and 2025–26, only interim estimates were available (see below).
2. **Canadian Sentinel Practitioner Surveillance Network (SPSN)**. A community-based primary care sentinel network coordinated by the British Columbia center for Disease Control, enrolling patients presenting with influenza-like illness (ILI) at sentinel practitioner sites across four Canadian provinces (British Columbia, Alberta, Ontario, and Quebec). VE is estimated via TND with adjustment for age, province, comorbidity, specimen collection interval, and calendar time modeled with spline functions (*123–125*). Estimates are available from the 2004–05 season onward, though not all seasons produced H3N2-specific estimates.
3. **I-MOVE / VEBIS (Europe)**. The Influenza Monitoring Vaccine Effectiveness in Europe (IMOVE) and, from 2021–22, the Vaccine Effectiveness, Burden, and Impact Studies (VEBIS) multicenter primary care case–control study, coordinated by Epiconcept (Paris) in collaboration with the European center for Disease Prevention and Control (ECDC). The network pools individual patient data from 8–11 study sites across EU/EEA member states using a one-stage logistic regression model with study site as a fixed effect, adjusting for age, sex, chronic conditions, and symptom onset date (*126–128*). H3N2-specific European VE estimates are available from the 2011–12 season onward.

These three networks were selected because they cover the major NH populations (United States, Canada, and Europe) for which consistent, long-running surveillance data exist, and because each estimates subtype-specific VE against medically attended, laboratory-confirmed influenza using the TND—the current standard for observational VE estimation (*33*)—enabling a standardized compilation.

#### Estimate selection and exclusions

For each network–season we extracted a single adjusted, all-ages, primary care/outpatient, H3N2-specific end-of-season VE point estimate and its 95% CI wherever available; hospital-based estimates were excluded because they measure a different clinical endpoint (prevention of hospitalization rather than prevention of medically attended illness) not directly comparable to outpatient VE (*129*). Seasons with no estimable H3N2-specific VE from any network were excluded, and individual networks contributed no estimate in seasons of insufficient H3N2 circulation (e.g., 15 A(H3N2) cases in the SPSN in 2013–14 (*130*); 29 in the US Flu VE Network in 2019–20 (*131*)).

#### Aggregation into a NH composite VE

All three networks operate in the NH and therefore evaluate the same WHO-recommended NH vaccine composition; restricting the composite to NH networks is consistent with evidence that influenza VE differs systematically between hemispheres (*27*). Because the three networks cover distinct populations of different sizes, we computed a population-weighted average VE across available networks for each season. Population weights were based on approximate mid-period (circa 2015) national population sizes:

- United States (CDC): 330 million
- European Union (I-MOVE/VEBIS): 450 million
- Canada (SPSN): 38 million

For a season in which all three networks reported an estimate, the composite VE was calculated as:

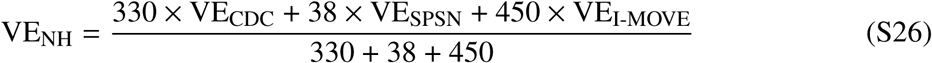

When one or more networks lacked an estimate (coded as NA), the weights were re-normalized over the available networks. For example, if only CDC and SPSN reported estimates:

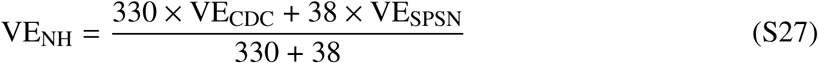

The composite’s sampling variance is derived by weighted-averaging of confidence limits. Writing *a_i_* for network *i*’s normalized population weight and *s_i_* for its own log(OR)-scale standard error (from its published CI), 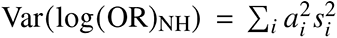, the variance of a fixed-weight linear combination of independent network estimates; the composite 95% CI is the back-transform of the log(OR)-scale point estimate ± *z*_0.975_ times the resulting standard error. Note that formal metaanalytic pooling (e.g., inverse-variance weighting) targets a different estimand – the single common mean of an assumed shared true effect – and was not used for the point estimate because the objective here is a summary of the NH population’s realized VE experience, a fixed-weight average of the networks’ own effects (*Basis for pooling across networks*, below). Cochran’s *Q* and *I*^2^, computed with the standard inverse-variance weights, are nonetheless reported per season as a diagnostic of whether the networks are consistent with a common season-level effect.

#### Basis for pooling across networks

Three considerations support combining the network estimates into a single NH outcome. All three networks estimate VE with the same test-negative design, the standard for observational VE (*33, 50, 62*). Pooling subtype-specific VE across surveillance locations is established practice: the largest H3N2 meta-analysis to date combined 56 test-negative studies spanning five continents and both hemispheres into a single random-effects estimate (*2*). And the three regions are not antigenically independent: global A(H3N2) circulates as a temporally structured metapopulation in which lineages migrate rapidly among Northern-Hemisphere temperate regions within each season, with no region maintaining a persistent local source (*132*). A given NH season therefore presents a largely shared antigenic variant to the US, Canadian, and European populations, all immunized against the same WHO-recommended NH composition.

These expectations are borne out where the networks overlap (Fig. S11): pairwise rank correlations of season-level VE ranged from Spearman *ρ* = 0.36 to 0.83 (11 jointly reported seasons), and a random-effects decomposition of per-network VE attributed 64% of the variance to differences between seasons and only 4% to differences between networks. The population-weighted NH composite therefore reflects a shared, season-driven antigenic exposure rather than an average over disparate regional regimes.

A season-level heterogeneity test points the same way. We computed Cochran’s *Q* per season with the standard inverse-variance weights, across the 15 seasons with two or three reporting networks. It was non-significant in every season (minimum *P* = 0.051, 2025–2026). *I*^2^ had a median of 0% and was exactly 0% in 9 of the 15, although, neither statistic, on its own, establishes homogeneity: with *k* ≤ 3 networks per season. Read together with the variance decomposition, the heterogeneity test therefore leaves the composite untroubled: the networks show no evidence of disagreement, and the fixed-weight average would stand even if they did.

#### Transformation for regression modeling

Influenza VE estimated via TND is defined as VE = (1 − OR) × 100, where OR is the adjusted odds ratio for influenza infection among vaccinated versus unvaccinated persons from logistic regression (*33*). For regression modeling, VE was backtransformed to the log odds ratio scale:

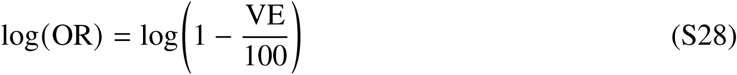

This transformation has three desirable properties for linear modeling: (i) it places the outcome on the natural scale of the logistic regression models from which VE was originally estimated; (ii) it avoids the heteroscedastic residuals that VE’s bounded range (constrained between −∞ and 100%) can produce in linear models; and (iii) it maps VE = 0% (no protection) to log(OR) = 0, providing a natural null reference. A higher (more positive) log(OR) corresponds to lower vaccine protection (i.e., OR closer to 1 or above), while a more negative log(OR) corresponds to higher VE.

#### Compiled VE estimates

Table S9 presents the complete compilation of H3N2-specific VE estimates by network and season, along with the population-weighted NH composite used in subsequent analyses. The 2025–26 season is included for completeness but was not part of the regression analysis sample (*N* = 16).

#### Per-season data sources

The published source for each network–season VE estimate in Table S9 is as follows. **CDC/US Flu VE Network:** 2004–05 (*120*); 2007–08 (*133*); 2010–11 (*134*); 2011–12 (*51*); 2012–13 (*135*); 2013–14 (*136*); 2014–15 (*137*); 2015–16 (*121*); 2016–17 (*122*); 2017–18 (*138*); 2018–19 (*26*); 2021–22 (*139*); 2023–24 (*140*); 2024–25 (*141*); 2025–26 (*142*). **SPSN (Canada):** 2006–07 (*123*); 2007–08 (*143*); 2010–11 (*144*); 2012–13 (*145*); 2014–15 (*124*); 2016–17 and 2017–18 (*125*); 2018–19 (*17*); 2019–20 (*30*); 2021–22 (*146*); 2022–23 (*147*); 2023–24 (*148*); 2024–25 (*149*); 2025–26 (*25*). **I-MOVE/VEBIS (Europe):** 2011–12 (*126*); 2012–13 (*150*); 2013– 14 (*151*); 2014–15 (*152*); 2016–17 and 2017–18 (*127*); 2018–19 (*153*); 2019–20 (*154*); 2021– 22 (*155*); 2022–23 (*128*); 2023–24 (*156*); 2024–25 (*157*); 2025–26 (*158*).

#### Limitations and caveats

Several methodological considerations should be noted when interpreting the compiled VE data.

##### Heterogeneity across networks

Notwithstanding this concordance, and although all three networks use the TND, they differ in participant recruitment (practitioner-based vs. multi-site enrollment), covariate adjustment strategies (categorical age groups vs. spline-modeled continuous age and calendar time), population age structure and vaccination coverage, circulating viral cluster composition, and vaccine products administered (e.g., differing proportions of egg-based vs. cell-based or adjuvanted vaccines). These differences may contribute to inter-network variation in VE estimates within the same season (*124*).

##### Population weighting versus meta-analytic pooling

Our population-weighted averaging treats each network’s point estimate equally per capita. This approach does not account for differences in statistical precision (sample size, number of H3N2 cases) across networks. In seasons where one network’s estimate has wide CIs (e.g., SPSN 2021–22: 36%; 95% CI: −38 to 71), its contribution to the point estimate is determined solely by population size, not by the informativeness of the estimate.

##### Minor comparability caveats in recent seasons

Two features of the recent US estimates slightly reduce their comparability with earlier seasons. First, the 2024–25 and 2025–26 CDC values are adult (aged ≥18 years) rather than all-ages outpatient H3N2-specific estimates. This shift is minor: in both seasons the network’s children and adult H3N2 estimates were close and non-significant, so using the adult stratum in place of an all-ages estimate changes the value only modestly, and only 2024–25 enters the *N* = 16 regression sample. Second, both seasons rely on interim rather than end-of-season estimates.

##### Interim vs. end-of-season estimates

The 2024–25 and 2025–26 estimates are interim (midseason) rather than end-of-season. Interim estimates may differ from final estimates because (i) additional cases accrue in the later part of the season, potentially with different circulating cluster distributions; (ii) waning VE over time may reduce end-of-season estimates relative to mid-season; and (iii) interim analyses typically have smaller sample sizes and wider CIs (*127, 159*).

##### Egg-adaptation and vaccine type

An important source of variation not captured in this compilation is the proportion of participants receiving egg-based vs. cell-based or recombinant vaccines, which differs across networks and has changed over time. Egg-adaptation during vaccine manufacturing introduces mutations at key antigenic sites (notably T160K in the HA glycoprotein of H3N2), which can substantially reduce immunogenicity against circulating wild-type viruses (*53,145,160*).

##### Exclusion of the SH

The VE models are restricted to NH seasons for three data reasons, none reflecting an assumption that the model is hemisphere-specific. First, SH antigenic sampling is too sparse to support reliable predictors: of the 2,053 characterized antigen strains on the map, only 180 are SH against 1,873 NH, a median of roughly 5 versus 55 strains per season (as few as one in some SH seasons). The season centroid, diversity, cluster-share, and stretch measures that drive our models cannot be estimated stably from so few isolates. Second, although SH H3N2 VE estimates are published—most consistently from Australian sentinel networks (*161*)—no continuous, multinetwork, subtype-specific SH series comparable to our NH composite was available; the few heterogeneous, intermittent season estimates are too sparse to constitute a robust out-of-sample test. Third, the modeled vaccination-coverage covariate is a population-weighted average of United States, Canadian, and European coverage, and we identified no large, consistent SH analog to substitute. Where SH antigenic dynamics can be characterized they are concordant with the NH series (Fig. 1C; Fig. 2B); because influenza VE also differs systematically between hemispheres (as noted above), transferring an NH-trained model to SH seasons would require explicit recalibration rather than serving as direct validation.

##### Source terms and attribution

Each value is an individual published point estimate with its 95% CI, transcribed from the network report cited for that season above (*Per-season data sources*). The two Unite

States values taken from the *MMWR* are works of the United States federal government in the public domain; their use here does not imply CDC endorsement. Estimates drawn from *Eurosurveillance*, *PLoS ONE*, *Vaccine X*, and *Influenza and Other Respiratory Viruses* are reused from open-access articles under their published licences, with attribution to the cited article. The remaining estimates are reproduced as individual published figures with citation.

##### Data availability

The compiled VE estimates are provided in Data S1 as figS11 h3n2 ve by network seaso whose cdc source, spsn source, and imove source columns carry the source citation for each network–season value; the per-season composite and its interval are in fig5 season master clean.csv.

The R code for computing population-weighted composites and transforming VE to log(OR) is provided in (*61*).

#### Collection and processing of vaccination coverage data

Prior-season vaccination coverage entered the models as a population covariate (vaccination coverage t and its one-season lag vaccination coverage tm1; Table 1). Because no all-ages, hemisphere-wide coverage figure exists, we assembled a single NH coverage series from the same three regions that define the VE composite—the United States, the European Union, and Canada—for the 2003–04 through 2024–25 seasons (table S17). This section describes the source systems, the harmonization of age denominators across regions, and the aggregation into the modeled covariate. *Source data systems.* United States coverage was taken from the CDC FluVaxView interactive dashboard, which reports end-of-season influenza vaccination coverage drawn from the National Health Interview Survey (NHIS) and the Behavioral Risk Factor Surveillance System (BRFSS) (*162–164*); for seasons before 2009–10, for which FluVaxView produces no series, we used published NHIS/BRFSS estimates (*163, 165–168*). European coverage was taken from the Eurostat EU-wide vaccination rate for people aged ≥65 years (online data code hlth ps immu), which provides the broadest single all-EU Figure (*169*); seasons not covered by Eurostat were filled with the ECDC/VENICE cross-country median coverage rate (*170–175*), and the earliest seasons (2001–02 to 2005–06) with a five-country household survey (*176*). Canadian coverage was taken from the Statistics Canada Canadian Community Health Survey (CCHS), which reports self-reported coverage for persons aged ≥12 years (*177–180*); the Public Health Agency of Canada (PHAC) Seasonal Influenza Vaccination Coverage Survey (*181*) was used only for seasons the CCHS did not cover, because it tends to report systematically higher coverage.

#### Metric selection and harmonization

For the United States we used the end-of-season maximum of the adult (≥18 years) coverage estimate as the primary series, retaining the ≥65-year series for the harmonization steps below. This adult basis—rather than the all-ages (≥6 months) figure that the CDC dashboard headlines—was chosen for cross-region comparability, because the European and Canadian systems do not report an all-ages figure (Europe reports only ≥65 years; the CCHS reports ≥12 years). To place all three regions on a common adult footing we applied two age-bridging steps. First, because the European series is reported only for ≥65 years, we inferred a European adult coverage rate as 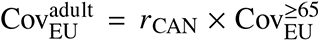, where *r*_CAN_ ≈ 0.47 is the mean ratio of Canadian general (≥12) to elderly (≥65) coverage over the pre-2014–15 seasons—the period before Canada’s universal vaccination recommendation and therefore the interval most comparable to the European programmes, which target older and at-risk groups. Second, for the pre-2009 United States seasons that lack a published adult estimate, we inferred adult coverage by scaling the ≥65-year value by the adult-to-elderly ratio observed in the earliest FluVaxView season with both strata (≈ 0.58). All regional values are listed in table S17.

#### Aggregation into a NH composite

Following the same population weighting used for the VE composite, we combined the three regional adult/general series into a single NH coverage rate using approximate mid-period (circa 2015) national population sizes as weights (United States 330 million, European Union 450 million, Canada 38 million):

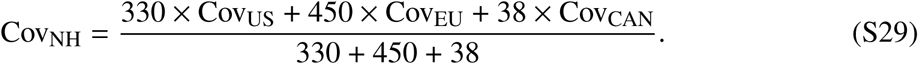

When a region lacked a coverage value in a given season, the weights were re-normalized over the available regions, exactly as for the VE composite. This NH rate is the modeled vaccination coverage t; its one-season lag vaccination coverage tm1 is the pre-season predictor retained in the inferential and predictive models.

#### Limitations and caveats

Several features of this compilation warrant caution. All three systems rely on self-reported vaccination status from population surveys, which tends to overestimate true coverage. The regional series use different age denominators (≥18, ≥65, and ≥12 years), which the ratio-based bridging only approximately reconciles; in particular, the European adult series is inferred rather than measured, and its level depends on the assumed Canadian ratio. Eurostat reports on a calendar-year basis, which we mapped to influenza seasons, whereas the other sources report by season. None of the sources report H3N2-specific coverage; all figures are for the seasonal vaccine, which includes an H3N2 component. Finally, the unweighted ECDC/VENICE cross-country median and the population-weighted Eurostat average can diverge, so the few VENICE-filled seasons are not strictly on the same footing as the Eurostat-based majority.

#### Source terms and attribution

The United States figures are drawn from material developed by the CDC, a work of the United States federal government in the public domain; their use here does not imply CDC endorsement. The European figures reuse Eurostat and ECDC material under the Creative Commons Attribution 4.0 International licence (https://creativecommons.org/ licenses/by/4.0/); ECDC is acknowledged as the creator of the ECDC/VENICE coverage rates, and both European series were modified by the calendar-year-to-season mapping and the age-bridging step described above. The Canadian CCHS figures are adapted from Statistics Canada, *Health characteristics, annual estimates* (table 13-10-0096) (*179*), which does not constitute an endorsement by Statistics Canada of this work; the 2022–23 Canadian value contains information licensed under the Open Government Licence–Canada (*181*). Coverage figures taken from the cited journal and report literature are reproduced as individual published estimates.

#### Data availability

The regional coverage inputs are provided in Influenza Vaccination Coverage for All (US FluVaxView export), us coverage pre09.csv, eu coverage elderly data.csv, and can coverage da the merged per-season series is provided in Data S1 as tableS17 global coverage comparison.csv.

The R code that filters, harmonizes, and population-weights these series into vaccination coverage t is provided in (*61*).

---

**Figure S1:**
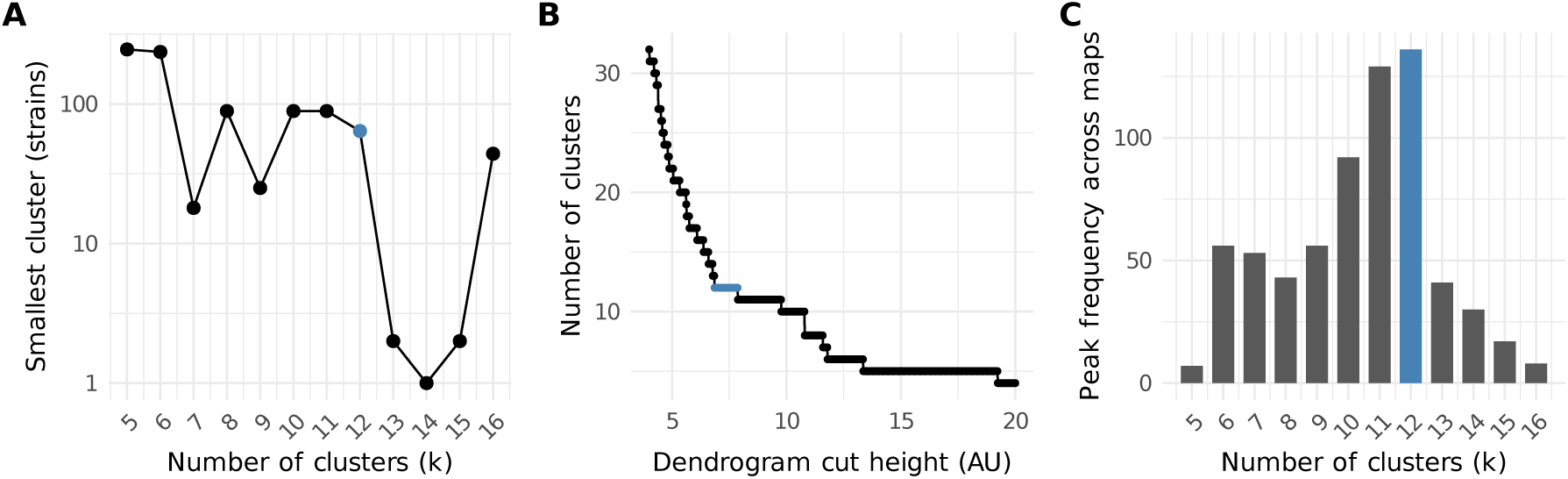
Choosing the number of antigenic clusters. We evaluated three diagnostics over *k* = 5–16; none presupposes compact or convex clusters. Panels **B** and **C** are the two views of the single hierarchy-level criterion described in the text. The adopted value, *k* = 12, is marked in blue. (**A**) Size of the smallest cluster in the evidence-accumulation consensus partition, rebuilt at each *k* (log scale). Twelve is the largest count at which no cluster falls below 50 strains (the smallest holds 64); at *k* = 13 the smallest drops to 2, and at *k* = 14 to 1. (**B**) Number of clusters versus Ward.D2 dendrogram cut height. A count that corresponds to a genuine level of the hierarchy persists over an interval of cut heights; twelve clusters persist over 1.43 AU, eleven over 0.05 AU. (**C**) Number of the 200 independent map generations in which a given count carries a prominent local maximum of the dendrogram height difference. The votes are spread across the mid-range, peak at *k* = 12, and drop by roughly two-thirds at *k* = 13 to single figures by *k* = 16.

**Figure S2:**
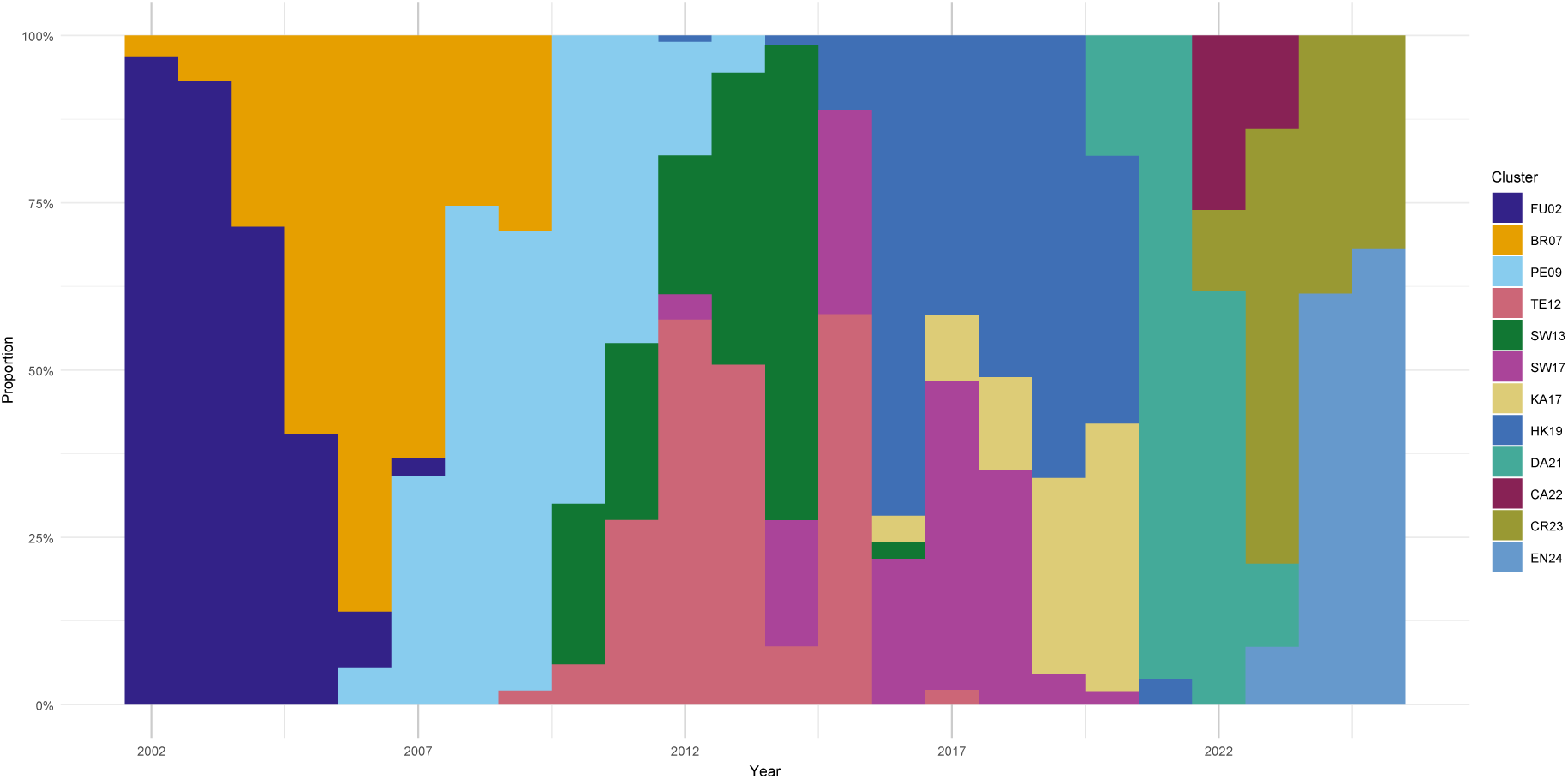
Antigenic cluster membership of NH H3N2 over time. Normalized stacked frequencies of the twelve antigenic clusters by year (each year sums to 100%). Successive clusters rise and fall in chronological order, but transitions occur through multi-year overlap zones in which two or more clusters co-circulate, rather than through instantaneous replacement.

**Figure S3:**
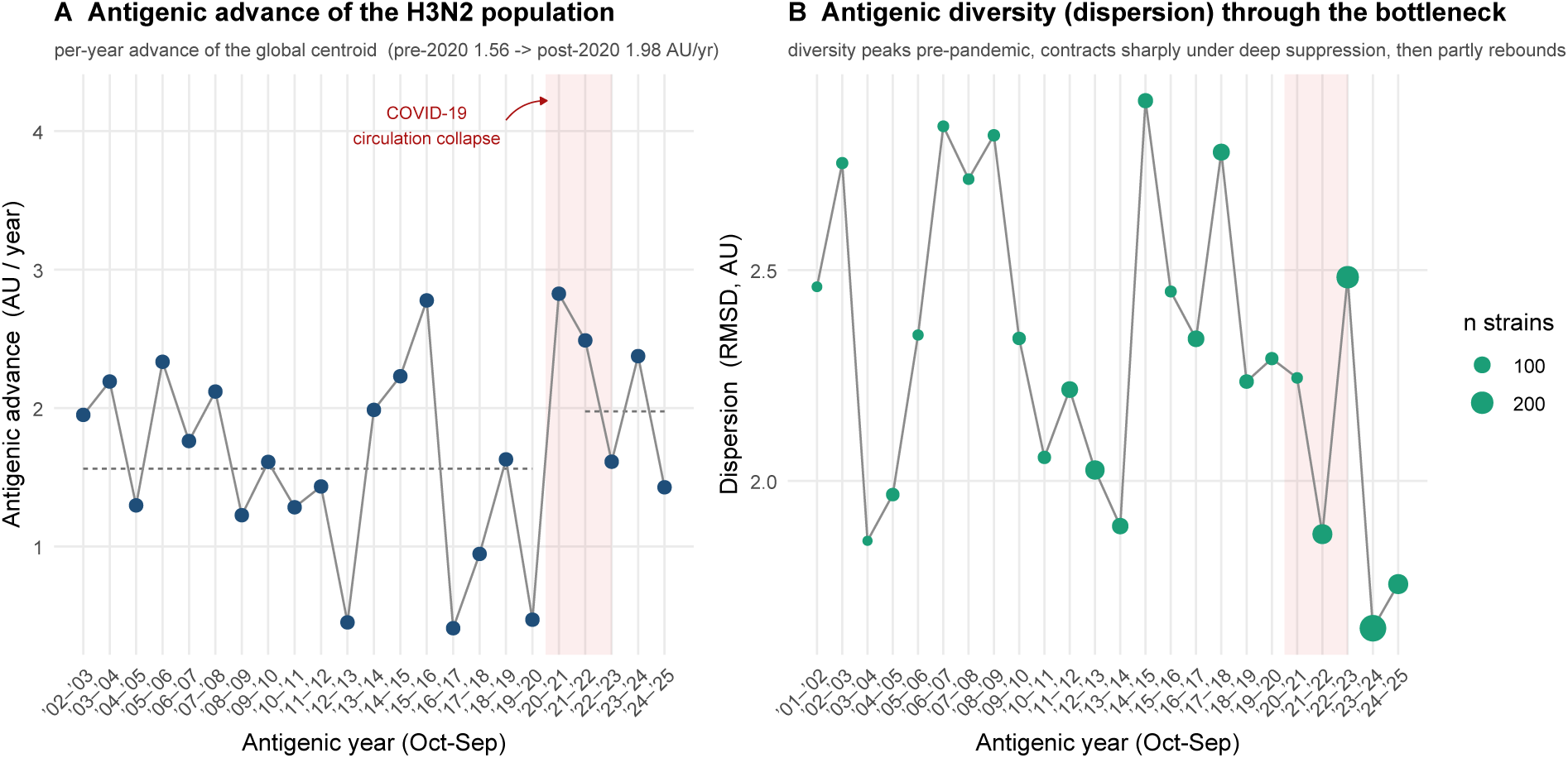
Antigenic advance and diversity of the H3N2 population across the COVID-19 pandemic. **A**) Per-year antigenic advance of the global population centroid—the displacement between consecutive yearly centroids (the first difference of the cumulative-distance staircase in Fig. 1). Both hemispheres are pooled, because antigenic advance is a property of the single globally circulating population rather than of a hemisphere’s vaccine programme. Every virus is binned by its collection date into a common October– September antigenic year. Each point is plotted at the landing year, so its height is the antigenic advance *into* that year from the previous one. Dashed segments mark the mean rate over the landings up to 2019– 2020 and over those from 2021–2022 onward (1.56 and 1.98 AU/year). Both windows differ from the ones the main text scales against: its pre-pandemic baseline starts at the 2005–2006 landing (1.51 AU/year) and its post-pandemic mean covers 2022–2023 through 2024–2025 only (1.80 AU/year), excluding the second collapse-era landing. The shaded band marks the collapse-era 2020–2021 and 2021–2022 landing years. Both collapse-era landings show large advances (2.83 and 2.49 AU, the largest and third largest of the record, with the 2.78 AU into 2015–2016 between them), and a further 2.38 AU advance followed into 2023–2024, after the resurgence of H3N2 circulation (*182*). (**B**) Antigenic diversity of the circulating population, the root-mean-square distance (dispersion, AU) of each year’s characterized strains from their yearly centroid, on the same global antigenic years as (**A**); points sized by strain count. Diversity contracted sharply in the deeply suppressed 2021–2022 year (1.87 AU) and rebounded in 2022–2023 (2.48 AU), before falling to the two lowest values of the record in 2023–2024 (1.65 AU) and 2024–2025 (1.75 AU), a contraction and recovery distinct from the advance in (**A**), as expected when a population passes through a circulation bottleneck. Characterization thinned at the collapse itself (38 strains in 2020–2021, compared with 26 to 297 across the record) and recovered in 2021–2022 (152 strains), so the 2020–2021 reading rests on the sparser of the two panels.

**Figure S4:**
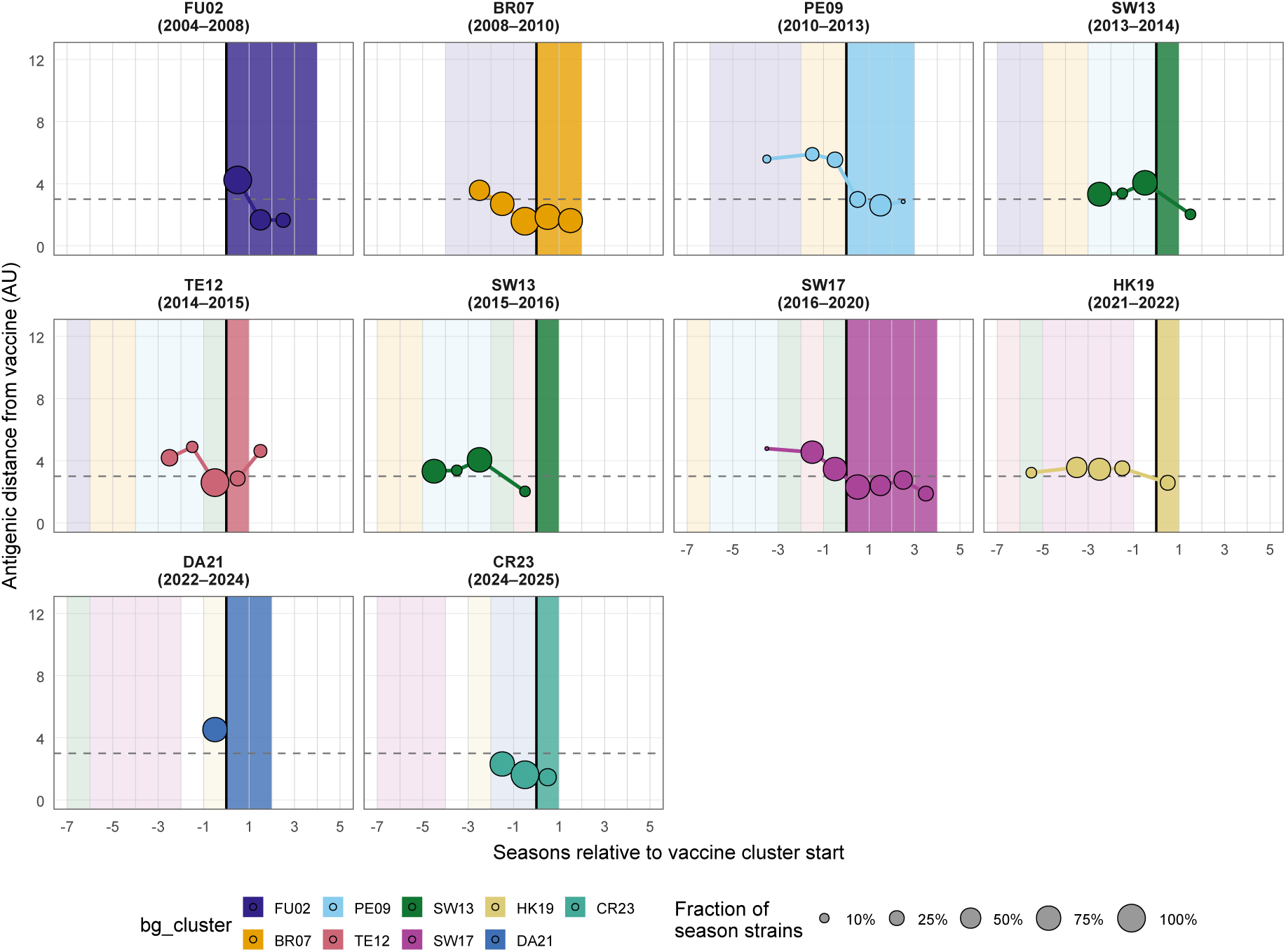
Cluster-aligned antigenic distance from the vaccine (SH). As in Fig. S5, but for SH vaccine clusters: each panel follows one SH antigenic cluster, re-centred so that season 0 (black vertical line) is the first season a vaccine from that cluster was recommended, with the dark band marking the cluster-match seasons, faint colored bands to its left the earlier clusters whose vaccines were standing beforehand, and unshaded seasons to the right those following the vaccine’s update to a later cluster. Bubbles show the mean antigenic distance of the cluster’s circulating strains from that season’s WHO-recommended vaccine (AU), sized by the cluster’s frequency (fraction of that season’s characterized strains); the dashed line marks 3 AU, the eight-fold reduction in HI titer at which a circulating virus is conventionally judged antigenically distinct from the vaccine reference strain (*25, 26*).

**Figure S5:**
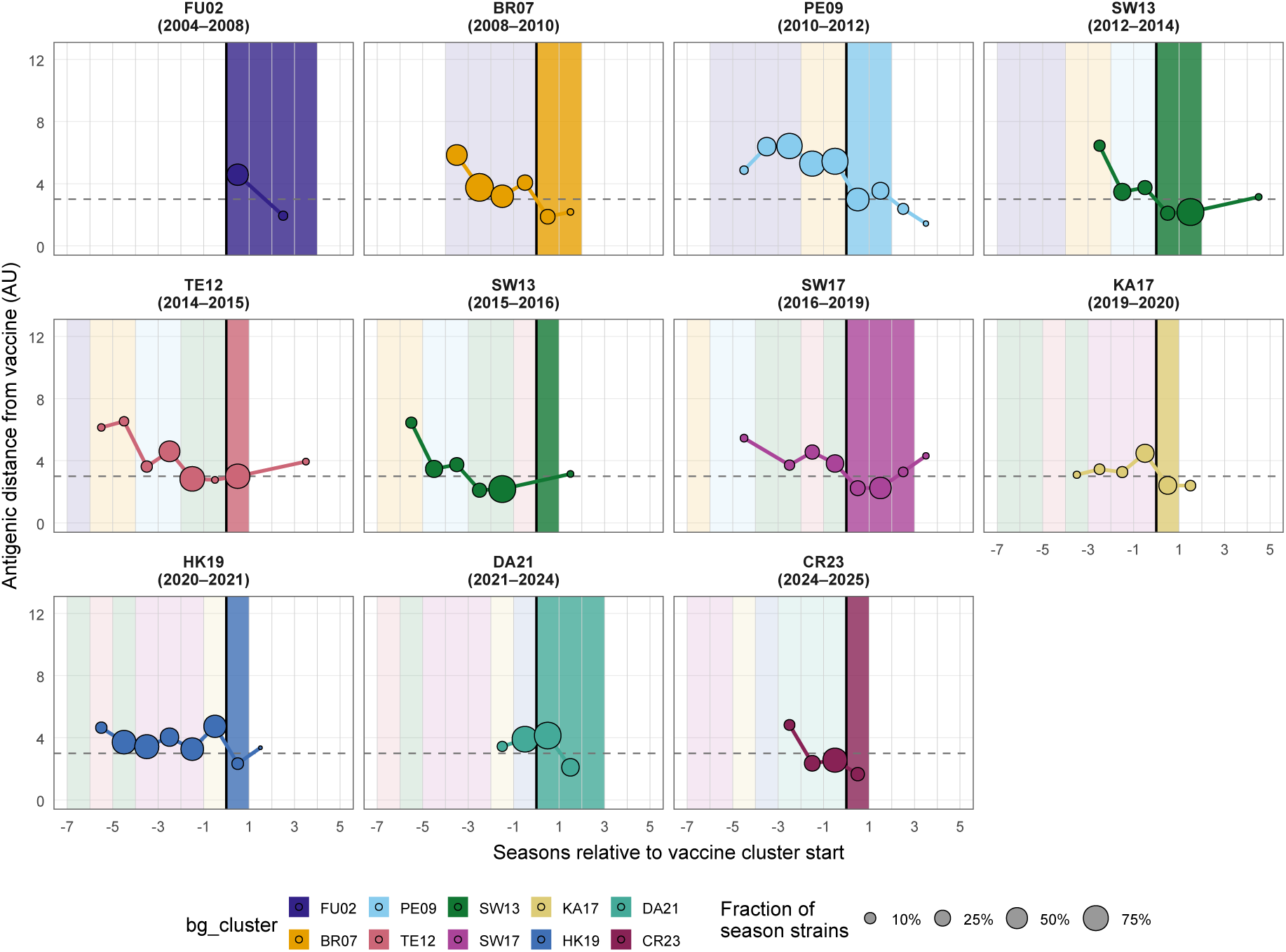
Cluster-aligned antigenic distance from the vaccine. Each panel follows one NH antigenic cluster, re-centred so that season 0 (black vertical line) is the first season a vaccine from that cluster was recommended; the strip label gives the cluster name and the calendar-year span of its vaccine era. Bubbles show the mean antigenic distance of the cluster’s circulating strains from that season’s WHO-recommended vaccine (vertical axis, AU), sized by the cluster’s frequency (fraction of that season’s characterized strains), with the line tracing this distance across seasons. The dark band marks the cluster-match seasons, when the standing vaccine belonged to the panel’s own cluster; faint colored bands to its left are the earlier vaccines’ clusters, and unshaded seasons to the right fall after the vaccine had been updated to a later cluster. Distances are therefore measured from the previous vaccines before the band, from the cluster’s own vaccine within it, and from later vaccines after it. The dashed line marks 3 AU, the eight-fold reduction in HI titer at which a circulating virus is conventionally judged antigenically distinct from the vaccine reference strain (*25,26*). For several clusters the defining strains circulated 6–8 AU from the contemporaneous vaccine for one to three seasons before the vaccine was updated to them, after which the distance typically fell below 3 AU.

**Figure S6:**
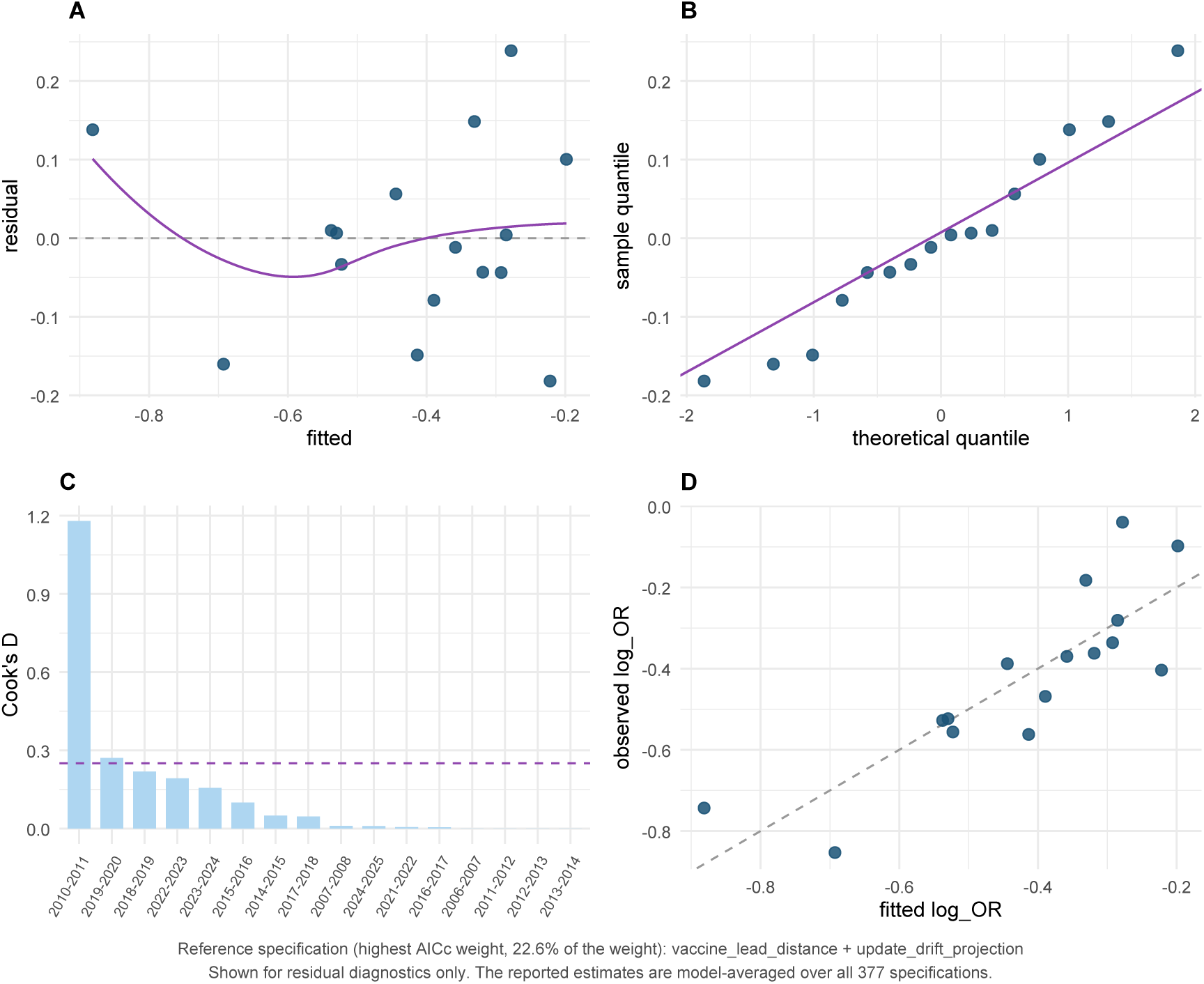
Residual diagnostics for the best-supported specification. Shown for the single specification carrying the largest AICc weight (prior-season lead distance and the one-season update–drift projection; 22.6% of the weight, *N* = 16). It is displayed because residual diagnostics require a fitted model, and it is *not* the reported estimate: every coefficient in the paper is averaged over the 377 specifications, and assumption satisfaction across that whole space is summarized in the Supplementary Text, *Inferential model: detailed results*. (**A**) Residuals versus fitted values. (**B**) Normal Q-Q plot. (**C**) Cook’s distance by season, with the 4 *n* = 0.25 threshold dashed. (**D**) Observed against fitted log OR . This specification satisfies the standard assumptions: Shapiro–Wilk *W* = 0.967, *p* = 0.79; Breusch–Pagan *χ*^2^ = 0.64, *p* = 0.73; Durbin–Watson = 2.14; two seasons exceed Cook’s *D* > 0.25.

**Figure S7:**
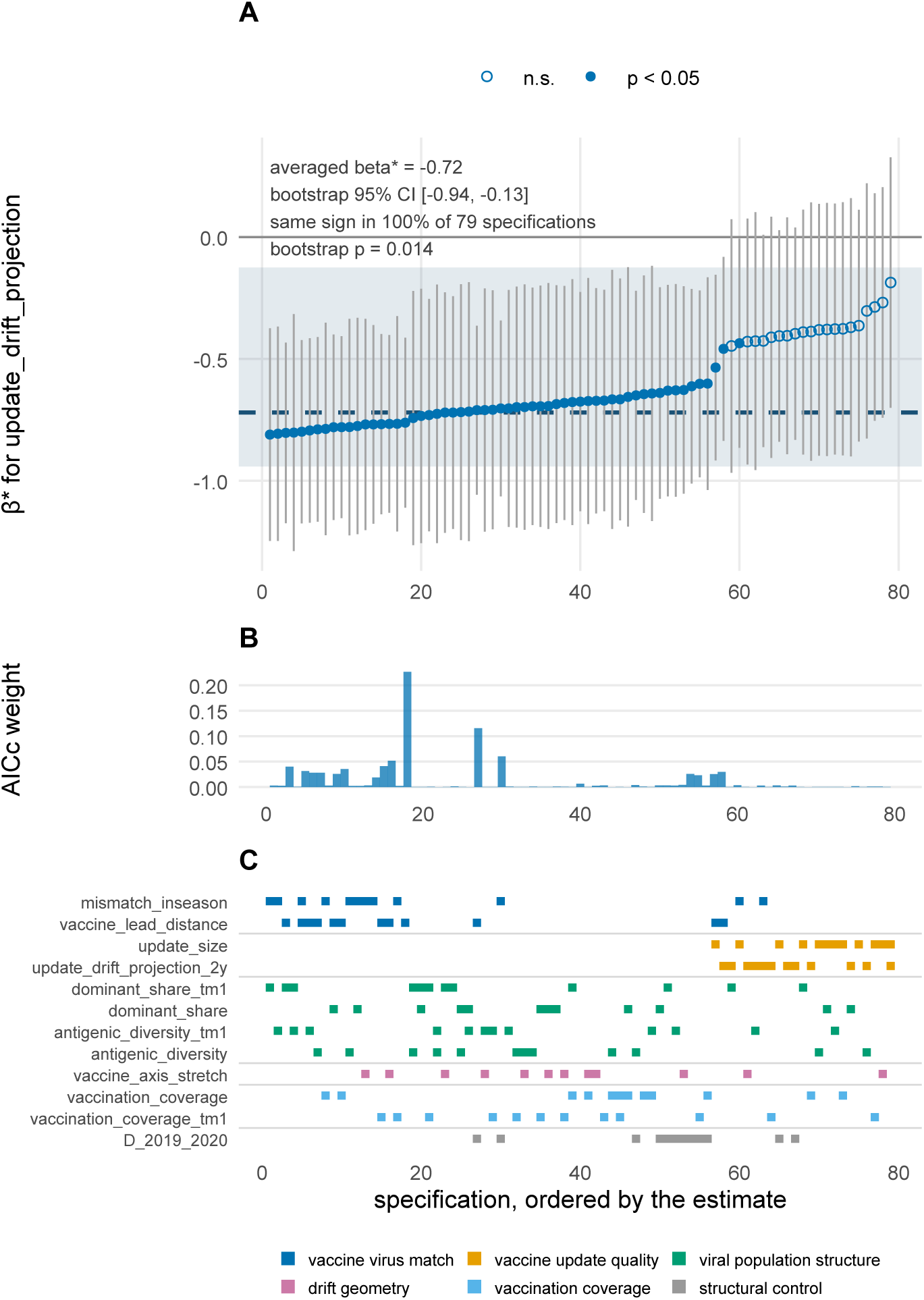
Specification curve for the one-season update–drift projection. Each of the 377 specifications of at most three predictors is fitted, and the 79 containing this variable are shown. (**A**) Estimated *β̄*^∗^ in each of the 79, sorted, each with that specification’s own 95% confidence interval; filled symbols mark specifications in which the variable is individually significant at *p* < 0.05. The dashed line is the model-averaged estimate and the shaded band its 95% bootstrap interval. Every estimate is negative and the bootstrap interval excludes zero, but the interval is wide: the estimate is robust to what is controlled for and imprecise because there are 16 seasons. (**B**) The AICc weight of each specification in the same order, which shows where the support sits along the curve rather than only where the estimates lie. (**C**) Which other variables enter each specification, colored by mechanistic category. Restricting to the 48 specifications carrying at most one indicator per category moves this variable’s average by only 0.02, so within-category competition does not drive its spread. Curves for all 13 candidates, including the three whose distributions are not unimodal, are in the accompanying multi-page file.

**Figure S8:**
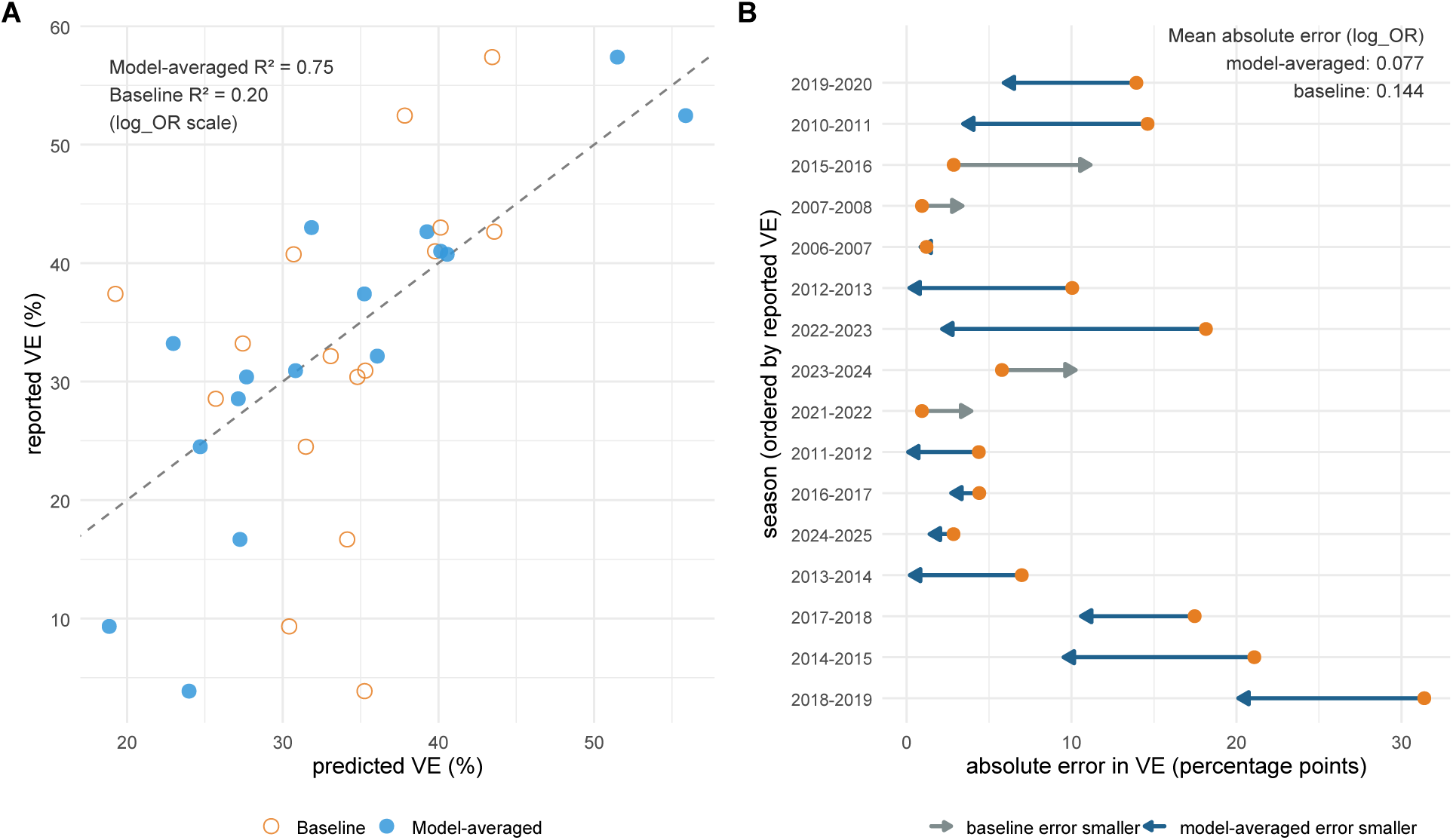
Explanatory performance of the model-averaged prediction versus a scalar-distance baseline. The geometry side is the AICc-weighted model-averaged prediction over all 377 specifications, *ŷ* = Σ*_m_ w_m_ŷ _m_*, which requires no selection step; the baseline is an ordinary least squares fit on the two scalar vaccine–virus distances alone. Both are in-sample, the appropriate comparison for an explanatory model (*114*). (**A**) Reported against predicted VE, one point per season, with the dashed line marking perfect fit. The averaged prediction reaches *R*^2^ = 0.75 against *R*^2^ = 0.20 for the baseline. (**B**) Change in absolute error by season, seasons ordered by reported VE. Each arrow runs from the baseline error to the model-averaged error, so an arrow pointing left is a season the geometry improves; the averaged prediction has the smaller error in 12 of the 16 seasons. Axes are on the VE scale for readability; the annotated *R*^2^ and mean absolute error are on the log(OR) scale, on which both models are fitted and evaluated.

**Figure S9:**
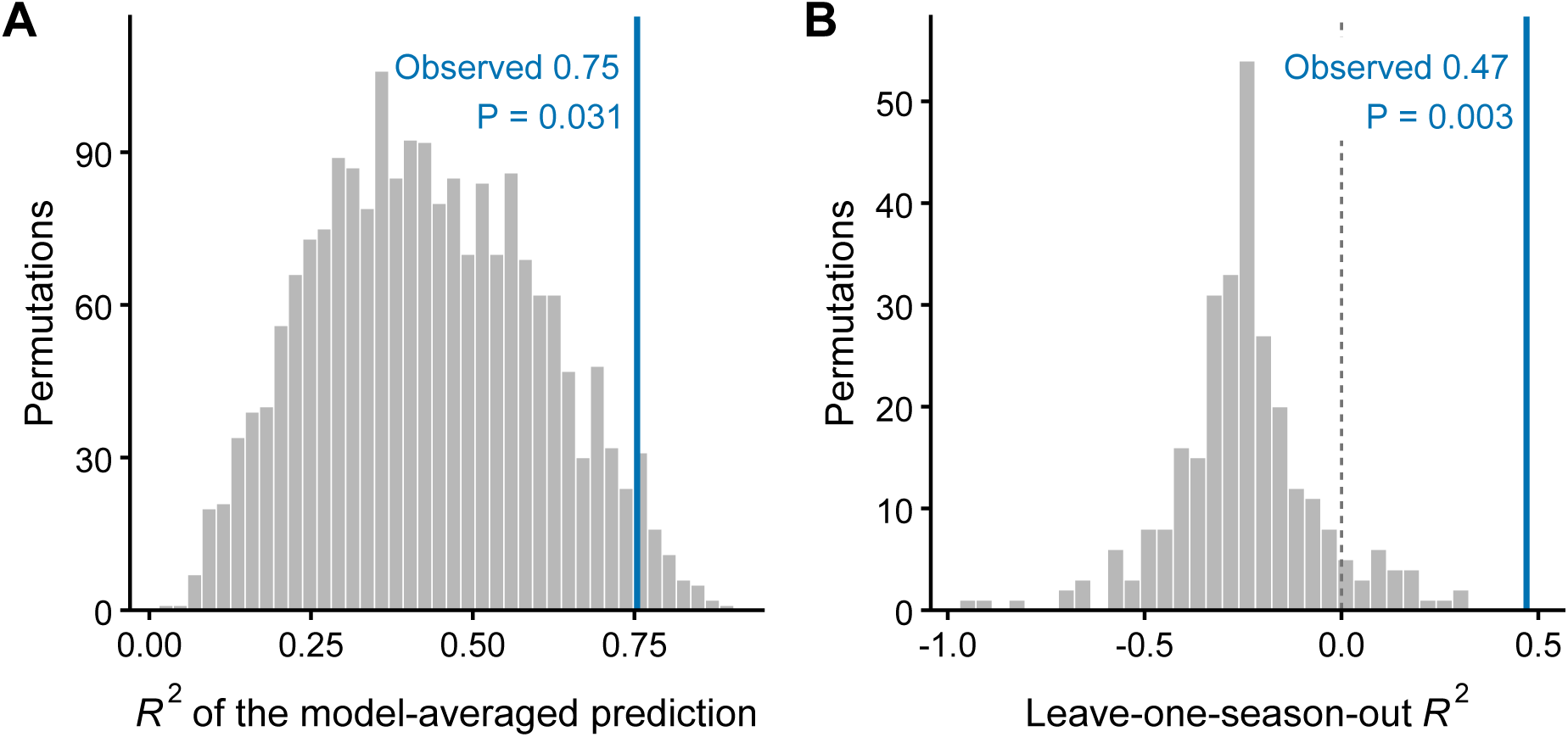
Permutation calibration of the inferential and predictive fits. Grey histograms show the values each procedure reached after VE was randomly reassigned among the 16 seasons; the vertical line marks the observed value. (**A**) *R*^2^ of the AICc-weighted model-averaged prediction, with the whole 377specification procedure repeated in each of 2,000 reassignments (*P* = 0.031). (**B**) Leave-one-season-out *R*^2^ of the predictive model, with all 16 folds refitted in each of 500 reassignments, each moving a VE estimate together with its sampling variance (*P* = 0.003; no reassignment reached the observed value). The dashed line marks zero, below which most held-out *R*^2^ values fall under the null. All *R*^2^ values are on the log OR scale.

**Figure S10:**
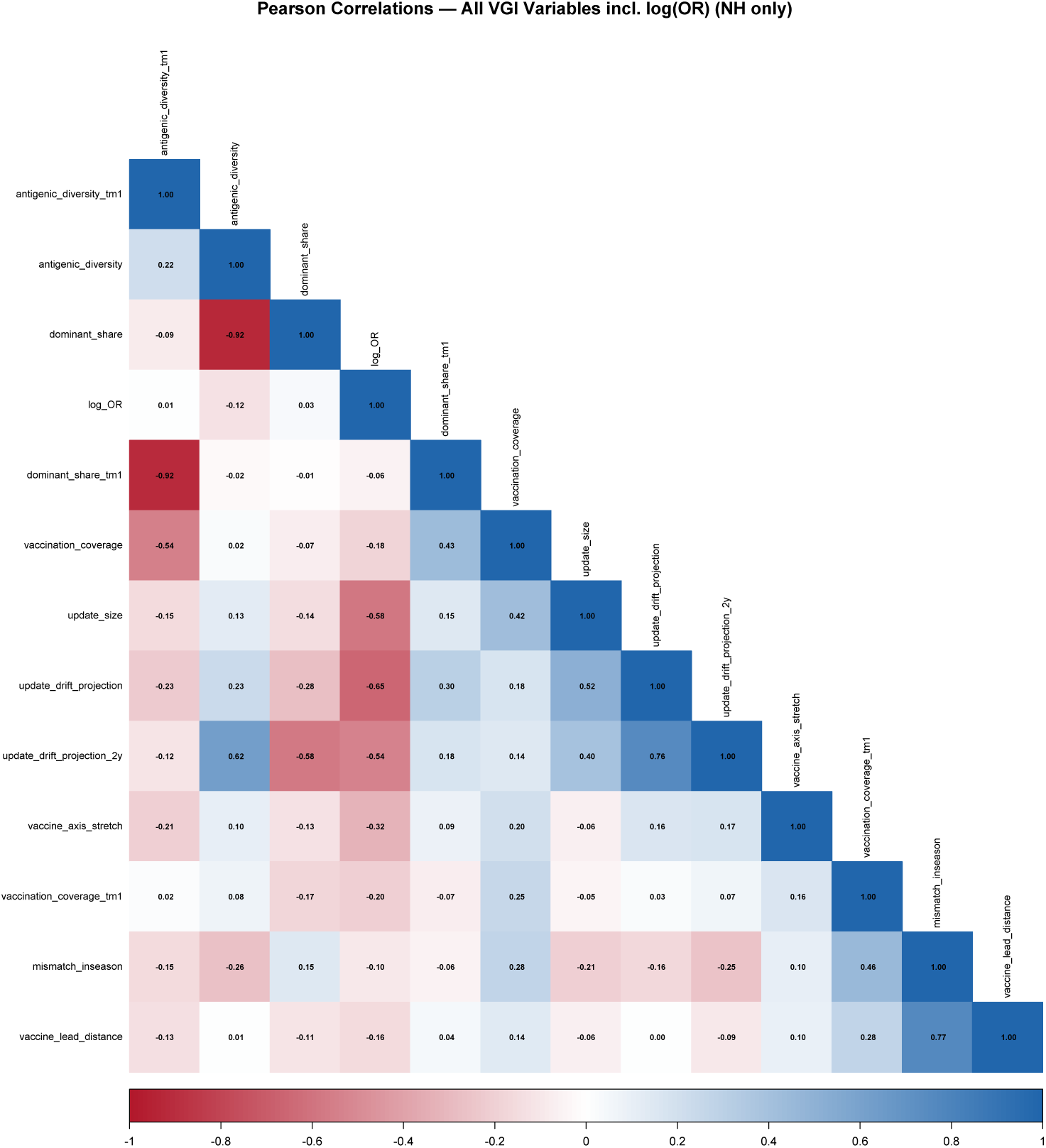
Correlation structure among antigenic predictor variables. Pearson correlation matrix for the predictor variables across seasons with complete data. Strong correlations (*r* > 0.7)—for example, between dominant_share_tm1 and antigenic diversity tm1 (*r* = 0.92)—are why no single specification can separate these variables at *N* = 16, and why the inferential estimates are averaged across specifications while the predictive model shrinks coefficients rather than selecting among them.

**Figure S11:**
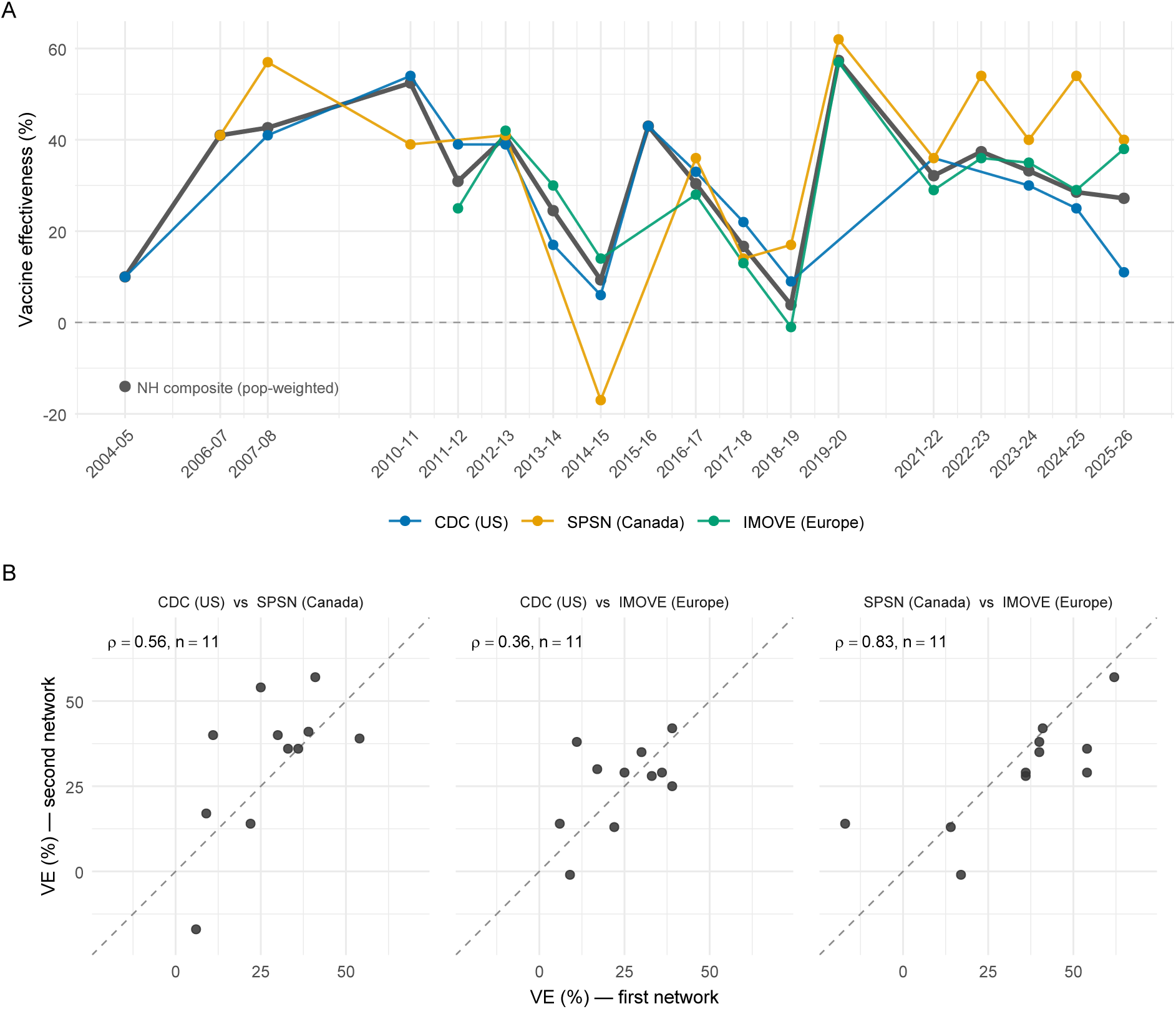
The three surveillance networks give concordant season-level A(H3N2) vaccine effectiveness. (**A**) H3N2-specific VE by season for the CDC (US), SPSN (Canada), and I-MOVE/VEBIS (Europe) networks, with the population-weighted NH composite (grey) used as the modeled outcome. (**B**) Pairwise agreement of network estimates against the identity line, annotated with Spearman *ρ* and the number of jointly reporting seasons. Between-season variance exceeds between-network variance ≈17-fold, indicating that VE tracks the shared circulating antigenic composition rather than the region of measurement.

**Figure S12:**
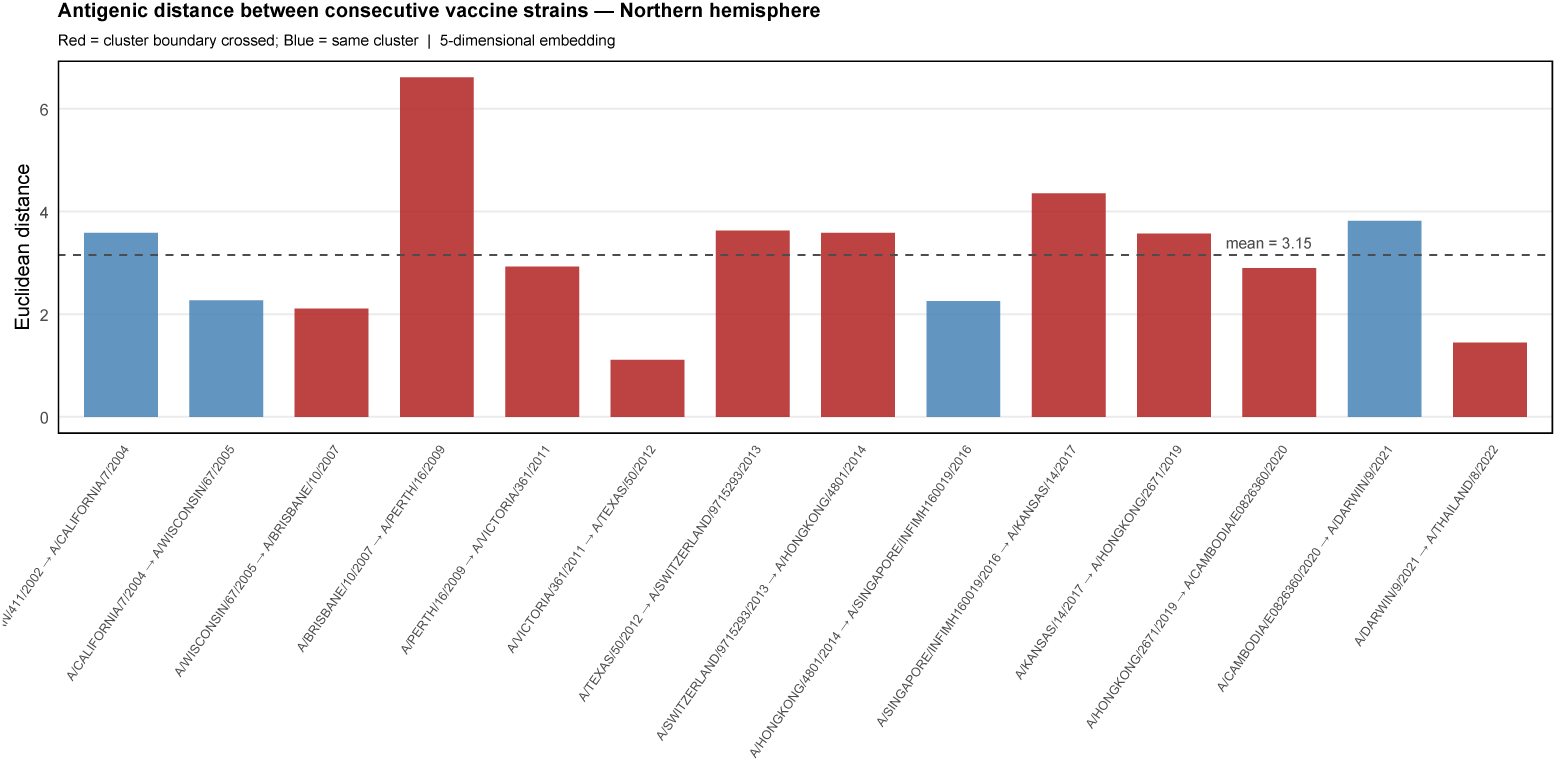
Antigenic distance between consecutive WHO-recommended H3N2 vaccine strains (NH). Euclidean distance in the five-dimensional latent antigenic map between each NH vaccine strain and its successor, from the 2004–2005 to the 2024–2025 season. Bars are colored by whether the update crossed an antigenic-cluster boundary (red) or remained within a cluster (blue); the dashed line marks the mean (3.15 AU). Consecutive updates are small (median 3.25 AU, interquartile range 2.26–3.62 AU; full range 1.11–6.61 AU across 14 updates, the largest being A/Brisbane/10/2007 to A/Perth/16/2009), so the vaccine tracks ongoing drift in short steps and the vaccine-to-population antigenic distance remains compressed.

**Figure S13:**
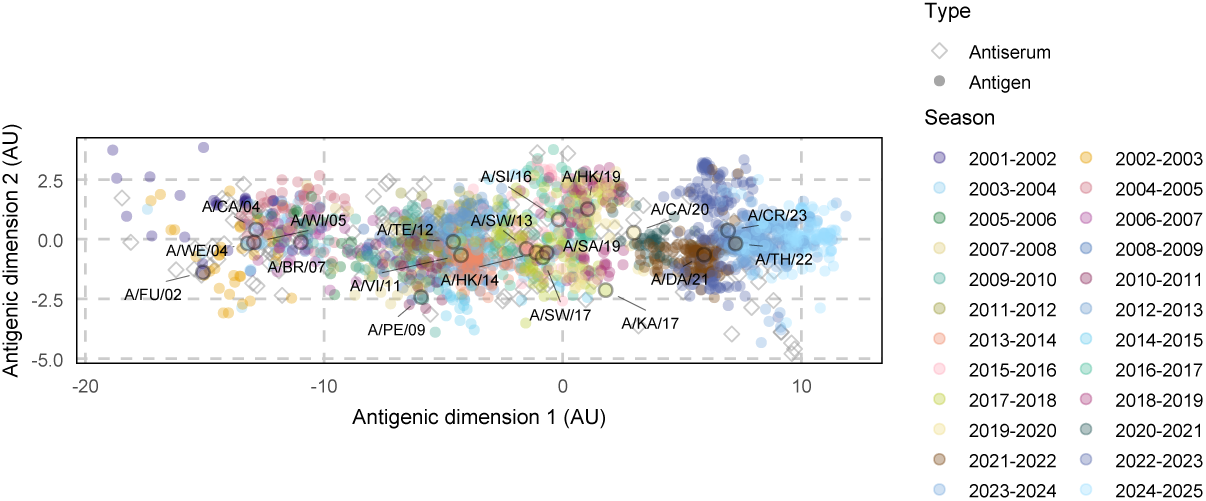
Directional antigenic trajectory of H3N2, 2002–2025. Posterior-mean latent positions (first two map dimensions) colored by season; filled symbols, viruses (antigens); open symbols, reference antisera. WHO vaccine strains are labeled. Strains progress along a dominant antigenic axis from the 2002 reference to the 2024–2025 clusters, illustrating the largely one-dimensional, canalized advance summarized in Fig. 1.

**Figure S14:**
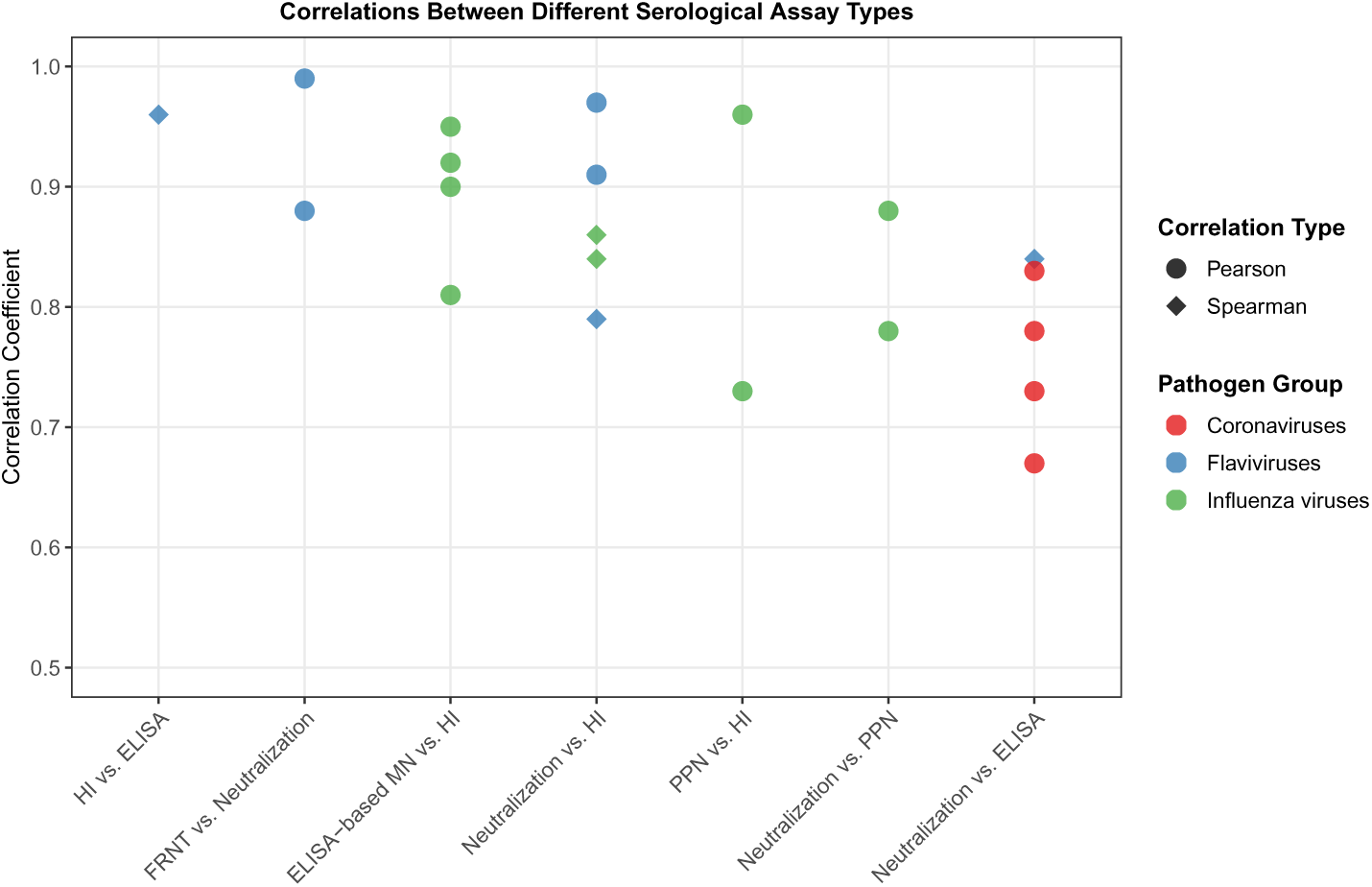
Cross-assay correlation coefficients between serological assay types. Each point represents a published correlation coefficient between two assay types from a single study. Colors indicate pathogen group (influenza viruses, flaviviruses, coronaviruses); shapes distinguish Pearson and Spearman correlation types. For influenza, neutralization versus HI correlations range from *ρ* = 0.84–0.86 (*96, 97*); ELISA-based microneutralization versus HI from *r* = 0.81–0.95 (*92*); pseudoparticle neutralization versus HI and MN from *r* = 0.73–0.96 (*98, 99*). Non-influenza studies (dengue (*101*), JEV (*102, 103*), TBE (*104*), SARS-CoV2 (*105*)) confirm that strong cross-assay convergence is a general property across viral families.

**Figure S15:**
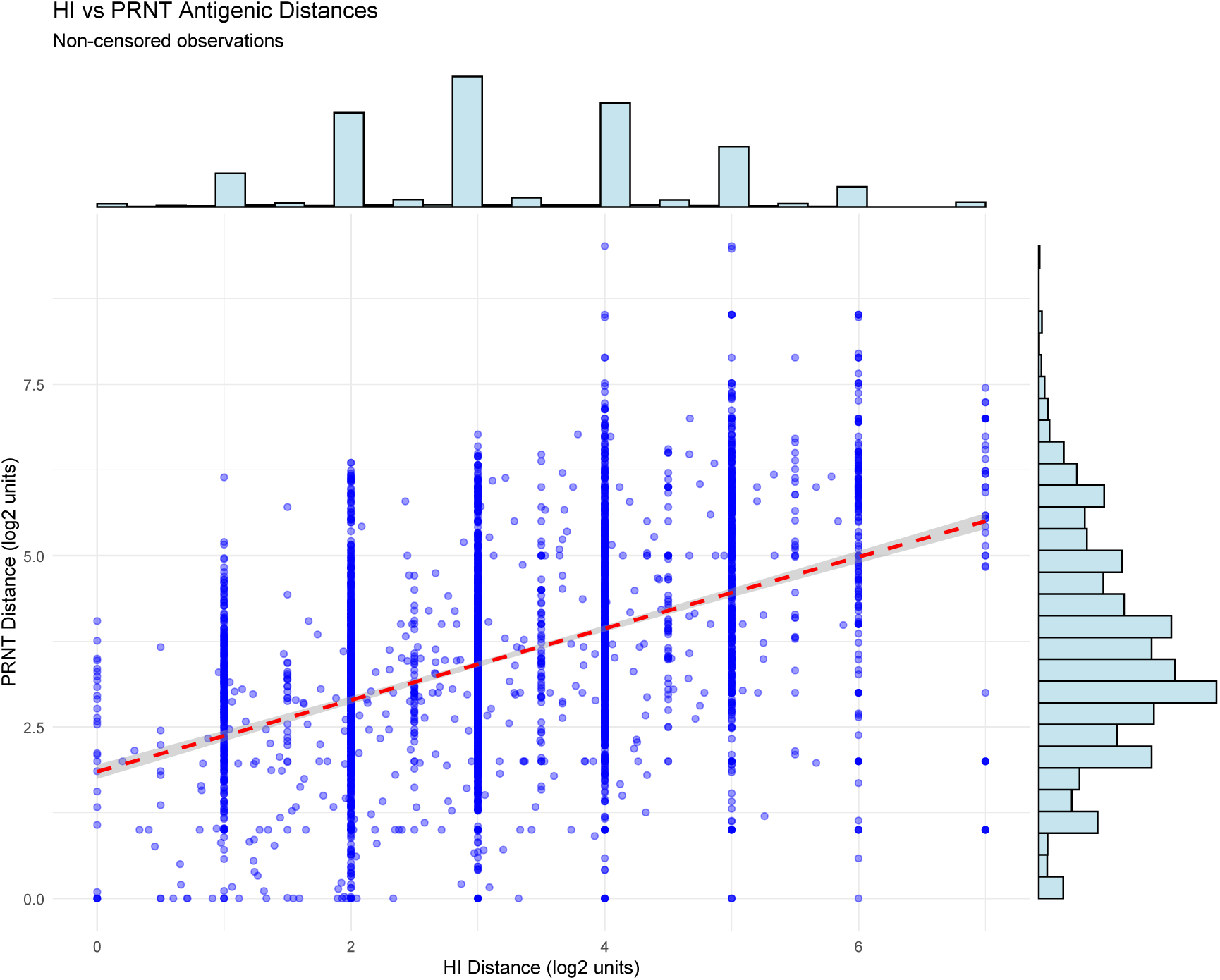
PRNT antigenic distances grow more slowly than HI but carry a positive offset. Non-censored PRNT versus HI antigenic distances (log_2_ units) for virus–serum pairs measured by both assays, with marginal histograms; the dashed red line is an ordinary least-squares fit. PRNT distance increases with HI distance but along a shallower-than-identity slope and from a positive intercept, so PRNT reads higher than HI at small distances yet compresses larger ones. The four-observation-process latent-distance model corrects for the measurement error that attenuates this raw slope and estimates the underlying scaling at *a*_2_ = 0.675 (90% credible interval 0.654, 0.697) with a positive PRNT offset *b*_2_ = 4.08, quantifying the same pattern shown here.

**Figure S16:**
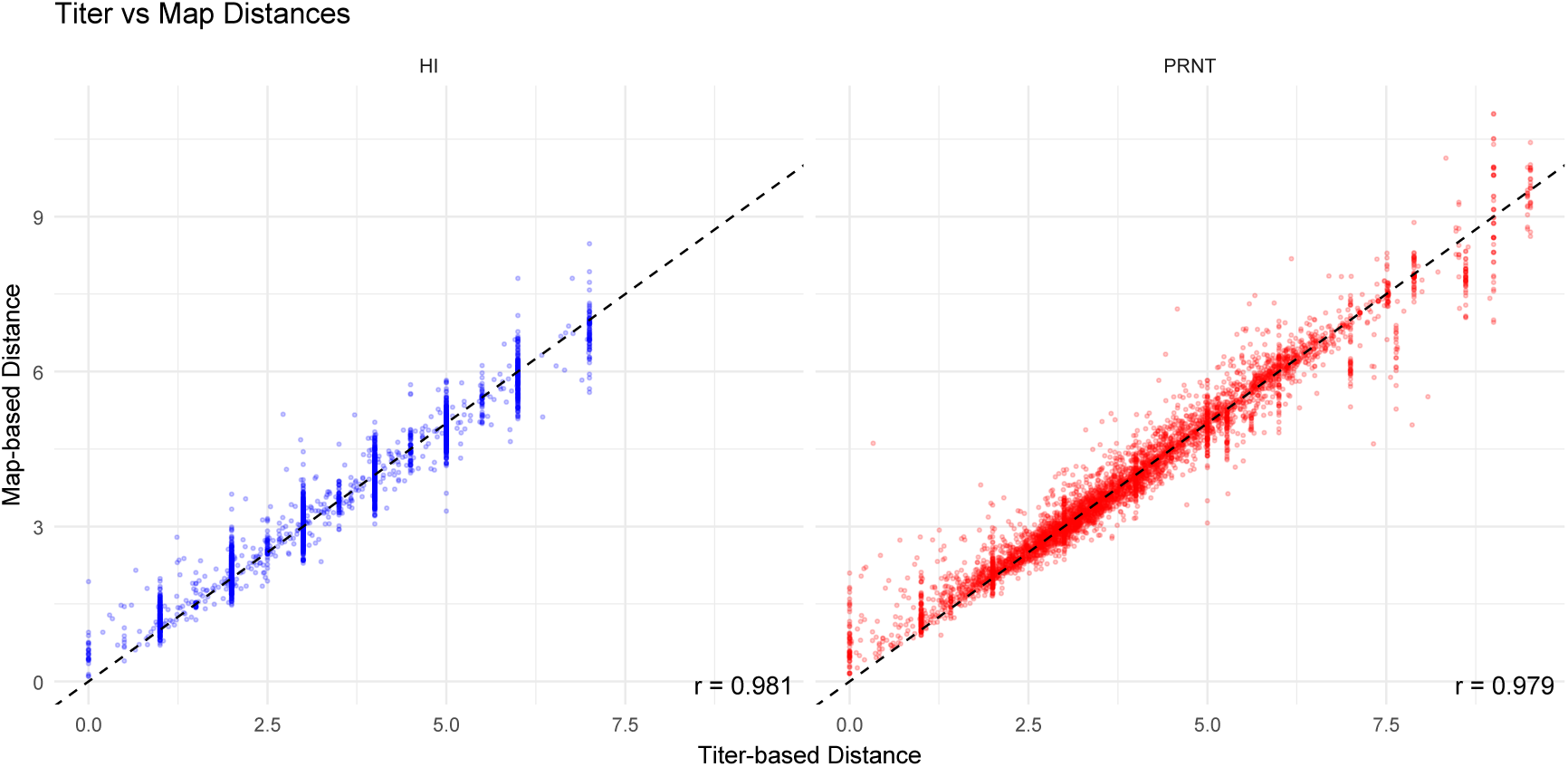
Map distances resolve antigenic differences finer than the discrete titer grid. Topolow map-based antigenic distance versus raw titer-based distance for HI (left) and PRNT (right), in log_2_ AU, for virus–serum pairs measured by each assay; the dashed line is the identity. Although the two are strongly correlated (Pearson *r* = 0.981 for HI, *r* = 0.979 for PRNT), titers take discrete two-fold values—the vertical bands—whereas map distances vary continuously within each band, because the map estimates each pairwise distance jointly from the entire titer network rather than from the single titration. This within-band spread is the cross-strain information the map contributes beyond the paired titer, motivating the inclusion of map distances as separate observations in the four-observation-process latent-distance model.

**Figure S17:**
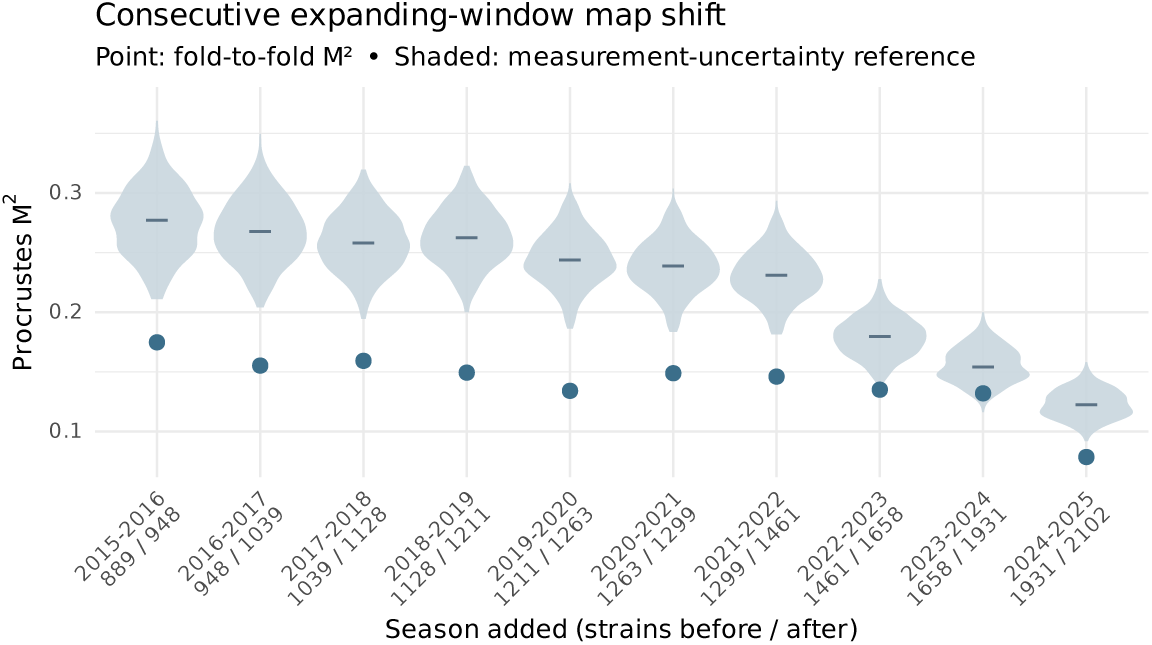
Antigenic map stability under prospective refitting. Each point is the Procrustes *M*^2^ between two consecutive expanding-window latent maps, the second including one more season of titers: the residual sum of squares after optimal rotation, scaling, and translation over the strains present in both maps; lower values indicate greater geometric consistency. The shaded violin at each season is the distribution of *M*^2^ between 300 pairs of latent maps from the measurement-uncertainty ensemble (one map per posterior draw of the latent antigenic distances, so pairs differ only through antigenic measurement uncertainty), restricted to the same strains; the horizontal dash marks its median. The observed *M*^2^ lies below the reference median in every season and below its 2.5th percentile in nine of ten. *x*-axis labels give the number of distinct strain names in the earlier and later map.

**Figure S18:**
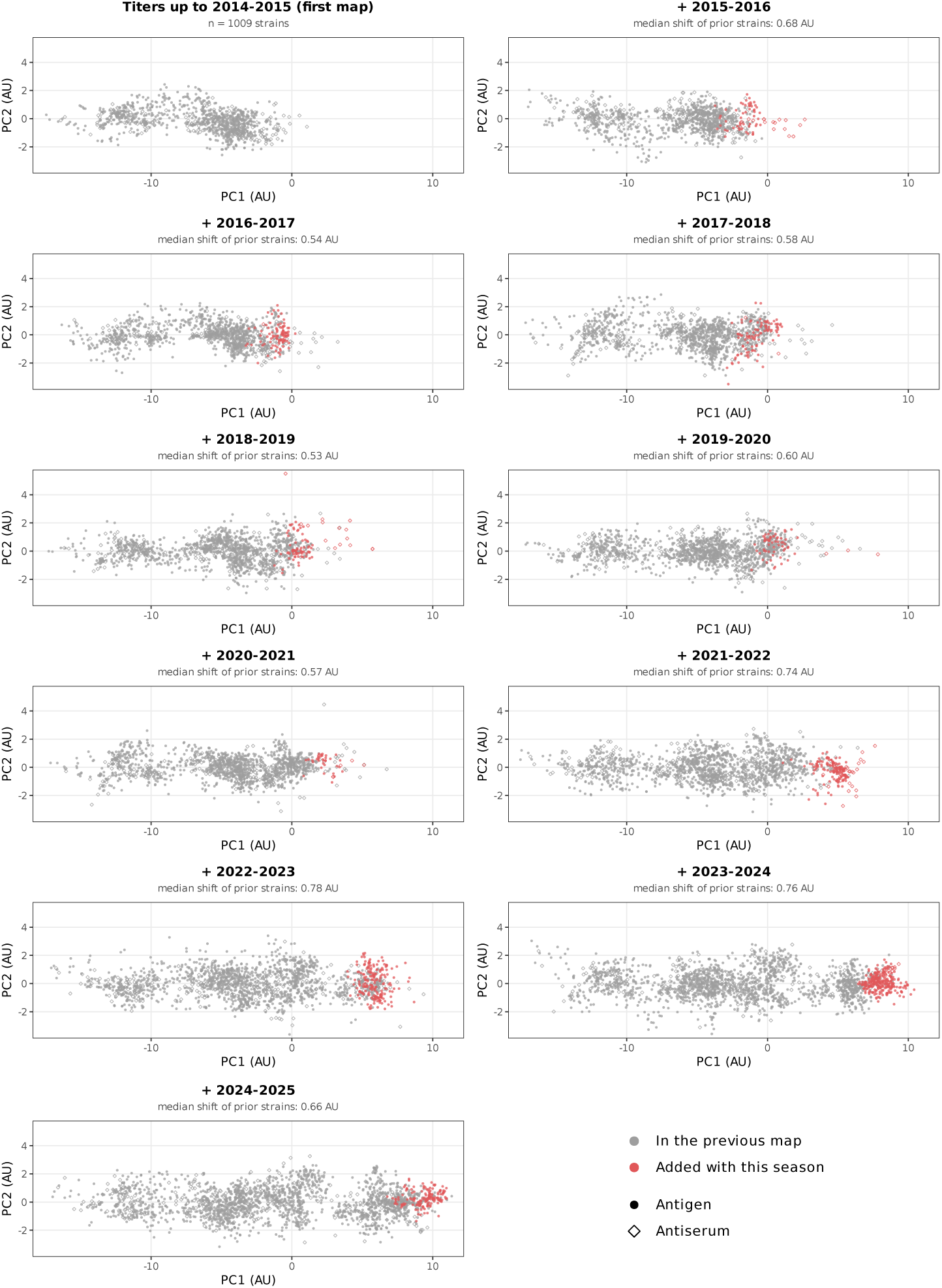
Antigenic-map progression across the expanding-window sequence. Each panel shows the latent antigenic map refit on titers up to the indicated season, drawn in one common frame: every map was aligned (rotation, reflection, translation, and one scale factor) to the full-data map and projected onto the full-data map’s first two principal components (94.7% of its variance), with identical axes in every panel. Grey, strains present in the previous map; red, strains added with that season; circles, antigens; diamonds, antisera. Panel subtitles give the median shift of the prior strains from the previous panel, in antigenic units (AU); per-season values are in table S15.

**Figure S19:**
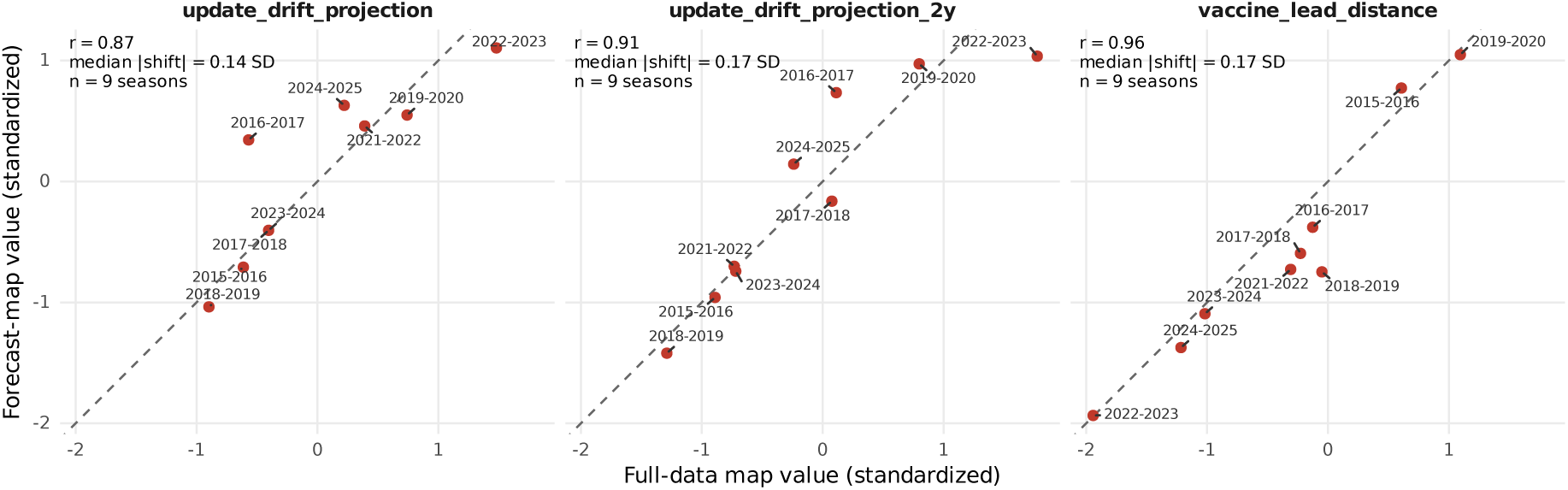
Predictor values at the prospective forecasts against their full-data values. Each point is one of the nine prospectively scored target seasons (2015–2016 to 2024–2025, excluding 2020–2021, which has no VE estimate). The *y*-axis is the predictor value used for that forecast, computed on the expanding-window map built from titers up to the preceding season; the *x*-axis is the value for the same season on the full-data map. Both are standardized with that predictor’s full-data mean and standard deviation (SD) across seasons.

**Figure S20:**
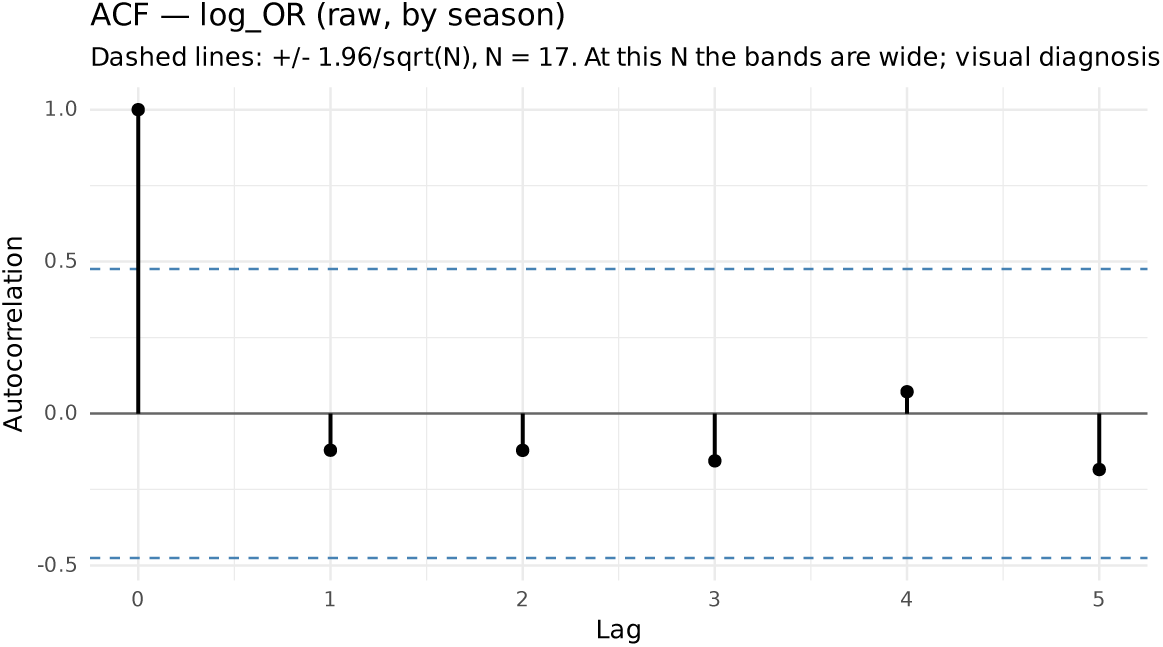
Autocorrelation function of seasonal. log OR . Sample autocorrelations at lags 1–3 with approximate confidence bounds. No lag shows significant autocorrelation, consistent with the formal tests reported in the text and supporting the static ordinary-least-squares specification.

**Figure S21:**
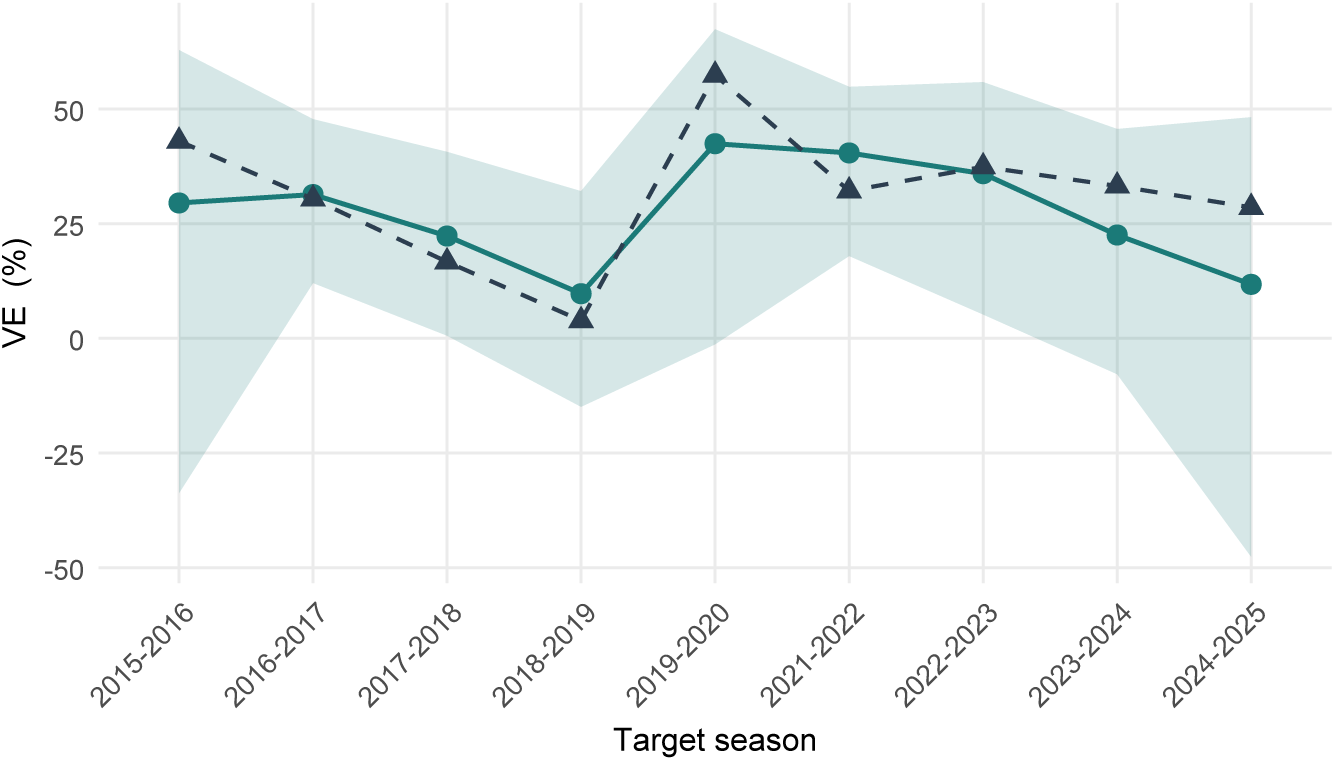
Prospective vaccine-effectiveness predictions, 2015–2016 to 2024–2025. Observed VE (black triangles, dashed) and expanding-window posterior mean predictions (teal), with the shaded blue band the 95% posterior predictive interval returned by the model fitted to each fold. Interval width varies by season with the training size and that season’s measurement error. The antigenic map and all features were refit per season using only prior data.

**Figure S22:**
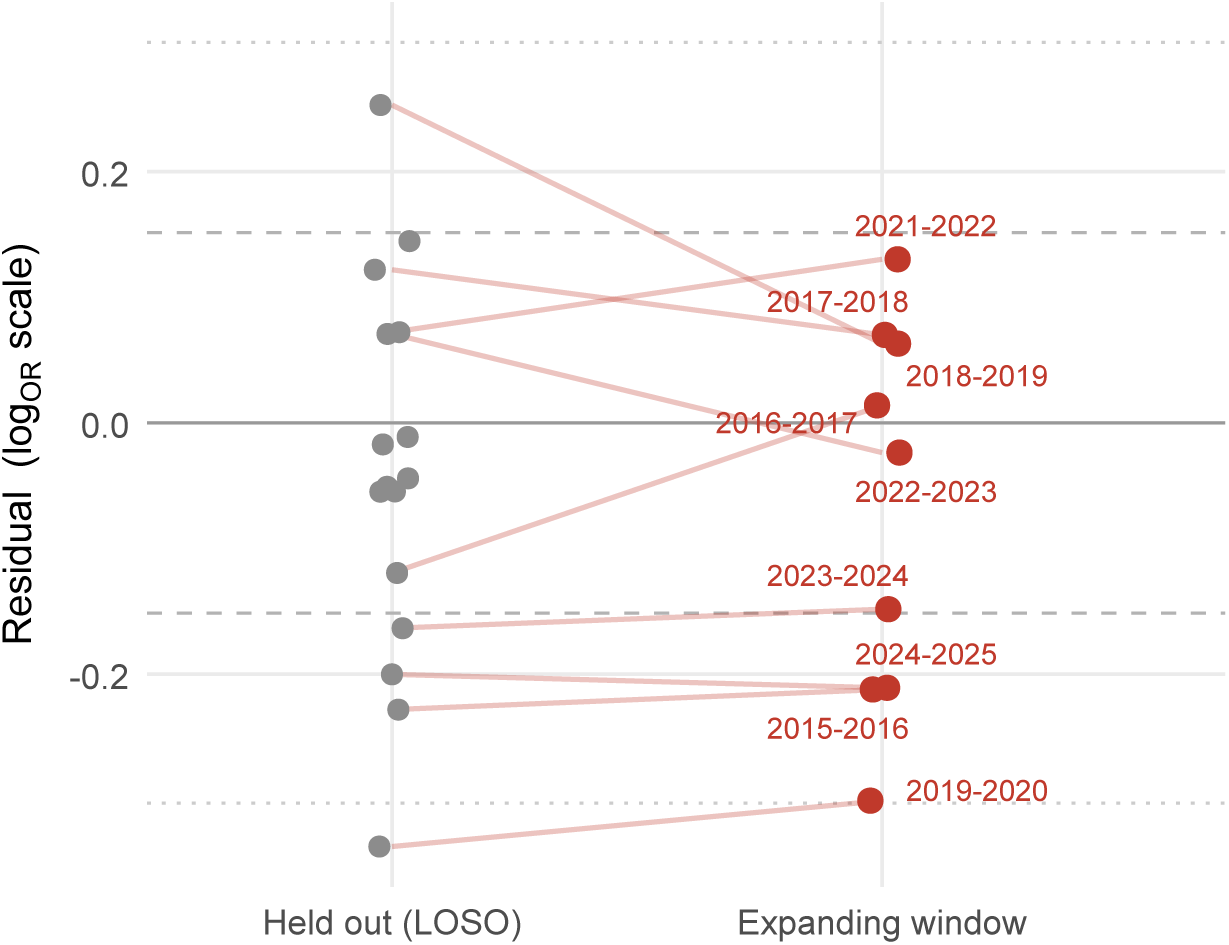
Prospective forecast errors lie within the held-out error distribution. Held-out residuals (grey, all seasons) and the nine expanding-window prospective residuals (red, labeled by target season), on the log OR scale. Dashed and dotted lines mark 1 and 2 standard deviations of the held-out residuals. All prospective residuals fall within the spread of the held-out residuals (nine of nine within the held-out range; table S16), so refitting the antigenic map and all features from prior data only modestly inflates forecast error relative to the held-out benchmark that anchors the predictive claim.

**Table S1:** Inferential model: model-averaged associations with. log OR . *N* = 16 seasons. Every specification of at most three predictors drawn from the 13 pre-specified candidates was fitted (377 in total, each with ≥ 12 residual degrees of freedom), and each variable’s coefficient was averaged over the 79 specifications containing it with AICc weights renormalized within that set (Eq. S9). No specification is selected, so no estimate is conditional on a selection event. *β̂* is on the log OR scale per unit of the variable; *β̄*^∗^ is standardized by the partial standard deviation (Eq. S11). “CI” is the percentile interval over 1,000 bootstrap resamples of the entire procedure; bias-corrected accelerated intervals are given for comparison. *P* is the two-sided bootstrap *P* value from those same resamples (Eq. S12), the achieved significance level of the interval beside it. “Sign” is the percentage of the 79 containing specifications sharing the reported sign, a robustness diagnostic rather than a measure of significance. “Sig.” is the percentage of containing specifications in which the variable is individually significant at *p* < 0.05. Δ*R*^2^ is the order-averaged incremental *R*^2^ (table S11).

| Variable | $\hat{\beta}$ | $\bar{\beta}^*$ | 95% CI | $P$ | Sign | Sig. |
| --- | --- | --- | --- | --- | --- | --- |
| *Vaccine-update orientation* |  |  |  |  |  |  |
| Update–drift projection (1-yr) | −0.085 | −0.72 | [−0.94, −0.13] | 0.014 | 100 | 75 |
| Update–drift projection (2-yr) | −0.042 | −0.38 | [−0.96, 0.10] | 0.102 | 100 | 58 |
| Vaccine-update size | −0.038 | −0.36 | [−0.95, 0.35] | 0.306 | 99 | 68 |
| *Vaccine–virus match* |  |  |  |  |  |  |
| Mismatch, in-season | −0.068 | −0.35 | [−0.76, 0.28] | 0.238 | 92 | 13 |
| Vaccine lead distance | −0.125 | −0.54 | [−0.87, 0.09] | 0.084 | 100 | 27 |
| *Vaccination coverage* |  |  |  |  |  |  |
| Prior-season coverage (%) | −0.014 | −0.25 | [−0.76, 0.20] | 0.218 | 100 | 3 |
| In-season coverage (%) | +0.005 | † | — | — | 90 | 0 |
| *Viral population structure* |  |  |  |  |  |  |
| Antigenic diversity, in-season | +0.133 | † | — | — | 58 | 0 |
| Dominant-cluster share (tm1) | +0.195 | +0.14 | [−0.42, 0.70] | 0.519 | 66 | 0 |
| Dominant-cluster share, in-season | −0.060 | † | — | — | 70 | 0 |
| Antigenic diversity (tm1) | −0.040 | −0.06 | [−0.60, 0.56] | 0.737 | 63 | 0 |
| *Drift geometry* |  |  |  |  |  |  |
| Vaccine-axis stretch | −0.067 | −0.21 | [−0.56, 0.30] | 0.440 | 100 | 0 |
| *Control* |  |  |  |  |  |  |
| $D_{2019-2020}^{\S}$ | −0.302 | −0.37 | [−0.73, −0.10] | 0.010 | 100 | 51 |
| Mean $R^2$ across the 377 specifications, AICc-weighted = 0.680 (range 0.000–0.757) | | | | | | |
| $R^2$ of the AICc-weighted model-averaged prediction = 0.754 | | | | | | |

*Note:* † The across-specification distribution of this estimate is not unimodal—its weighted mean and median disagree in sign, or fewer than 60% of containing specifications share the reported sign—so no averaged effect is reported, no interval or $P$ value for its sign is meaningful, and the distribution is shown instead (Fig. S7); the raw coefficient is retained for reference. § An indicator for a single season is estimable only in the resamples containing that season, so this row’s interval and $P$ value rest on 612 of the 1,000 resamples and are not on the same footing as the others. The bootstrap distributions are right-skewed. Under BCa the intervals for the pandemic-season control, [−0.64, −0.06], and for the 1-yr update–drift projection, [−1.02, −0.37], exclude zero, whereas those for in-season mismatch, [−0.76, 0.30], and for the 2-yr update–drift projection, [−0.85, 0.25], do not. We report the percentile interval throughout; the two methods agree for every candidate<sup>S7</sup> whether the interval excludes zero.

**Table S2:** Predictive model: Bayesian ridge regression coefficients. Dependent variable: log OR . *N* = 16 seasons. Three pre-season predictors; no pandemic dummy and no variable-selection screening. Predictors are standardized, so each coefficient is the change in log OR per one standard deviation of that predictor. Because log OR and VE move in opposite directions, a negative coefficient corresponds to higher effectiveness. The fit carries measurement error on both the outcome and the antigenic predictors (Methods, *Predictive model: Bayesian ridge regression*); the update–drift projection’s 95% credible interval excludes zero, while those of vaccine lead distance and prior-season coverage do not.

| Variable | Posterior mean | 95% CrI | Pr(sign) |
| --- | --- | --- | --- |
| (Intercept) | −0.385 | [−0.461, −0.313] | — |
| Update–drift projection | −0.139 | [−0.219, −0.051] | 0.997 |
| Vaccine lead distance | −0.065 | [−0.140, +0.012] | 0.952 |
| Coverage (tm1) | −0.053 | [−0.122, +0.020] | 0.933 |

Held-out $R^2 = 0.47$ (measurement-noise ceiling 0.483), RMSE = 0.152
CRPS = 0.088 (historical mean 0.128)
Prospective (expanding-window) $R^2 = 0.455$ ( $n = 9$ seasons; ceiling 0.533)
$\tau = 0.109$ , $\sigma_u = 0.065$ [0.004, 0.155]
*Note:* Coefficients are shrunk toward zero by the prior and index predictive contribution rather than inferential effect sizes. CrI: credible interval. Pr(sign) is the posterior probability that the coefficient carries the sign of its mean—a statement about the direction of the association, not a hypothesis test. $\tau$ is the estimated shrinkage scale and $\sigma_u$ the season-to-season variability unexplained by geometry, both on the $\log(\text{OR})$ scale. Held-out $R^2$ is the leave-one-season-out value (sixteen exact refits); the prospective $R^2$ additionally holds out the antigenic map (expanding window, 2015–2016 to 2024–2025; $n = 9$ ). The measurement-noise ceiling is the largest observed-scale $R^2$ attainable given the sampling error of the published VE estimates (Methods, *Reported measures of predictive accuracy*).

**Table S3:** Antigenic measurement error of the predictive model’s predictors. For each predictor, the noise SD is the mean across-draw standard deviation over *D* = 400 antigenic configurations drawn from the latent-distance posterior and passed through the production feature chain. The between-season SD is the predictor’s spread across every mapped season for which it is defined (23, 21 and 20 seasons for coverage, vaccine lead distance, and the update–drift projection). The noise-to-signal ratio divides the former by the latter. The between-season SD is not the same quantity as *s_x_* in Table S10, which is taken over the 16 analysis seasons. Values are on each predictor’s raw scale. Vaccination coverage is not antigenic and is measured exactly, so its noise SD is zero. Candidate strains within a season are scored with a season-centred SD: because they share the prior-season centroid, the vaccine anchor, and the drift axis, part of their positional error cancels in the candidate contrast.

| Predictor | Noise SD | Between-season SD | Noise/signal |
| --- | --- | --- | --- |
| Update–drift projection | 0.408 | 1.529 | 0.267 |
| Vaccine lead distance | 0.243 | 1.049 | 0.231 |
| Vaccination coverage (tm1) | 0.000 | 3.312 | 0.000 |

**Table S4:** Prior sensitivity of the predictive model. Posterior summaries under the reported prior (*τ* and *σ_u_* scales of 0.10) and their range across twelve prior combinations (*τ* scale 0.05, 0.10, 0.20, 0.40 *σ_u_* scale 0.05, 0.10, 0.20). Coefficients are posterior means in log OR per predictor standard deviation. All fits converged (*R̂* < 1.01, no divergent transitions).

| Variable | Reported prior | Range across grid |
| --- | --- | --- |
| Coverage (tm1) | −0.053 | [−0.056, −0.048] |
| Vaccine lead distance | −0.065 | [−0.075, −0.053] |
| Update–drift projection (1-yr) | −0.139 | [−0.154, −0.115] |
| $\tau$ (posterior mean) | 0.109 | [0.075, 0.173] |
| $\sigma_u$ (posterior median) | 0.061 | [0.039, 0.080] |

*Note:* Every coefficient retained its sign in all twelve settings, and no posterior mean moved by more than 0.6 of its posterior standard deviation under the reported prior. The 95% credible interval of the update–drift projection excluded zero in all twelve settings. Re-running the substitution analysis under a tight prior ( $\tau$ and $\sigma_u$ scales of 0.05) and a vague prior ( $\tau$ scale 0.40, $\sigma_u$ scale 0.20) left the model’s candidate unchanged in all 16 of the 16 seasons under both, and gave a pooled mean gain of 10.0 pp and 10.9 pp, against 10.4 pp under the reported prior.

**Table S5:**
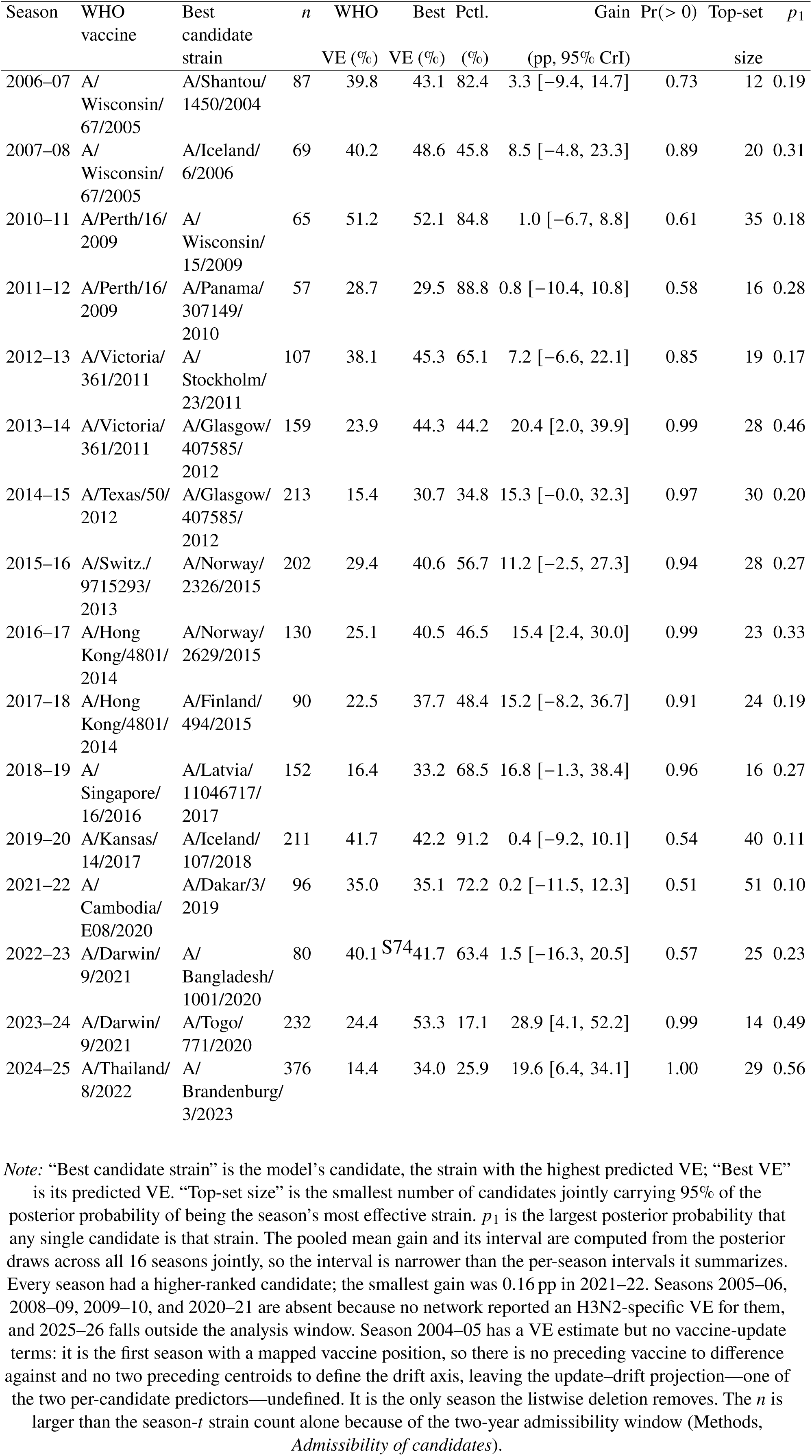
Scenario analysis of substitute candidates: WHO vaccine percentile and optimal strains. For each season *t*, the model was refitted on *N* − 1 seasons, then VE was predicted for all serologically characterized candidate strains assayed within the two years ending 15 January of the year of the vaccine composition meeting (Methods, *Admissibility of candidates*). Rank percentile is the posterior mean proportion of candidates with lower predicted VE than the WHO vaccine; lower percentiles indicate a lower relative ranking of the WHO vaccine. *n*: number of candidate strains evaluated, including the deployed vaccine. Gain: posterior mean difference in predicted VE between the model’s candidate and the WHO vaccine, with its 95% credible interval and the posterior probability that it is positive.

**Table S6:** Detection-anchored vaccine response: model versus WHO. For each of the twelve antigenic clusters in NH circulation, “First dominant” is the earliest season the cluster was the season’s dominant cluster, and is the reference the two lag columns are measured from; “First >10%” is the earliest season the cluster exceeded 10% of that season’s characterized NH strains, the presence criterion used throughout this paper, reported as context for when the cluster became established; “Model flags” is the earliest season the model’s candidate (max predicted VE among *t* 1/*t* 2 strains) belonged to that cluster; “WHO adopts” is the earliest season the recommended vaccine strain belonged to it. “Model lag” and “WHO lag” are seasons from the first dominant season to flag/adoption, *negative when the cluster was nominated before it dominated*, which is what strain selection is trying to achieve. “Response gap” = WHO lag model lag, positive when the model flagged the cluster earlier than the WHO adoption and negative when later. Entries are “—” when a system never chose the cluster; KA17 is marked “never” because it never dominated NH circulation, so it has no anchor and no lags even though the WHO adopted it for 2019–2020. Dominance, rather than first detection, is the anchor because a cluster being detectable does not by itself call for a vaccine update; because both lags share the anchor, the response gap is unaffected by this choice. Descriptive; conditional on the twelve-cluster resolution.

| Cluster | First<br>>10% | First<br>dominant | Model<br>flags | WHO<br>adopts | Model<br>lag | WHO<br>lag | Response<br>gap |
| --- | --- | --- | --- | --- | --- | --- | --- |
| FU02 | 2001–02 | 2004–05 | — | 2004–05 | — | 0 | — |
| BR07 | 2003–04 | 2005–06 | 2006–07 | 2008–09 | 1 | 3 | 2 |
| PE09 | 2006–07 | 2007–08 | 2010–11 | 2010–11 | 3 | 3 | 0 |
| TE12 | 2011–12 | 2011–12 | 2015–16 | 2014–15 | 4 | 3 | –1 |
| SW13 | 2010–11 | 2013–14 | 2011–12 | 2012–13 | –2 | –1 | 1 |
| HK19 | 2014–15 | 2015–16 | 2013–14 | 2020–21 | –2 | 5 | 7 |
| SW17 | 2014–15 | 2017–18 | 2018–19 | 2016–17 | 1 | –1 | –2 |
| DA21 | 2020–21 | 2020–21 | 2022–23 | 2021–22 | 2 | 1 | –1 |
| CA22 | 2022–23 | 2022–23 | — | — | — | — | — |
| CR23 | 2022–23 | 2023–24 | — | 2024–25 | — | 1 | — |
| EN24 | 2023–24 | 2024–25 | 2024–25 | — | 0 | — | — |
| KA17 | 2018–19 | never | — | 2019–20 | — | — | — |

Model lag: –2 to 4 seasons (8 clusters nominated); anticipated dominance in 2
WHO lag: –1 to 5 seasons (9 adopted clusters that dominated); anticipated in 2
Response gap, clusters both chose ( $n = 7$ ): mean 0.857 seasons (range –2 to 7)
Median lag on those 7: model 1 season, WHO 3 seasons
Concurrent dominant-cluster match: model 5/16, WHO 5/16 seasons
Model early-flag hit rate: 8/8 clusters became dominant or adopted
*Note:* Negative lags are anticipatory, not speculative: no cluster was nominated by either system before it had appeared in NH circulation at all. The model anticipated dominance for two clusters (SW13, two seasons ahead; HK19, two seasons ahead) and the WHO vaccine for two (SW13 and SW17, one season each). All eight clusters the model flagged went on to dominate circulation or enter the vaccine, so none was a false positive. FU02, KA17, CA22, and CR23 carry no model flag because the model never

**Table S7:** Posterior predictive recovery of the four data sources by the four-observation-process latent-distance model. For each observation process we compared observed values against the model’s posterior predictive means (all quantities on the log_2_ scale; the two titer observation processes evaluated on non-censored pairs). RMSE is the root-mean-square error of the posterior predictive mean; posterior predictive *R*^2^ is the squared Pearson correlation between observed and predicted values; 95% coverage is the fraction of observations falling within the 95% posterior predictive interval. Posterior predictive coverage of the 95% intervals ranged from 94.9% to 99.5% across the four observation processes, with the PRNT map observation process falling just below the nominal 95%. Point recovery was near-perfect for the two HI-based observation processes (*R*^2^ = 0.99) but lower for the noisier PRNT-based observation processes (*R*^2^ = 0.23–0.34), reflecting the intrinsically higher PRNT measurement noise.

| Data source | RMSE ($\log_2$) | Posterior predictive $R^2$ | 95% coverage |
| --- | --- | --- | --- |
| HI titer | 0.147 | 0.989 | 99.4% |
| PRNT titer | 1.416 | 0.231 | 96.9% |
| HI map distance | 0.120 | 0.993 | 99.5% |
| PRNT map distance | 1.429 | 0.338 | 94.9% |

**Table S8:** Convergence and predictive comparison of the five candidate unification models. ΔELPD is the leave-one-out expected log pointwise predictive density (*64*) relative to the best model (M LAT 4OBS, the four-observation-process latent-distance model); more negative is worse. Models with *R*^^^ > 1.01 failed to converge and were excluded from the ELPD ranking. M COR produced 2,996 divergent transitions; the four-observation-process model produced none. The leave-one-out approximation is unreliable for these latent-variable models—each virus–serum pair carries its own latent position, so the estimated effective number of parameters is large (*p*_loo_ 4,715, roughly one per four of the 20,748 pointwise terms) and 8.9% of pointwise Pareto-*k* values exceeded 0.7. The ELPD ranking is therefore corroborative only, with posterior predictive checks and prior sensitivity analysis serving as the primary validation.

| Model | Structure | $\hat{R}_{\text{max}}$ | ΔELPD ± SE |
| --- | --- | --- | --- |
| M_LAT_4OBS | Latent, four observation equations | 1.003 | 0 (reference) |
| M_INDP | No shared latent variable | 1.007 | $-19,603 \pm 169$ |
| M_LAT_2OBS | Latent, titers only | 2.25 | excluded* |
| M_COR | No shared latent, 4-variate normal | 4.47 | excluded* |
| M_LAT_CORR | Latent, within-assay correlated errors | 1.074 | excluded* |

\*Excluded from ELPD ranking owing to non-convergence ( $\hat{R} > 1.01$ ).

**Table S9:** Influenza A(H3N2)-specific vaccine effectiveness (VE) estimates by surveillance network and season, with population-weighted NH composite. Estimates are adjusted and outpatient/primary care unless noted; all are all-ages except the 2024–25 and 2025–26 CDC values, which are adult (aged ≥18 years) U.S. Flu VE outpatient estimates. — indicates data not available from that network. Seasons with no estimate from any network (2005–06, 2008–09, 2009–10, 2020–21) are omitted. The published source of each network–season estimate is listed in *Collection and processing of influenza A(H3N2) vaccine effectiveness estimates*, *Per-season data sources*.

| Season | CDC (US) |  | SPSN (Canada) |  | I-MOVE (Europe) |  | NH |  |
| --- | --- | --- | --- | --- | --- | --- | --- | --- |
|  | VE% | 95% CI | VE% | 95% CI | VE% | 95% CI | VE% | 95% CI |
| 2004–05<sup>a</sup> | 10 | (–36, 40) | — | — | — | — | 10.0 | (–36.0, 40.0) |
| 2006–07 | — | — | 41 | (6, 63) | — | — | 41.0 | (6.0, 63.0) |
| 2007–08<sup>b</sup> | 41 | (24, 53) | 57 | (32, 73) | — | — | 42.7 | (24.8, 55.1) |
| 2010–11 | 54 | (42, 64) | 39 | (0, 63) | — | — | 52.5 | (37.7, 63.9) |
| 2011–12 | 39 | (23, 52) | — | — | 25 | (–6, 47) | 30.9 | (6.3, 49.1) |
| 2012–13 | 39 | (29, 47) | 41 | (17, 59) | 42 | (15, 61) | 40.7 | (20.7, 55.3) |
| 2013–14 | 17 | (–45, 53) | — | — | 30 | (–34, 63) | 24.5 | (–38.7, 58.8) |
| 2014–15 | 6 | (–5, 17) | –17 | (–50, 9) | 14 | (–6, 31) | 9.3 | (–7.6, 24.3) |
| 2015–16 | 43 | (4, 66) | — | — | — | — | 43.0 | (4.0, 66.0) |
| 2016–17 | 33 | (23, 41) | 36 | (18, 50) | 28 | (17, 38) | 30.4 | (19.5, 39.8) |
| 2017–18 | 22 | (12, 31) | 14 | (–8, 31) | 13 | (–15, 34) | 16.7 | (–3.8, 32.7) |
| 2018–19 | 9 | (–4, 20) | 17 | (–13, 39) | –1 | (–24, 18) | 3.9 | (–15.4, 19.8) |
| 2019–20 | — | — | 62 | (37, 77) | 57 | (27, 75) | 57.4 | (27.8, 75.2) |
| 2021–22 | 36 | (20, 49) | 36 | (–38, 71) | 29 | (12, 42) | 32.1 | (12.9, 46.2) |
| 2022–23<sup>c</sup> | — | — | 54 | (38, 66) | 36 | (25, 45) | 37.4 | (26.0, 46.6) |
| 2023–24 | 30 | (8, 47) | 40 | (5, 61) | 35 | (20, 48) | 33.2 | (14.5, 48.2) |
| 2024–25<sup>c</sup> | 25 | (–6, 48) | 54 | (29, 70) | 29 | (–22, 60) | 28.5 | (–13.2, 55.6) |
| 2025–26<sup>c</sup> | 11 | (–18, 33) | 40 | (28, 49) | 38 | (29, 46) | 27.2 | (10.0, 40.9) |

<sup>a</sup> Overall influenza VE used because >90% of circulating viruses were A(H3N2) (120).
<sup>b</sup> VE against influenza A used because all characterized A viruses were H3N2 (133).
<sup>c</sup> All 2024–25 and 2025–26 estimates are interim (mid-season) rather than end-of-season. In addition, the CDC column uses adult (aged ≥18 years) outpatient U.S. Flu VE Network H3N2-specific estimates; for these two seasons the network reported H3N2 outpatient VE only by age stratum (children and adults), with no pooled all-ages estimate (141, 142).

**Table S10:** Standardized coefficients of the inferential and predictive models. Standardized coefficients for all 13 candidate variables, with *s_y_* log OR = 0.215. Signs indicate the direction of association with log OR (negative = higher VE). *β̂* Infer. is the raw model-averaged coefficient and *β̄*^∗^ (Infer.) its standardization by the partial standard deviation (Eq. S11); within that column, a larger *β̄*^∗^ is a stronger partial association. *β*_std_ (Pred.) is a separate estimate rather than a rescaling of *β̂* Infer.: it is the coefficient of the Bayesian ridge fit, standardized by the marginal standard deviation (Eq. S25). Those coefficients are shrunk toward zero by the ridge penalty, so their relative magnitudes are not a variable-importance ordering and are not read as one; the variance a variable accounts for is given by the incremental *R*^2^ of Table 1. *s_x_*, the predictor’s marginal standard deviation, is shown only for the three predictive predictors.

| Variable | $s_x$ | $\hat{\beta}_{\text{Infer.}}$ | $\bar{\beta}^*$ (Infer.) | $\beta_{\text{std}}$ (Pred.) |
| --- | --- | --- | --- | --- |
| Update–drift projection (1-yr) | 1.74 | −0.085 | −0.72 | −0.62 |
| Vaccine lead distance | 0.87 | −0.125 | −0.54 | −0.31 |
| Update–drift projection (2-yr) | — | −0.042 | −0.38 | — |
| $D_{2019-2020}$ | — | −0.302 | −0.37 | — |
| Vaccine-update size | — | −0.038 | −0.36 | — |
| Mismatch (in-season) | — | −0.068 | −0.35 | — |
| Coverage (tm1) | 3.71 | −0.014 | −0.25 | −0.25 |
| Vaccine-axis stretch | — | −0.067 | −0.21 | — |
| Dominant-cluster share (tm1) | — | +0.195 | +0.14 | — |
| Antigenic diversity (tm1) | — | −0.040 | −0.06 | — |
| Antigenic diversity (in-season) | — | +0.133 | † | — |
| Coverage (in-season) | — | +0.005 | † | — |
| Dominant-cluster share (in-season) | — | −0.060 | † | — |

*Note:* In the $s_x$ and $\beta_{\text{std}}$ (Pred.) columns, “—” marks a variable that is not one of the three predictive predictors. † marks the three in-season structural variables whose across-specification estimate is not unimodal, so no averaged $\bar{\beta}^*$ is reported (the raw coefficient is retained for reference; distribution in Fig. S7). The two standardized columns are not on the same scale: the inferential column uses the partial standard deviation (Eq. S11), required because its estimate is averaged across specifications that condition on different covariates, and the predictive column the marginal standard deviation (Eq. S25) of one fixed specification. Both divide by $s_y$ , so the predictive coefficients appear here in outcome standard-deviation units; Table S2 reports the same coefficients in $\log(\text{OR})$ units.

**Table S11:** Explained variance by mechanistic category. Within each of the 377 specifications, the incremental *R*^2^ of every predictor was averaged over all orders of entry, which apportions that specification’s *R*^2^ among its predictors exactly; the shares were then averaged across specifications with the AICc weights and summed within category. The category shares therefore sum to the AICc-weighted mean *R*^2^ of the space (0.680) by construction, which is the arithmetic check on the decomposition rather than a result. Because the full averaging-over-orderings decomposition would require the saturated 13-predictor model, which has 2 residual degrees of freedom at *N* = 16, this is the decomposition restricted to the three-predictor lattice (Methods, *Inferential model: multimodel estimation*).

| Mechanistic category | Variables | $\Delta R^2$ | Share of total |
| --- | --- | --- | --- |
| Vaccine-update orientation | 3 | 0.428 | 63.0% |
| Vaccine–virus match | 2 | 0.176 | 25.8% |
| Pandemic-season control | 1 | 0.059 | 8.6% |
| Vaccination coverage | 2 | 0.010 | 1.5% |
| Drift geometry | 1 | 0.005 | 0.8% |
| Viral population structure | 4 | 0.002 | 0.3% |
| Total | 13 | 0.680 | 100% |

*Note:* Per-variable shares, largest first: update–drift projection 1-yr 0.383, vaccine lead distance 0.159, $D_{2019-2020}$ 0.059, vaccine-update size 0.023, update–drift projection 2-yr 0.022, in-season mismatch 0.016, prior-season coverage 0.008, vaccine-axis stretch 0.005; the five remaining variables together account for less than 0.004.

**Table S12:** How controlling for one mechanistic category changes another variable’s estimate. For each variable, the specifications containing it were split by whether they also include an indicator of the stated category, and the two halves were averaged separately, with the weights renormalized within each half. The difference is the part of the variable’s estimate attributable to confounding with that category. A single fitted model cannot report this quantity: it gives one estimate, conditional on the covariates it includes. Entries are *β̄*^∗^. Rows list every contrast with Δ 0.10 for the eight variables whose averaged estimate is reportable and at least 0.15 in magnitude. Of the other five variables, three have no reportable averaged estimate and two are indistinguishable from zero, so their shifts are not interpretable

| Variable | Controlling for | Without | With | $\Delta$ |
| --- | --- | --- | --- | --- |
| Update–drift projection (2-yr) | Viral population structure | −0.30 | −0.65 | −0.34 |
| Vaccine-update size | Drift geometry | −0.31 | −0.64 | −0.33 |
| Vaccine-update size | Vaccination coverage | −0.30 | −0.59 | −0.29 |
| Vaccine-update size | Viral population structure | −0.32 | −0.60 | −0.28 |
| Vaccine-update size | Vaccine–virus match | −0.50 | −0.25 | +0.25 |
| Prior-season coverage | Vaccine–virus match | −0.35 | −0.15 | +0.20 |
| $D_{2019-2020}$ | Vaccine-update orientation | −0.55 | −0.37 | +0.18 |
| $D_{2019-2020}$ | Vaccine–virus match | −0.48 | −0.31 | +0.17 |
| Vaccine lead distance | Vaccine-update orientation | −0.38 | −0.54 | −0.16 |
| Vaccine-axis stretch | Viral population structure | −0.20 | −0.36 | −0.15 |
| Prior-season coverage | Pandemic-season control | −0.21 | −0.35 | −0.14 |
| Prior-season coverage | Viral population structure | −0.25 | −0.39 | −0.14 |
| $D_{2019-2020}$ | Vaccination coverage | −0.35 | −0.49 | −0.14 |
| Vaccine lead distance | Pandemic-season control | −0.56 | −0.42 | +0.14 |
| Mismatch, in-season | Vaccine-update orientation | −0.21 | −0.35 | −0.14 |
| $D_{2019-2020}$ | Viral population structure | −0.36 | −0.50 | −0.14 |
| Update–drift projection (1-yr) | Vaccine–virus match | −0.60 | −0.74 | −0.14 |
| Update–drift projection (2-yr) | Vaccination coverage | −0.37 | −0.49 | −0.12 |
| Mismatch, in-season | Viral population structure | −0.34 | −0.45 | −0.12 |
| Vaccine-axis stretch | Vaccine–virus match | −0.29 | −0.18 | +0.10 |

*Note:* A negative $\Delta$ on a negative estimate means the association strengthens once the stated category is controlled for. The pattern is consistent with mutual but asymmetric masking between the two leading categories. These are subset averages without intervals; a $\Delta$ of this size is within the range each estimate spans across other conditioning choices. Controlling for vaccine-update orientation strengthens both vaccine–virus match estimates: by roughly two-thirds for in-season mismatch (−0.21 to −0.35) and by two-fifths for vaccine lead distance (−0.38 to −0.54). Controlling for vaccine–virus match strengthens the one-season projection by a comparable amount, 0.14 (−0.60 to −0.74), but leaves the two-season projection

**Table S13:** Held-out VE predictions (Bayesian ridge regression). For each season, the model was refitted on the other *N* − 1 seasons. Posterior mean predicted VE (converted from log OR) with 95% credible intervals on the expected value. Observed VE from population-weighted TND estimates. The wider 95% predictive intervals, which also carry the season-level and measurement variance and are the intervals assessed for calibration, contained the observed value in all 16 seasons.

| Season | Obs. VE (%) | Pred. VE (%) | 95% CrI | Residual |
| --- | --- | --- | --- | --- |
| 2006–2007 | 41.0 | 40.0 | [30.6, 48.0] | +1.0 |
| 2007–2008 | 42.7 | 39.7 | [19.2, 55.7] | +3.0 |
| 2010–2011 | 52.5 | 49.8 | [30.3, 65.1] | +2.7 |
| 2011–2012 | 30.9 | 27.8 | [21.2, 34.4] | +3.1 |
| 2012–2013 | 40.7 | 37.4 | [27.2, 47.2] | +3.3 |
| 2013–2014 | 24.5 | 23.6 | [16.0, 31.3] | +0.9 |
| 2014–2015 | 9.3 | 21.5 | [10.2, 32.2] | −12.2 |
| 2015–2016 | 43.0 | 28.4 | [15.8, 40.0] | +14.6 |
| 2016–2017 | 30.4 | 21.6 | [10.0, 31.6] | +8.8 |
| 2017–2018 | 16.7 | 26.2 | [18.6, 34.0] | −9.6 |
| 2018–2019 | 3.9 | 25.4 | [8.7, 38.0] | −21.5 |
| 2019–2020 | 57.4 | 40.3 | [28.0, 52.6] | +17.1 |
| 2021–2022 | 32.1 | 36.9 | [28.0, 45.1] | −4.7 |
| 2022–2023 | 37.4 | 41.7 | [18.7, 58.2] | −4.3 |
| 2023–2024 | 33.2 | 21.4 | [9.7, 31.2] | +11.9 |
| 2024–2025 | 28.5 | 12.7 | [−10.0, 31.4] | +15.8 |

**Table S14:** Per-variable leakage audit. L1 (response leakage): use of *VE t* or a function of it. L2 (covariate-construction leakage): dependence on the joint antigenic map fit with titers from seasons > *t*. “Map-derived” variables inherit L2 from the map; coverage and the pandemic indicator do not. Model(s) marks whether a variable is in the three-predictor predictive model (A1), the 13-variable inferential pool, or both; covers all 13 candidate variables in the pre-specified inferential pool (construct groups + structural controls).

| Variable | Model(s) | L1 | L2 | Mitigation |
| --- | --- | --- | --- | --- |
| update_drift_projection | Both | None | Moderate | Position stability;<br>forecast-value agree-<br>ment |
| vaccine_lead_distance | Both | None | Moderate | Position stability;<br>forecast-value agree-<br>ment |
| update_drift_projection_2y | Inferential | None | Moderate | Position stability;<br>forecast-value agree-<br>ment |
| vaccination_coverage_tm1 | Both | None | None | Not map-derived |
| vaccination_coverage | Inferential | None | None | Not map-derived |
| dominant_share_tm1 | Inferential | None | Moderate | Cluster-label transfer |
| dominant_share | Inferential | None | Moderate | Cluster-label transfer |
| antigenic_diversity_tm1 | Inferential | None | Moderate | Cluster-label transfer |
| antigenic_diversity | Inferential | None | Moderate | Cluster-label transfer |
| vaccine_axis_stretch | Inferential | None | Moderate | Position stability |
| update_size | Inferential | None | Low | Vaccine-to-vaccine dis-<br>tance; minimal map<br>dependence |
| mismatch_inseason | Inferential | None | Moderate | Position stability |
| $D_{2019-2020}$ | Inferential | None | None | Calendar indicator |

**Table S15:**
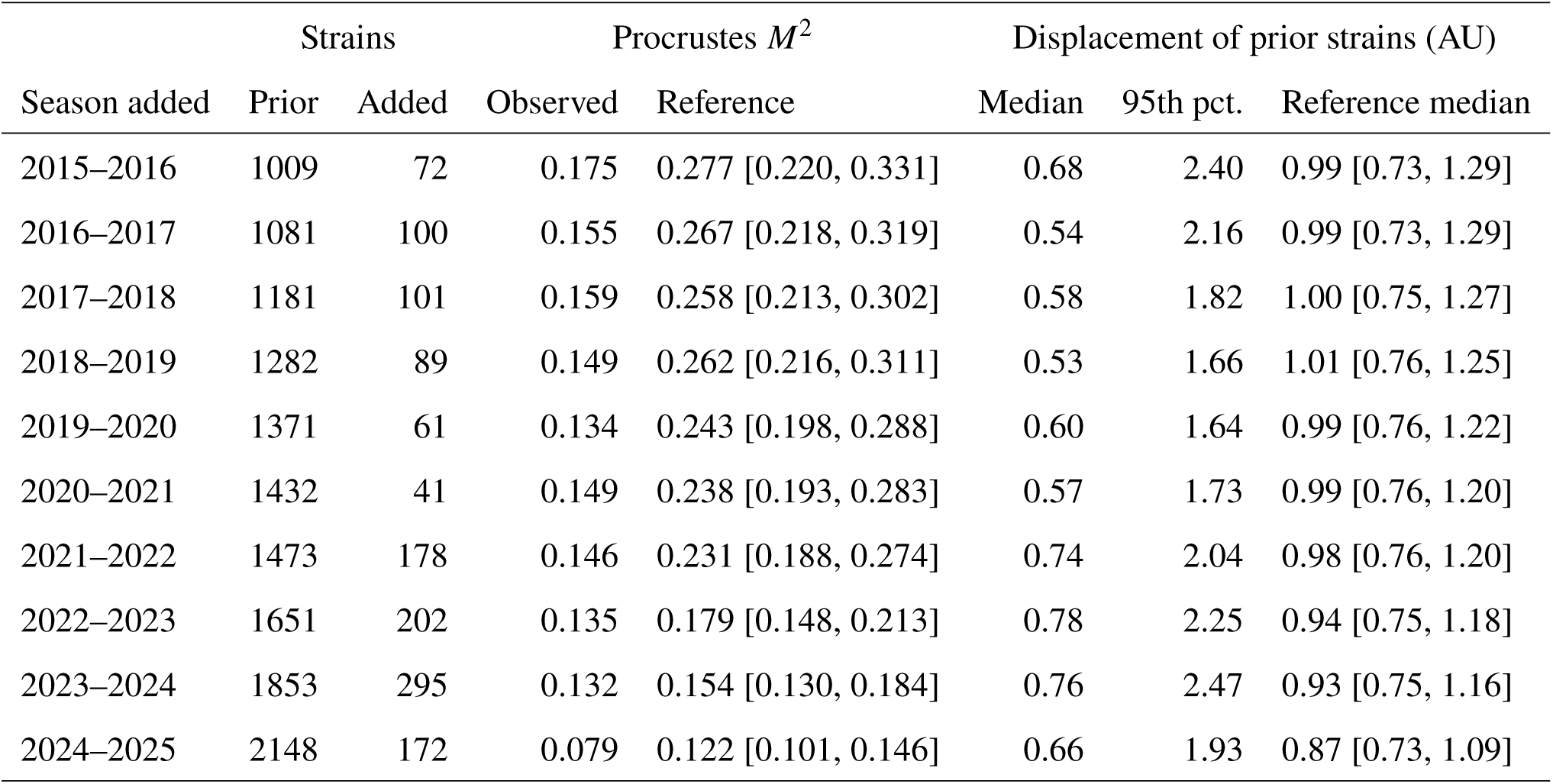
Antigenic-map stability as each season is added. For each season whose titers were added to the expanding-window map, the strains already present in the previous map (prior strains) are compared between the two consecutive maps. *M*^2^ is the symmetric Procrustes residual between the two maps over the prior strains (unitless; 0 means identical). Displacement is the distance in antigenic units (AU) between a prior strain’s positions in the two consecutive maps after both are aligned to the full-data map and projected onto its first two principal components (94.7% of its variance), the frame shown in Fig. S18; the median and 95th percentile are over the prior strains. Each reference column gives the same statistic between pairs of maps from the measurement-uncertainty ensemble—one latent map per posterior draw of the latent antigenic distances, so that pairs differ only through antigenic measurement uncertainty—restricted to the same strains.

| Season added | Strains | | Procrustes $M^2$ | | Displacement of prior strains (AU) | | |
| --- | --- | --- | --- | --- | --- | --- | --- |
|  | Prior | Added | Observed | Reference | Median | 95th pct. | Reference median |
| 2015–2016 | 1009 | 72 | 0.175 | 0.277 [0.220, 0.331] | 0.68 | 2.40 | 0.99 [0.73, 1.29] |
| 2016–2017 | 1081 | 100 | 0.155 | 0.267 [0.218, 0.319] | 0.54 | 2.16 | 0.99 [0.73, 1.29] |
| 2017–2018 | 1181 | 101 | 0.159 | 0.258 [0.213, 0.302] | 0.58 | 1.82 | 1.00 [0.75, 1.27] |
| 2018–2019 | 1282 | 89 | 0.149 | 0.262 [0.216, 0.311] | 0.53 | 1.66 | 1.01 [0.76, 1.25] |
| 2019–2020 | 1371 | 61 | 0.134 | 0.243 [0.198, 0.288] | 0.60 | 1.64 | 0.99 [0.76, 1.22] |
| 2020–2021 | 1432 | 41 | 0.149 | 0.238 [0.193, 0.283] | 0.57 | 1.73 | 0.99 [0.76, 1.20] |
| 2021–2022 | 1473 | 178 | 0.146 | 0.231 [0.188, 0.274] | 0.74 | 2.04 | 0.98 [0.76, 1.20] |
| 2022–2023 | 1651 | 202 | 0.135 | 0.179 [0.148, 0.213] | 0.78 | 2.25 | 0.94 [0.75, 1.18] |
| 2023–2024 | 1853 | 295 | 0.132 | 0.154 [0.130, 0.184] | 0.76 | 2.47 | 0.93 [0.75, 1.16] |
| 2024–2025 | 2148 | 172 | 0.079 | 0.122 [0.101, 0.146] | 0.66 | 1.93 | 0.87 [0.73, 1.09] |

**Table S16:** Calibration of expanding-window prospective errors against held-out errors. Ten target seasons (2015–2016 to 2024–2025), nine with VE estimates; the antigenic map and all features were refit using only data available before each season. Benchmarks are the corresponding quantities when only the outcome is held out and the map is fitted on the full panel. The two CRPS values are each scored against their own reference forecast—fold-training-mean for the prospective column, historical mean for the held-out benchmark—and are not combined into one skill score (Methods).

| Metric | Value | Benchmark |
| --- | --- | --- |
| MAE, log(OR) scale | 0.130 | Held-out 0.122 (ratio 1.07) |
| MAE, VE (pp) | 8.7 | Held-out 8.4 (ratio 1.03) |
| RMSE, log(OR) scale | 0.159 | Held-out 0.152 (ratio 1.05) |
| Out-of-sample $R^2$ | 0.455 (ceiling 0.533) | Held-out 0.47 (ceiling 0.483) |
| CRPS, log(OR) scale | 0.084 (fold-training-mean<br>0.130; difference 0.046) | Held-out 0.088 (historical mean 0.128;<br>difference 0.040) |
| Residuals within held-out range | 9 / 9 | — |
| 95% predictive-interval coverage | 9 / 9 | expected 8.55 |
| Spearman's $\rho$ | 0.767 | — |

*Note:* Intervals are posterior predictive intervals from the model itself, not constructed from the residuals they are assessed against. CRPS is the continuous ranked probability score (lower is better). Each column reports the model's mean CRPS, its own reference forecast, and the paired difference between them on the same seasons: the prospective reference is the mean of each fold's training log(OR), the held-out reference the forecasting distribution of VE in the other training seasons. The two references are not interchangeable, so the differences are not comparable across columns (Methods). Differences are reported rather than fractional reductions because a ratio of two means has a small, noisy denominator at this number of seasons. At nine seasons $R^2$ and Spearman's $\rho$ are unstable, and the CRPS comparison—which uses the whole forecast distribution for every season—is the more reliable summary.

**Table S17:** NH influenza vaccination coverage by region and season, with the population-weighted composite used as the modeled covariate. Values are the percentage of the indicated age group vaccinated against seasonal influenza. United States values are adult (18 years) end-of-season estimates; European values are adult coverage inferred from the Eurostat 65-year rate; Canadian values are general (12 years) CCHS estimates (see *Collection and processing of vaccination coverage data* for sources and the harmonization procedure). The composite is the population-weighted NH rate (vaccination coverage t); its one-season lag (vaccination coverage tm1) is the modeled pre-season predictor, so each season contributes to the regression through the following season’s observation. — indicates no value available for that region and season; the composite re-normalizes weights over the available regions.

| Season | US ($\geq 18$) | EU (adult, inf.) | Canada ($\geq 12$) | NH composite |
| --- | --- | --- | --- | --- |
| 2003–04 | 42.7 | 29.7 | — | 35.2 |
| 2004–05 | 35.5 | 30.6 | 36.0 | 32.8 |
| 2005–06 | 38.3 | 32.0 | — | 34.7 |
| 2006–07 | 39.3 | — | 32.0 | 38.5 |
| 2007–08 | 40.2 | 24.0 | 30.0 | 30.8 |
| 2008–09 | 39.0 | 24.7 | 30.0 | 30.7 |
| 2009–10 | 40.4 | 23.7 | 26.0 | 30.5 |
| 2010–11 | 40.5 | 24.6 | 28.0 | 31.2 |
| 2011–12 | 38.8 | 21.9 | 28.0 | 29.0 |
| 2012–13 | 41.5 | 21.2 | 28.0 | 29.7 |
| 2013–14 | 42.2 | 21.3 | 31.0 | 30.2 |
| 2014–15 | 43.6 | 19.7 | 33.3 | 30.0 |
| 2015–16 | 41.7 | 18.8 | 31.6 | 28.6 |
| 2016–17 | 43.3 | 18.9 | 32.6 | 29.4 |
| 2017–18 | 37.1 | 18.9 | 31.5 | 26.8 |
| 2018–19 | 45.3 | 19.2 | 33.6 | 30.4 |
| 2019–20 | 48.4 | 19.8 | 39.0 | 32.2 |
| 2020–21 | 50.2 | 20.5 | 40.0 | 33.4 |
| 2021–22 | 49.4 | 23.9 | 34.3 | 34.7 |
| 2022–23 | 46.9 | 22.8 | 43.0 | 33.5 |
| 2023–24 | — | 22.2 | — | 22.2 |
| 2024–25 | 41.9 | 22.2 | — | 30.5 |

**Caption for Data S1. Source data for every figure and table.** A compressed archive of 58 comma-separated files holding the numbers plotted in each figure and printed in each table. A file serving several items is included once under the item name where it first appears. The archive also contains data s1 manifest.csv, which lists each display item, its file, any other items that file supports, the row count, and a one-line description of the contents. Notably, fig4 all candidate predictions.csv gives the substitute-candidate scenario predictions for 2,310 candidate strains × seasons. Its columns are the season and candidate name, the posterior mean predicted log(OR) and VE (%) with posterior SD and quantiles, the probabilities of being the season optimum and of exceeding the WHO pick, the per-candidate predictors (vaccine lead distance, update drift projection) with prior-season coverage, and Mahalanobis distances flagging candidates outside the training predictor range. Predictions for the 16 WHO picks are in fig4 who predictions.csv.

## Notes

https://doi.org/10.5281/zenodo.22902309

### Competing Interest Statement

The authors have declared no competing interest.

### Summary of Updates

Title changed from "Geometry of antigenic evolution improves influenza vaccine selection" to "Directional antigenic drift forecasts influenza vaccine effectiveness and guides strain selection". Author list and affiliations unchanged. Abstract rewritten. It now reports the inferential result that geometric features of antigenic space accounted for about three-quarters of the variation in vaccine effectiveness. Antigenic threshold changed throughout from 2 AU to 3 AU, the conventional eight-fold antigenic-distinction criterion. All affected figures, tables, and derived numbers were regenerated at the new threshold, including the candidate-coverage match radius in Figure 5. Section on antigenic evolution across the pandemic gap recomputed on global antigenic years rather than Northern Hemisphere seasons (pre-pandemic mean 1.5 AU per year; 2.8 AU then 2.5 AU across the two pandemic seasons; dispersion minimum 1.9 AU). Statistical support added: permutation calibration of the in-sample fit (P = 0.03), the leave-one-season-out forecast skill (P = 0.003), and the expanding-window skill (P = 0.008), with a new supplementary figure and a new Methods section. Bootstrap resampling is now reported over 1,000 resamples of the whole procedure. Figures revised: Figure 2 gained a Southern Hemisphere panel (the separate supplementary figure was removed); Figure 3 is labeled a schematic; Figure 5 panels were relabeled and given a pooled trend line; Table 1 uses human-readable variable names. Supplementary materials updated: new prior-sensitivity table, new antigenic-map stability table and figures, and renumbering to Figures S1 to S22 and Tables S1 to S17. Data S1 now supplies source data for every figure and table, including per network-season vaccine effectiveness estimates with their individual source citations and the vaccination coverage series. Sourcing and reuse terms are documented, and code was published.

