## Supplementary material for "Directional antigenic drift forecasts influenza vaccine effectiveness and guides strain selection": Data S1: README.txt

Data S1 -- source data for "Antigenic geometry predicts H3N2 influenza vaccine effectiveness"
Omid Arhami and Pejman Rohani

WHAT THIS IS
  The machine-readable numbers behind every figure and table in the paper.
  It is source data, not a reproduction pipeline: analysis code, Stan models
  and the derived antigenic coordinates are in the archive cited in the Data,
  code and materials availability statement.

FILE NAMING
  <display item>_<original pipeline filename>.csv
  e.g. fig4_all_candidate_predictions.csv, tableS3_uq_predictor_noise_summary.csv
  A few legacy filenames embed an EARLIER supplementary table numbering
  (table_S8_*, table_S12_*). The prefix is authoritative; ignore the number
  inside the original filename.

FILES SHARED BY SEVERAL DISPLAY ITEMS
  Copied once, named for the item where they first appear in the paper. The
  'also_supports' column of data_s1_manifest.csv lists every other item a file
  backs, so look there rather than expecting one file per item.

MANIFEST
  data_s1_manifest.csv -- display item, file, other items it supports, row
  count, size, the path it came from in the analysis repository, and a
  one-line description of its contents.

DISPLAY ITEMS WITH NO DATA FILE
  fig3 -- Illustrative geometric schematic; panel coordinates are hard-coded in scripts/figures/fig_variable_construction.R. Its panel F coverage series is the one in tableS17_global_coverage_comparison.csv.
  figS14 -- Published pairwise assay correlations taken from the cited literature; the values are hard-coded in scripts/Assays_corrs.R and are not ours to tabulate as data.
  figS15 -- Plots raw Crick HI and PRNT titers, which are not redistributed; request them from the Francis Crick Institute.
  figS16 -- Plots raw Crick HI and PRNT titers against map distances; same restriction as fig. S15.
  tableS14 -- Per-variable leakage judgments authored by the analysts; there is no underlying data file.

RESTRICTED DATA
  Raw hemagglutination-inhibition and plaque-reduction neutralization titers
  were obtained from the WHO Collaborating Centre at the Francis Crick
  Institute and are not redistributed here. They may be requested directly
  from the Crick. The antigenic coordinates derived from them are included.

Generated by scripts/build_data_s1.R on 2026-09-21.
